# Menarche onset is an inflection point for mental health and brain development

**DOI:** 10.64898/2026.04.13.718198

**Authors:** Carina Heller, Holly Sullivan-Toole, Martin Gell, Sanju Koirala, John McClellan France, Ran Barzilay, Tyler M. Moore, Ka I Ip, Damien A. Fair, Brenden Tervo-Clemmens, Arielle S. Keller, Adriene M. Beltz, Emily G. Jacobs, Bart Larsen

**Author notes:** Corresponding author Carina Heller, Ph.D. Masonic Institute for the Developing Brain, Department of Pediatrics, University of Minnesota, MN, USA.

## Abstract

Menarche is a normative milestone of female puberty, yet its role in adolescent mental health and brain development remains poorly understood. Using longitudinal data from 5,016 females (7 annual visits, ages 10–16 years) in the Adolescent Brain Cognitive Development Study, we found that menarche onset functions as an inflection point for the development of internalizing symptoms and gross brain morphometry. The onset of menarche, largely independent of timing and socio-environmental factors, preceded a significant spike in internalizing symptoms, while altering the rate of ongoing structural brain development. Following menarche onset, individuals with faster declines in gray matter volume and surface area also had heightened internalizing symptoms. These findings suggest that menarche is not only a reproductive milestone but a neuroendocrine driver of adolescent brain and mental health trajectories. This normative and easily identifiable marker could define a critical window for mental health screenings with greater precision than current age-based guidelines.

## Main

Adolescence is a period of profound brain development^1,2^ during which sex-related psychiatric vulnerabilities emerge^3^. Females face a two-fold increased risk for internalizing mental health disorders across the lifespan compared to males^4,5^, implicating female pubertal processes in shaping neurobiologically-underpinned mental health trajectories. Menarche, the onset of menstruation, is a normative milestone in female puberty that marks a fundamental transition across biological, physiological, and psychosocial processes of maturation^6,7^. While earlier menarche timing (age at menarche) is a known predictor of depression risk^8–10^, menarche onset itself has not been investigated as a milestone of brain development and symptom emergence. Here, we investigated the trajectories of internalizing symptoms and gross brain morphometry according to *chronological* and *gynecological* age. While chronological age measures years from birth, gynecological age centers the developmental timeline relative to each individual’s specific menarche onset (centered at zero), allowing us to align individual developmental trajectories to a shared pubertal milestone, regardless of when it chronologically occurred.

We analyzed longitudinal data from a large, heterogeneous sample of *N*=5,655 females from the Adolescent Brain Cognitive Development (ABCD) Study, spanning six years of development (ages 10–16 years). Our analyses integrated self-reported menarche timing with longitudinal measures of internalizing symptoms, structural brain morphometry, and the general exposome (a multivariate index of socio-environmental context^11^). During the study period, 88.7% (*n*=5,016) of participants self-reported menarche (Extended Data Fig. 1a). Consistent with U.S. population-based reports^12–14^, mean menarche timing was 11.8 years (SD=1.2 years) and varied significantly with youths’ exposomes (β=0.277, *P*<0.001) such that lower general exposome scores (reflecting higher environmental adversity and lower socioeconomic status) were associated with earlier menarche timing. Accordingly, all following models adjusted for general exposome and, where appropriate, menarche timing.

First, we examined the development of parent-reported Child Behavior Checklist (CBCL)^15^ internalizing symptoms according to chronological and gynecological age. Analysis of chronological age replicated the established finding^3^ that internalizing symptoms increased with chronological age. Our nonlinear model estimated that internalizing symptoms significantly increased between ages 11.8–15.2 years (*P*_Bonferroni_<0.001; Extended Data Fig. 2, Supplementary Table 1), with maximum acceleration of symptoms (the inflection point) coinciding with the average age of menarche in this sample, at 11.8 years (95% credible interval [CI]=11.1–14.4 years (y), Supplementary Table 2).

Analysis of gynecological age further demonstrated that the increase in internalizing symptoms was tightly aligned with menarche onset: Our model estimated that internalizing symptoms rose significantly from approximately one month post-menarche onset to 3.3 years post-menarche onset, with maximum acceleration at 0.39 years post onset (95% CI: −0.20– 0.95y; **Fig. 1a**, Supplementary Table 1–2). Notably, the estimate was more precise for gynecological age, suggesting menarche onset is a precise inflection point demarcating changes in internalizing symptoms (**Fig. 1b**). To confirm the role of menarche as an inflection point, we replicated this finding using an alternate youth-report measure of internalizing symptoms^16^ (Extended Data Fig. 3, Supplementary Table 3–4), in split-half samples (Extended Data Fig. 4, Supplementary Table 5–6), when accounting for familial dependencies (Supplementary Table 7–8), and when estimating gynecological age based on parent-report (Extended Data Fig. 5, Supplementary Table 9–10).

**Fig. 1.**
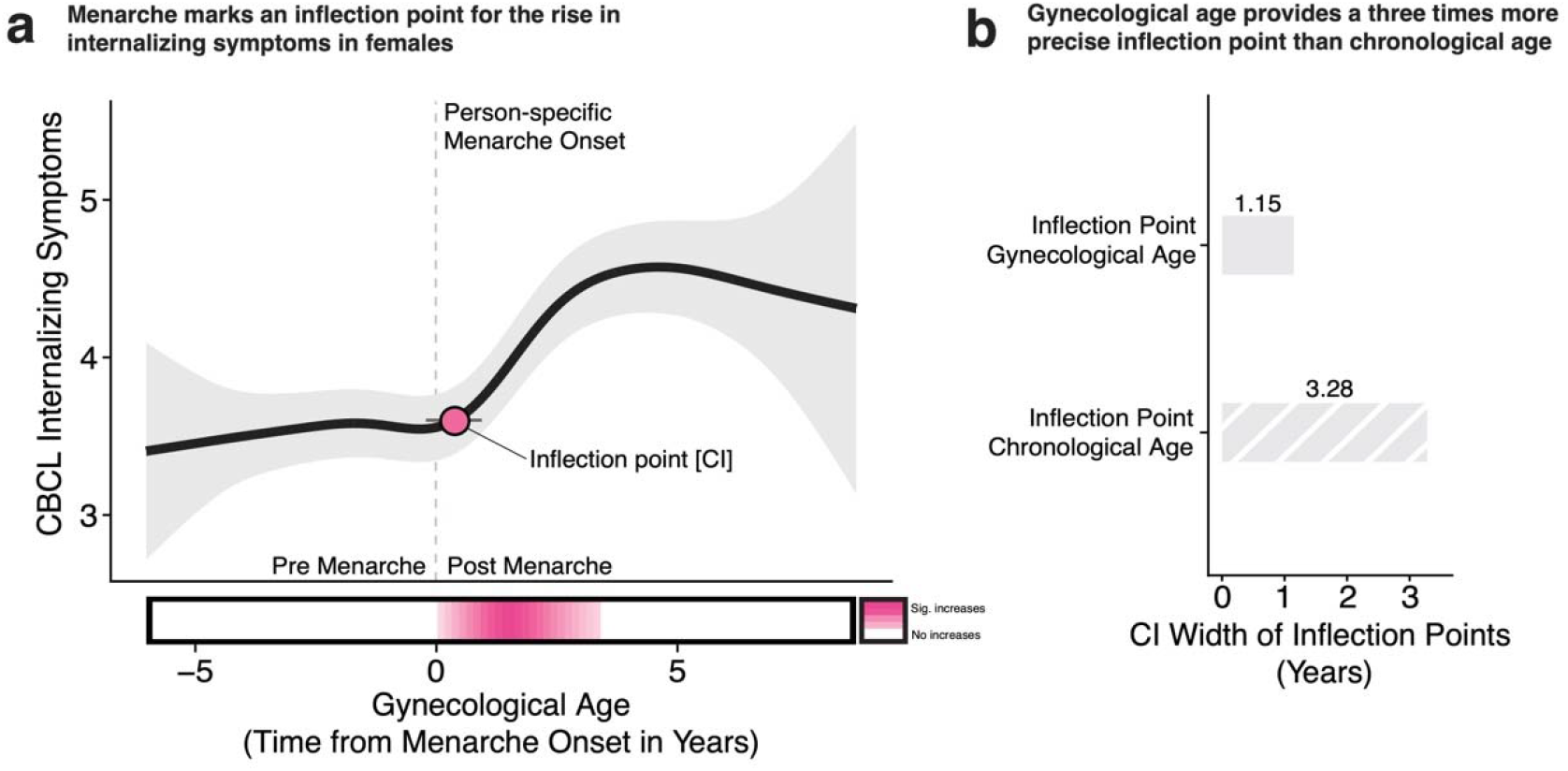
Menarche onset aligns with inflection points in the rise of internalizing symptoms (*n* = 5,016 female participants). **a**, Black line depicts predicted values from generalized additive mixed models (GAMMs) relating gynecological age (time relative to menarche onset) to Child Behavior Checklist (CBCL) internalizing symptoms (raw scores). Light gray shaded regions represent 95% credible intervals (CI). The inflection point, derived from second derivative maximum indicating the point of fastest acceleration, is shown as a pink point with a black outline. The gray horizontal line represents the 95% CI of the inflection point derived from posterior sampling. The dashed vertical line marks person-specific menarche onset (gynecological age = 0). The colored horizontal bar along the x-axis denotes the period of significant change derived from first derivative estimates: pink indicates significant increases, white indicates no significant change. Note the location of the inflection point close to menarche onset, suggesting that the rise in internalizing symptoms accelerates around the time of menarche. **b**, 95% credible interval (CI) widths for the inflection points in internalizing symptoms derived from gynecological and chronological age models, expressed in years. Gray solid and striped bars represent CI widths derived from gynecological and chronological age models, respectively. Black numerical labels on top of the bars indicate the CI width in years. Note the consistently narrower CI for gynecological age, reflecting greater temporal precision in localizing the inflection point for the rise in internalizing symptoms relative to chronological age.

We did not observe a precise menarche-related inflection point for CBCL externalizing symptoms, which decreased across adolescence (*P*_Bonferroni_<0.001; Supplementary Fig. 1, Supplementary Table 1–2). Further, while we replicated the known association between earlier menarche timing and internalizing symptoms (*P*_Bonferroni_<0.001; Extended Data Fig. 1b)^8–10,17^, this association was no longer significant when gynecological age was included in the same model (timing: *P*_Bonferroni_=0.128; gynecological age: *P*_Bonferroni_<0.001, Supplementary Table 1), indicating that the proximity to menarche, rather than relative timing, is the predominant predictor of internalizing symptoms in this sample. Overall, these findings show that while chronological age identifies an inflection point for internalizing symptoms around average menarche timing, gynecological age more precisely identifies this transition point by anchoring developmental changes to an individual’s specific menarche onset.

We next examined the development of gross brain morphology according to chronological and gynecological age. Analysis of chronological age replicated established neurodevelopmental patterns: Whole brain volume, subcortical volume, and surface area significantly increased until early-to mid-adolescence before declining, whereas gray matter volume and cortical thickness significantly decreased and white matter volume significantly increased (Extended Data Fig 6a; Supplementary Table 11). Notably, all developmental inflection points again aligned with the group-average menarche timing (between ages 11.1 and 12.4, smallest 95% CI: 11.6–12.7, widest 95% CI: 10.7–16.4, Supplementary Table 12).

Analysis of gynecological age revealed that inflection points for gross brain morphology were again tightly aligned with menarche onset: Across all gray and white matter metrics, model-estimated inflection points were identified within 0.1 to 0.4 years post-menarche onset (smallest 95% CI: 0.2–0.3, widest 95% CI: -0.7–0.4, **Fig. 2a**; Extended Data Fig 6b; Supplementary Table 12). Notably, the estimates were more precise for gynecological age, suggesting menarche onset is a precise inflection point demarcating changes in gross brain morphometry (**Fig. 2b**). In contrast, fluid-content metrics (lateral ventricle and cerebrospinal fluid volumes) showed no inflection points around menarche onset (Supplementary Table 12), suggesting structural remodeling rather than altered fluid content with menarche onset. Again, these results replicated across split-half samples (Extended Data Fig. 7, Supplementary Table 13–14), when accounting for familial dependencies (Supplementary Table 15–16), and when estimating gynecological age based on parent-report (Extended Data Fig. 5, Supplementary Table 17–18).

**Fig. 2.**
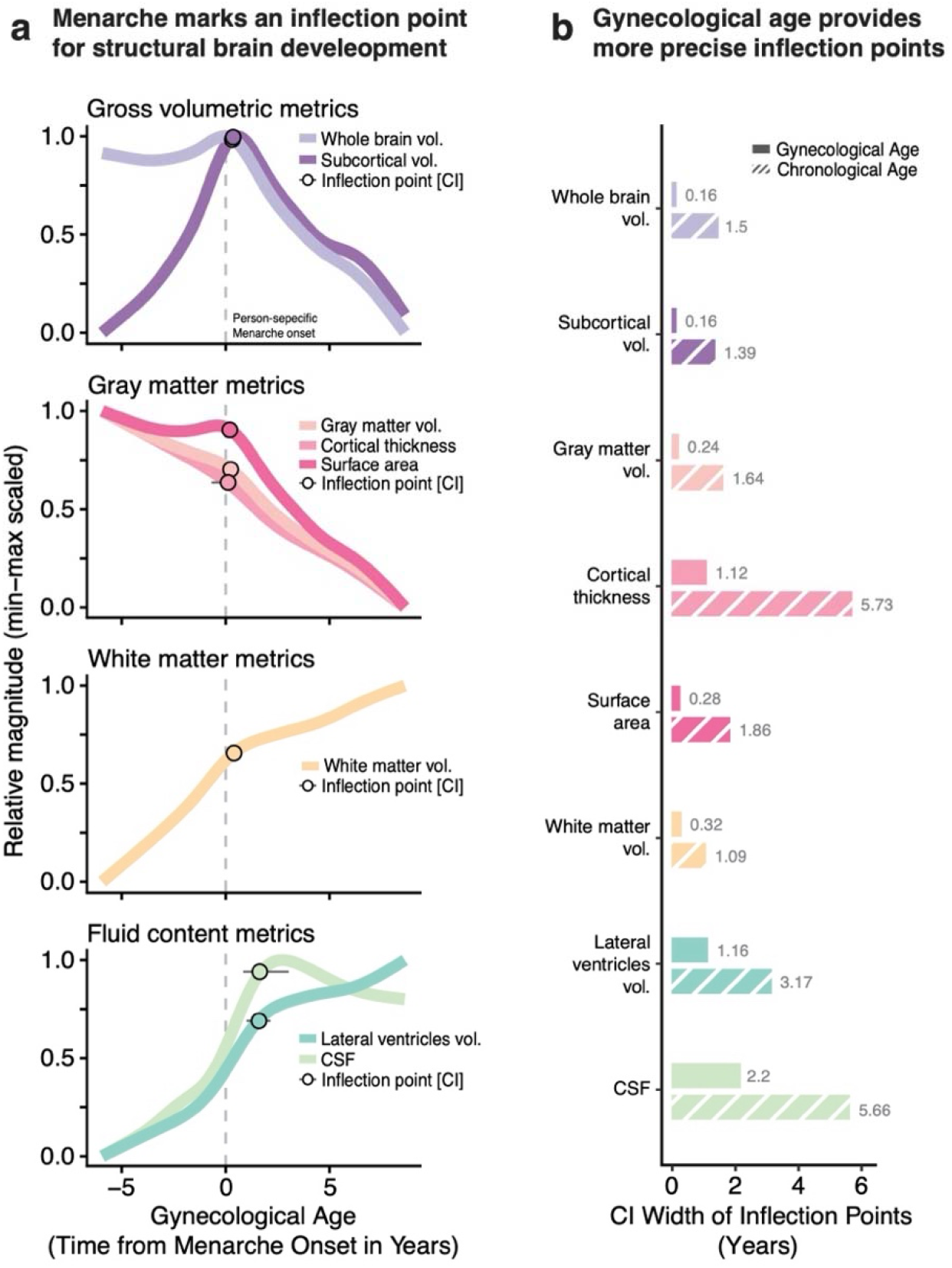
Menarche onset aligns with inflection points in structural brain development (*n* = 4,978 female participants). **a**, Colored lines depict predicted values from generalized additive mixed models (GAMMs) relating gynecological age (time relative to menarche onset) to structural brain measures. Brain measures are displayed as relative magnitude (min-max scaled) to enable comparison across measures on a common scale. Inflection points, derived from second derivative minima, are shown as colored points with black outlines. For trajectories with decreasing slopes (e.g., declining gray matter volume), the minimum of the second derivative indicates the point of greatest acceleration in decline (steepest decrease in slope). For trajectories with increasing slopes (e.g., rising in internalizing symptoms), the minimum indicates the point of greatest deceleration (where the rate of increase slows most). Gray horizontal lines represent 95% credible intervals (CI) derived from posterior sampling. Dashed vertical lines mark person-specific menarche onset (gynecological age = 0). Note the clustering of inflection points near menarche onset for brain tissue metrics: Gross volumetric, gray matter, and white matter metrics show inflection points temporally aligned with menarche, whereas fluid-content metrics did not. **b**, 95% credible intervals (CI) widths for inflection points in structural brain measures derived from gynecological and chronological age models, expressed in years. Solid colored bars represent CI widths of inflection points derived from gynecological age; striped colored bars represent CI widths of inflection points derived from chronological age. Gray numerical labels adjacent to bars indicate CI width in years. Note the consistently narrower CIs for gynecological age, reflecting greater temporal precision in localizing inflection points relative to chronological age. For **a** and **b**, CSF = cerebrospinal fluid.

Aside from cortical thickness (*P*_Bonferroni_=0.039; Supplementary Table 11), menarche timing was not significantly related to structural brain measures (all other *P*_Bonferroni_≥0.174). As with internalizing symptoms, while chronological age identified inflection points for structural brain changes around average menarche timing, gynecological age provided a more precise predictor for inflection points by anchoring developmental changes to an individual’s specific menarche onset.

Next, we examined whether menarche onset-related brain changes influenced the observed increases in internalizing symptoms. While absolute values of brain morphology (e.g., larger or smaller) did not predict internalizing symptoms (all *P*_Bonferroni_>.05; Supplementary Table 19), the individual *pace* of post-menarche change in gray matter volume (*P*_Bonferroni_=0.042) and surface area (*P*_Bonferroni_=0.012) were significantly associated with internalizing symptoms. Specifically, more rapid reductions in volume and area following menarche onset were associated with greater internalizing symptoms (Extended Data Fig. 8; Supplementary Table 20). Given that the magnitude of these associations was modest, they should be considered as preliminary evidence warranting further investigation.

To support the specificity of our findings, we conducted multiple sensitivity analyses. First, we repeated all main analyses incorporating interaction terms to examine whether menarche onset effects were moderated by menarche timing (early versus late) and general exposome level (low versus high), beyond simply treating these variables as covariates. Crucially, the main effect of menarche onset on both internalizing symptoms and brain morphometry remained significant, consistent with our primary models. However, we observed that the magnitude of this effect was context-dependent: Increases in internalizing symptoms following menarche were steeper in the high exposome group (with lower environmental adversity and higher socioeconomic status) and with earlier menarche timing. While brain development continued to reveal an inflection point at menarche onset, the extent of brain change was moderated by both general exposome and menarche timing (Extended Data Fig. 9–10; Supplementary Table 21–22).

Finally, to ensure that menarche onset, rather than other pubertal milestones, best explained changes in brain and mental health, we repeated analyses using both youth self- and parent-reported onset of alternative puberty markers: Growth spurt, breast development, skin changes, and body hair growth. No inflection points for either internalizing symptoms nor structural brain changes aligned with the onset of any alternative puberty markers (Extended Data Fig. 5, Supplementary Tables 9–10,23–26).

These findings support menarche onset as an inflection point for changes in brain and mental health trajectories, suggesting that this normative, easily identifiable milestone is not only a reproductive event but a significant neuroendocrine marker of adolescent brain development and mental health risk. Menarche marks the activation of cyclical fluctuations in gonadal hormones that target estrogen and progesterone receptors throughout the brain and are involved in developmental processes such as synaptogenesis and myelination^18–21^. The onset of ovarian cycling introduces a powerful neuroendocrine signal that may drive the observed inflection points in brain morphometry, creating a period of heightened neuroplasticity and subsequent mental health vulnerability. Yet menarche remains an underutilized milestone in clinical practice, often documented but rarely integrated into mental health risk assessment. By destigmatizing conversations around menstruation^22^ and recognizing menarche as a critical developmental marker, clinicians can leverage this observable biological event to identify when the window of mental health vulnerability is open for each individual. Current guidelines that recommend depression screening at age 12^23^ are imprecise, especially for those reaching menarche before the average age of 11.8. This precision is critical for targeted screening: Rather than universal age-based monitoring, clinicians should initiate mental health screenings at the observable milestone of person-specific menarche onset, after which risk acceleration begins. Importantly, incorporating gynecological age as a core developmental marker in neuroscience and mental health research can refine models of adolescent neurodevelopment by anchoring the unfolding of brain development to this specific pubertal milestone.

## Supporting information

Legends for Extended Data Figures

## Methods

### Data source

The ABCD Study (https://abcdstudy.org) is a prospective, observational, 10-year longitudinal investigation into brain development, spanning from ages 9 to 10 years at baseline through adulthood and comprising 21 study sites^1^. The study’s rationale and design aspects have been described in detail elsewhere^2^. Approval for the ABCD Study was given from the central Institutional Review Board at the University of California, San Diego and from the local Institutional Review Boards at University of Maryland, Baltimore, University of Colorado, Boulder, University of Minnesota, the Laureate Institute for Brain Research, Oregon Health and Science University, University of Vermont, University of Pittsburgh, Virginia Commonwealth University, University of Rochester, University of Florida, Medical University of South Carolina, University of Michigan, University of Utah, SRI International, University of Wisconsin-Milwaukee, Children’s Hospital of Los Angeles, Florida International University, Washington University in St. Louis, and Yale University. All procedures performed in this study involving human participants were in accordance with the ethical standards of the institutional and/or national research committee and with the 1964 Helsinki declaration and its later amendments or comparable ethical standards.

### Sample selection

Data used in these analyses (v6.0; https://dx.doi.org/10.15154/z563-zd24) included 5,673 female youth across up to seven visits at annual intervals. Participants were required to have valid baseline measures of socio-environmental variables, yielding a sample of 5,655 individuals, of whom 5,016 additionally provided information on age at menarche. Sample sizes and mean ages at each visit were: baseline, *n* = 5,016, *M* = 9.96 years (*SD* = 0.62); follow-up 1, *n* = 4,873, *M* = 10.97 (*SD* = 0.64); follow-up 2, *n* = 4,847, *M* = 12.07 (*SD* = 0.67); follow-up 3, *n* = 4,690, *M* = 12.95 (*SD* = 0.65); follow-up 4, *n* = 4,422, *M* = 14.17 (*SD* = 0.71); follow-up 5, *n* = 4,084, *M* = 15.07 (*SD* = 0.67); follow-up 6, *n* = 2,342, *M* = 16.07 (*SD* = 0.65). Approximately 90% of participants had five or more visits. Analyses involving mental health measures included *n* = 5,016 participants with valid internalizing and/or externalizing symptom data from at least one assessment. Analyses involving neuroimaging measures included *n* = 4,978 participants who had structural MRI data from at least one assessment that passed quality control checks. Neuroimaging measures were acquired biennially. Parents and guardians provided written informed consent as part of the ABCD study.

### Menarche timing variable

A total of 5,016 adolescents provided information on age at menarche, assessed via youth self-report using items from the Pubertal Development Scale (PDS) administered annually. Participants were asked, “*Have you begun to menstruate (started to have your period)?*” (variable: *ph_y_pds_f_002*), and, if yes, “*How old were you when you started to menstruate?*” (variable: *ph_y_pds_f_002_01*). Age at menarche was coded as missing if participants reported an age at menarche but did not indicate that menstruation had begun. In seven cases, implausible values (e.g., age at menarche = 0) were identified; these values were recoded based on consistent reports at other follow-up assessments (e.g., set to 9 years when supported by follow-up data). For participants with inconsistent reports of age at menarche across assessments (e.g., reporting age 10 at one visit and age 11 at subsequent visits), the median reported age was used.

For sensitivity analyses, we also assessed age at menarche via the parent-report using items from the PDS. Parents were asked, “*Has your child begun to menstruate?*” (variable: ph_p_pds f_002), and if yes, “*How old were they when they started to menstruate?*” (variable: ph_p_pds f_002 01). Age at menarche was coded as missing if parents reported an age at menarche but did not indicate that menstruation had begun. In case of implausible values (e.g., age at menarche = 99); these values were recoded based on consistent reports at other follow-up assessments or set to missing. For participants with inconsistent reports of age at menarche across assessments (e.g., reporting age 10 at one visit and age 11 at subsequent visits), the median reported age was used.

Both youth and parent report of menarche timing correlated with *r* = 0.85, making it a consistent measure across both raters.

### Gynecological age variable

To derive a person-specific metric centered on menarche *onset*, we calculated gynecological age subtracting each participant’s age at menarche from their chronological age at each assessment timepoint. This transformation centers the variable at zero, where zero denotes menarche onset, negative values indicate the time before menarche (e.g. –1 = one year before menarche), and positive values indicate the time after menarche (e.g. +2 = two years post-menarche). This approach aligns all participants to a common developmental reference point, their person-specific menarche onset, allowing us to examine developmental trajectories relative to this biological milestone rather than chronological age alone. Gynecological age and menarche timing were correlated with *r* = -0.44, chronological age and menarche timing with *r* = 0.08, and gynecological and chronological age with *r* = 0.86. The strong correlation between chronological and gynecological age indicates collinearity, precluding their simultaneous inclusion in statistical models. Menarche timing showed low correlations with both variables and could therefore be retained in all models.

Gynecological age was additionally derived from parent-reported menarche onset for sensitivity analyses. Gynecological age and menarche timing were correlated with *r* = -0.44, chronological age and menarche timing with r = 0.07, and gynecological and chronological age with r = 0.86. The strong correlation between chronological and gynecological age indicates collinearity, precluding their simultaneous inclusion in statistical models. Menarche timing showed low correlations with both variables and could therefore be retained in all models.

### Alterative puberty markers

To evaluate the specificity of our findings with regard to menarche onset, we assessed four alternative puberty markers using items from the PDS: growth spurt, breast development, body hair growth, and skin changes. The alternative puberty markers differ from menarche in several important ways that bear on their interpretation: First, growth spurt, body hair growth, and skin changes reflect adrenarche, while breast development reflects gonadarche (the same neuroendocrine axis as menarche), and menarche itself is the latest-occurring of all markers assessed^3^, meaning it is unsurprising that it carries distinct developmental information relative to the other markers. Second, while menarche timing was consistently reported across parent and youth raters (*r* = 0.85), earlier pubertal changes are known to be better captured by parent report, as youth self-awareness of these gradual processes may be limited^4,5^. We therefore assessed each alternative marker using both parent and youth report. For each alternative puberty marker we derived the onset age as follows: Growth spurt onset age was derived from variables *ph_y_pds_001* and *ph_p_pds_001*, breast development onset age from variables *ph_y_pds_f_001* and *ph_p_pds_f_001*, body hair growth onset age from variables *ph_y_pds_002* and *ph_p_pds_002*, and skin changes onset age from variables *ph_y_pds_003* and *ph_p_pds_003*. For each marker, onset age was defined as the age at the first study visit where the parent or participant self-reported the marker had ‘barely started’ (PDS score = 2) or was ‘definitely underway’ (PDS score = 3). Finally, to derive a person-specific index of puberty marker onset, age at puberty marker onset was subtracted from chronological age at each assessment, making the variables to be centered at zero to denote the time of puberty marker onset, with negative values indicating time before puberty marker onset and positive values indicating time after puberty marker onset. This was done for growth spurt, breast development, body hair growth, and skin changes separately.

### Mental health variables

Females are twice as likely as males to experience depressive symptoms across the lifespan, with this sex differences emerging during adolescence^6^. To assess this sex-specific vulnerability in females, we examined internalizing symptoms, which comprise depressive and anxiety symptoms, using the parent-reported Child Behavior Checklist (CBCL)^7^ for ages 6–18. We also examined externalizing symptoms to determine the specificity of internalizing symptom increases to females, given that externalizing symptoms show a different developmental pattern and are more prevalent in males^8^. The CBCL is a widely used assessment comprising 120 items rated on a 3-point scale (0 = no true; 1 = somewhat/sometimes true; 2 = very true/often true) based on the child’s behavior over the past 6 months. The CBCL organizes scores into eight syndrome scales; Anxious-depressed, withdrawn-depressed, somatic complaints, rule-breaking behavior, aggressive behavior, social problems, thought problems, and attention problems. These can be combined into broader indices: internalizing symptoms (comprising anxious-depressed, withdrawn-depressed, and somatic complaints; variable name: *mh_p_cbcl_synd_int_sum*) and externalizing problems (comprising rule-breaking behavior and aggressive behavior; variable name *mh_p_cbcl_synd_ext_sum*). Here, we utilized the tabulated raw sum scores. The CBCL was administered annually across baseline and the six follow-ups.

For sensitivity analyses, we also derived internalizing symptoms from the youth self-reported Brief Problem Monitor (BPM)^9^. The BPM is a brief 19-item instrument derived from the longer (CBCL), Teacher’s Report Form, and Youth Self-Report, and is designed for repeated monitoring of emotional and behavioural functioning. Each item is rated on a 3-point scale (0 = no true; 1 = somewhat/sometimes true; 2 = very true/often true) over the past week. The raw sum score of the internalising problems subscale (variable name: *mh_y_bpm_int_sum*) was used in the present study. Ten BPM assessments were available: one six months follow-up after baseline, semi-annual assessments for follow-up 1, 2, and 3, and annual assessments for follow-up 4, 5, and 6.

### Structural brain metrics

All individuals included in this study underwent biannual 3 Tesla structural MRI scans according to standardized protocols^10^. T1-weighted images were acquired at local study sites, while processing and quality control assessments were conducted at the Data Analysis, Informatics and Resource Center (DAIRC) of the ABCD Study^11^. FreeSurfer v5.3.0 (https://surfer.nmr.mgh.harvard.edu) was used, following the ABCD Study’s standardized processing pipeline, to process the images and obtain structural brain measures from cortical 34 and eight subcortical regions within each brain hemisphere from the Desikan-Killiany atlas^12^. Quality control, including both automated procedures and expert visual inspection was performed by DAIRC. For further details on processing and quality control procedures of raw and processed data, see Hager et al. (2019)^11^. We used the following variables for our analyses: whole brain volume (variable name: *mr_y_smri_vol_aseg_whb_sum*), subcortical volume (variable name: *mr_y_smri_vol_aseg_scgv_sum*), gray matter volume (variable name: *mr_y_smri_vol_dsk_sum*), cortical thickness (variable name: *mr_y_smri_thk_dsk_mean*), surface area (variable name : *mr_y_smri_area_dsk_sum*), white matter volume (variables names: *mr_y_smri_vol_aseg_cwm_lh_sum, mr_y_smri_vol_aseg_cwm_rh_sum*), lateral ventricle volume (variable name: *mr_y_smri_vol_aseg_lvs_sum*), and CSF (variable name: *mr_y_smri_vol_aseg_csf_sum*). Only data that passed quality control checks and were recommend for inclusion were used for analyses (variable name: *mr_y_qc_incl_smri_t1_indicator*).

### Socio-environmental variables

To account for socio-environmental factors known to influence both psychopathology^13^ and menarche timing^14^, we quantified the *general exposome*^15,16^. The general exposome represents the totality of co-occurring environmental exposures and experiences that an individual encounters throughout the lifespan^17^, in contrast to examining single environmental features (e.g. household income) in isolation. The framework has been previously applied to and validated in the ABCD dataset^18–21^, combining youth-report, parent-report, and geocoded data from the child’s home address to derive a composite variable capturing co-occurring physical/chemical exposures (e.g. air pollution), psychosocial experiences (e.g. family dynamics), and broad socioeconomic context (e.g. area deprivation index). Here, we used the general exposome score derived from a longitudinal bifactor analysis^19^the baseline (ages 9–10) assessment, wherein higher scores reflect higher socioeconomic status and lower environmental adversity. Sensitivity analyses using race/ethnicity, area deprivation index, and combined household income separately yielded consistent results with the general exposome. We therefore included the general exposome as a composite covariate rather than controlling for these factors separately, providing more parsimonious models.

### Statistical approach

#### Software and packages

All statistical analyses were conducted using RStudio (version 2024-04-24; R version 4.4.0) with the following R packages: *tidyverse* (version 2.0.0), *mgcv* (version 1.9-3), *gratia* (version 0.11.1), *lme4* (version 1.1-37), and *parallel* (version 4.4.0). Data visualizations were generated using *ggplot2* (version 4.0.0), and *ggpattern* (version 4.0.0).

#### Generalized additive mixed models

We used GAMMs as our primary analytical framework to assess relationships of mental health symptoms and brain measures with chronological age, menarche timing, and gynecological age. GAMMs offer several advantages for examining developmental trajectories: They allow predictor variables to influence outcomes through smooth, nonlinear functions while penalizing complexity to avoid overfitting, thereby flexibly capturing linear and nonlinear relationships without requiring a priori specification of functional form. This approach is particularly well-suited for developmental data where relationships may exhibit complex, non-monotonic patterns such as periods of acceleration, deceleration, or inflection points.

In all GAMMs, the default value of *k* = 10 was used as an upper limit of for degrees of freedom of each smooth function, allowing sufficient flexibility while setting an upper limit on nonlinear complexity. We included data collection site and participant ID as random intercepts in all models to account for nested data structure and repeated measures. Mental health outcomes (internalizing and externalizing symptoms) were modeled using a negative binomial family, whereas brain measures (whole brain volume, subcortical volume, gray matter volume, cortical thickness, surface area, white matter volume, lateral ventricles volume, CSF) were modeled using a Gaussian family. All models were fit using restricted maximum likelihood (REML) estimation. Correction for multiple comparisons was performed using Bonferroni adjustment. Corrections were applied separately within each family of models: Models with mental health variables as outcomes and models with brain measures as outcomes.

#### Calculation of inflection points

We extracted key developmental features to characterize trajectory dynamics from fitted models. First derivatives of the smooth functions were computed to identify periods of significant increases and decreases in mental health symptoms and brain measures across development. Second derivatives were then computed to quantify the rate of change in slopes (i.e., acceleration or deceleration). We identified critical transition points by extracting the extrema (minima and maxima) of the second derivatives. For trajectories with increasing slopes (e.g., rising in internalizing symptoms), the maximum of the second derivative indicates the point of greatest acceleration (steepest increase in slope), while the minimum indicates the point of greatest deceleration (where the rate of increase slows most). For trajectories with decreasing slopes (e.g., declining gray matter volume), the minimum of the second derivative indicates the point of greatest acceleration in decline (steepest decrease in slope), while the maximum indicates the point of greatest deceleration in decline (where the rate of decrease slows most). We refer to these points as *inflection points* throughout this paper to denote critical transition periods in development. Credible intervals for second derivative extrema were estimated using posterior sampling with 1,000 draws from the posterior distribution of the fitted smooth functions. This approach captures the uncertainty around the estimated timing of each inflection point, providing a precision estimate that quantifies the range within which the true inflection point is likely to occur.

#### Evaluation of socio-environmental context as a covariate predicting menarche timing

Pubertal timing has been previously shown to depend on socio-environmental context. We first replicated this association in the study sample^8^. We fit a GAMM with menarche timing as the outcome variable and general exposome as a smooth predictor variable, with a random intercept for data collection site. The random intercept for participant ID was not necessary in this case because each participant only has one value for menarche timing and general exposome. As expected, we observed a significant association and thus retained general exposome as a covariate in all subsequent analyses.

### Mental health analyses

#### Evaluation of menarche timing on mental health

Mental health symptoms are known to be associated with menarche timing^22,23^. To quantify this association in the study, we fit separate GAMMs for each mental health outcome with menarche timing as a smooth predictor term *s*(). The following model formula was implemented in R:

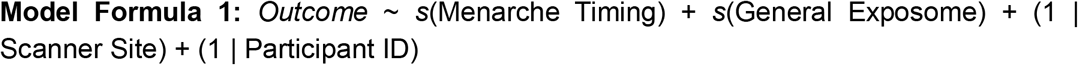

We evaluated the significance and shape of the fitted smooth term for menarche timing, identifying periods of significant change and the location of inflection points in the developmental trajectories. The fitted smooth term for menarche timing approximated a linear relationship and thus did not produce inflection points requiring extraction of second derivative extrema.

#### Evaluation of chronological age on mental health

Internalizing and externalizing mental health symptoms are known to fluctuate over the adolescent years^6,24–26^. To quantify this association in the study sample, we fit separate GAMMs for each mental health outcome (internalizing and externalizing symptoms) with chronological age as a smooth predictor term *s*(). The following model formula was implemented in R:

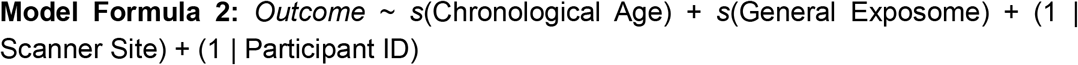

We evaluated the significance and shape of the fitted smooth term for chronological age, identifying periods of significant change and the location of inflection points in the developmental trajectories. Inflection points were identified by extracting the extrema (minima and maxima) of the second derivatives.

#### Evaluation of gynecological age on mental health

Our analysis of chronological age suggested an inflection point in internalizing symptoms falling at the average age of menarche. To test whether taking into account person-specific menarche timing would improve our ability to capture this critical developmental turning point, we repeated our analysis using gynecological age. Specifically, we fit GAMMs with gynecological age as a primary smooth predictor while co-varying for menarche timing to isolate the effect of time relative to menarche onset from the effect of early versus late menarche timing. The following model formular was implemented in R:

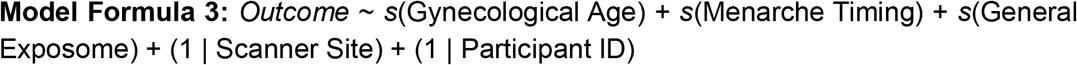

We then evaluated the significance and shape of the fitted smooth term for gynecological age, identifying periods of significant change and the location of inflection points in the developmental trajectories. As for chronological age, inflection points were identified by extracting the extrema (minima and maxima) of the second derivatives.

#### Comparison of inflection points for chronological age and gynecological age

To evaluate whether gynecological age provides more precise estimates of developmental inflection points than chronological age, we compared the credible intervals of the second derivative extrema from model formulas 2 and 3. Narrower intervals indicate less error in our estimate of the location, indicating a sharper and more precise estimate.

### Brain measure analyses

We repeated model formulas 1–3 for brain measures (whole-brain volume, subcortical volume, gray matter volume, cortical thickness, surface area, white matter volume, lateral ventricles volume, and CSF), following the same analytical approach as described for mental health outcomes. We examined the developmental trajectories for both chronological and gynecological age and extracted the inflection points. As above, we compared the credible intervals of our estimates to evaluate precision.

### Moderation analyses

We next examined whether brain morphometric changes following menarche moderate the observed relationship between gynecological age and internalizing symptom severity. We tested this in two complementary ways: First, by examining whether *absolute brain morphometry* (i.e., brain size or thickness at a given time point) moderates symptom trajectories; and second, by examining whether individual *rates of brain change* from pre-to post-menarche moderate symptom trajectories. The first approach tests whether brain morphometry itself (bigger or smaller) matters for mental health vulnerability, while the second tests whether the individual pace of brain change (faster or slower) across the menarche transition predicts symptom severity.

### Brain morphometry as a moderator

We fit separate GAMMs for each brain measure with internalizing symptoms as outcome. These models included main effects of gynecological age and brain measures, plus a tensor product interaction term *ti*() to test the moderation. A significant interaction indicates the association between gynecological age and symptoms differs by more or less absolute brain volume. The following model formula was implemented in R:

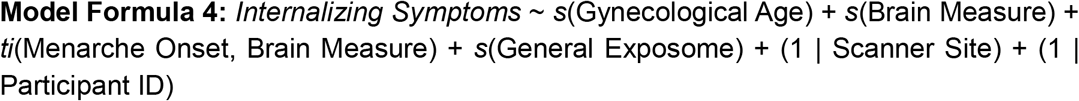

#### Individual post-menarche brain change as a moderator

While model formula 4 tested whether absolute brain morphometry moderates symptom severity, we further hypothesized that individual rates of change in brain structure following menarche might predict symptom severity, irrespective of between-person differences in brain size. To test this, we used a *two-stage broken-stick modeling approach*. In the first stage, we estimated individual-specific post-menarche brain change slopes using linear mixed-effects (LME) models. We created a *time post menarche* variable by setting all pre-menarche values in gynecological age to 0 (values before menarche onset were coded as 0, while post-menarche values reflected the elapsed time since menarche). This specification allows the brain trajectory to have a “break point” at menarche, with potentially different slopes before and after menarche. We fit the following LME model formula for each brain metric in R:

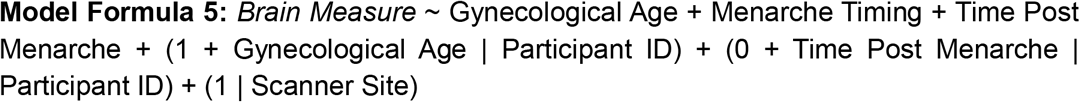

This model included: fixed effects of gynecological age (overall trend), menarche timing (early versus late), and time post menarche (post-menarche slope), random intercepts for participant ID and data collection site, random slopes for gynecological age (allowing individual variation in overall trajectories) and time post menarche (allowing individual variation in post-menarche change rate). In the second stage, we extracted each participant’s estimated post-menarche brain slope from the LME models and tested whether these individual slopes predicted internalizing symptoms, using GAMMs with a tensor product interaction term *ti*(). The following model formula was implemented in R:

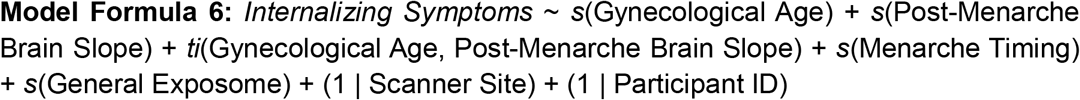

This two-stage approach tests whether faster or slower rates of brain change following menarche onset are associated with greater symptom severity, and whether this this association varies across gynecological age. This approach focuses on within-person changes relative to their own pre-menarche baseline rather than absolute brain morphometric values.

### Sensitivity analyses

To confirm the specificity of our findings, we conducted a series of sensitivity analyses as described below.

#### Moderating effects of menarche timing and exposome

Our primary models (model formula 3) included menarche timing and general exposome as covariates, controlling for their main effects. However, these factors might also moderate the effect of gynecological age on internalizing symptoms and brain measures. For example, the menarche onset might only be an inflection point for internalizing symptoms in early menarche timing but not in late menarche timing. To test this, we repeated model formula 3 analyses incorporating tensor product interactions *ti*(). The following model formula was implemented in R:

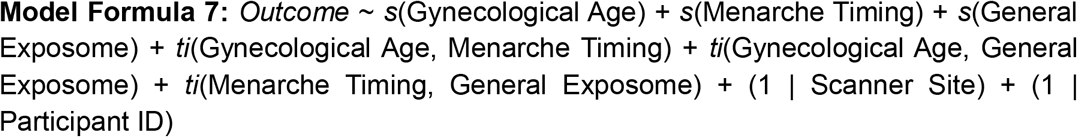

#### Split-half reliability

To test the robustness and replicability of our findings, we replicated our primary analyses using the ABCD Analyst-Recommended Matched Samples (ARMS) framework^27^. The ARMS approach provides two independent, socio-demographically balanced samples (*n* = 2,450 and *n* = 2,395) matched on key demographic variables including race/ethnicity, socioeconomic status, and study site. This split-half reliability analysis tests whether our findings replicate across independent subsamples, providing evidence for the stability of the observed effects. Bonferroni correction was applied across both samples.

#### Familial dependencies

To account for non-independence introduced by siblings in the dataset, we reran our primary analyses restricting the sample to one randomly selected sibling per family (*n* = 4,398). This analysis tests whether our findings were independent of familial affiliation.

#### Comparison of alternative pubertal milestones

To determine whether our findings were specific to menarche or generalize to the onset of other pubertal milestones, we repeated model formula 3 substituting gynecological age (time since menarche) with alternative developmental timelines centered on alternate PDS items reflecting growth spurt, breast development, skin changes, and body hair growth. For each alternate developmental variable, we calculated time since onset using the same approach as gynecological age. Each alternative marker was tested in separate models. Bonferroni correction was performed across all puberty markers. This analysis tested whether developmental inflection points in brain and mental health align specifically with menarche onset or whether they occur at the onset of other pubertal events, thereby establishing whether menarche uniquely reflects organization in adolescent neurodevelopment. This analysis was done for both youth- and parent-reported PDS items.

#### Rater bias

To evaluate whether our findings were sensitive to the reporter (youth vs parent), we ran our primary analyses replacing parent-reported internalizing symptoms derived through the CBCL with youth self-reported internalizing symptoms derived through the BPM. We further compared our results when to models calculating gynecological age based on parent-reported menarche timing rather than youth-report. As described above, we also exhaustively evaluated the role of alternative puberty markers across all youth and parent-reported data.

## Data availability

The dataset that supports the findings of this analysis is under controlled access, available on the ABCD Data Repository through the NIH Brain Development Cohort (NBCD) Data Hub (https://abcdstudy.org).

## Code availability

All analysis scripts are publicly available on our GitHub repository: https://github.com/carina-heller/ABCD_menarche_project

## Acknowledgments

Data used in the preparation of this article were obtained from the Adolescent Brain Cognitive Development (ABCD) Study (https://abcdstudy.org), held in the NIMH Data Archive (NDA). This is a multisite, longitudinal study designed to recruit more than 10,000 children aged 9-10 and follow them over 10 years into early adulthood. The ABCD Study is supported by the National Institutes of Health Grants (U01DA041022, U01DA041028, U01DA041048, U01DA041089, U01DA041106, U01DA041117, U01DA041120, U01DA041134, U01DA041148, U01DA041156, U01DA041174, U24DA041123, U24DA041147). A full list of supporters is available at https://abcdstudy.org/nih-collaborators. A listing of participating sites and a complete listing of the study investigators can be found at https://abcdstudy.org/principal-investigators.html. ABCD consortium investigators designed and implemented the study and/or provided data but did not necessarily participate in analysis or writing of this report. This manuscript reflects the views of the authors and may not reflect the opinions or views of the NIH or ABCD consortium investigators. The ABCD data repository grows and changes over time. The ABCD data used in this report came from https://dx.doi.org/10.15154/z563-zd24 (Release 6.0). This work was supported by the German Research Foundation (grant number: 544183227 to C.H.), the P&S Fund and the Brain and Behavior Research Foundation Young Investigator Award (grant number 33219 to C.H.), the Brain and Behavior Research Foundation Young Investigator Award (grant number: 32974 to A.S.K.) and the National Institute of Mental Health (R00MH127293 to B.L.).

## Author contributions

C.H. was responsible for conceptualization, methodology, data curation, formal analysis, visualization, funding acquisition, and writing the original draft of the manuscript. H.S.-T. was responsible for code review and contributed to visualization and writing (review and editing). M.G. contributed to methodology and writing (review and editing). S.K. contributed to methodology and writing (review and editing). J.M.F. contributed to visualization and writing (review and editing). R.B. contributed to methodology and writing (review and editing). T.M.M. contributed to methodology and writing (review and editing). K.I.I. contributed to interpretation of results and writing (review and editing). D.A.F. contributed to visualization, interpretation of results, and writing (review and editing). B.T.-C. contributed to methodology, visualization, interpretation of results, and writing (review and editing). A.S.K. contributed to methodology, interpretation of results, and writing (review and editing). A.M.B. contributed to interpretation of results and writing (review and editing). E.G.J. contributed to interpretation of results and writing (review and editing). B.L. was responsible for supervision and resources and contributed to conceptualization, methodology, formal analysis, visualization, funding acquisition, and writing (review and editing).

## Extended Data Figures

**Extended Data Fig. 1.**
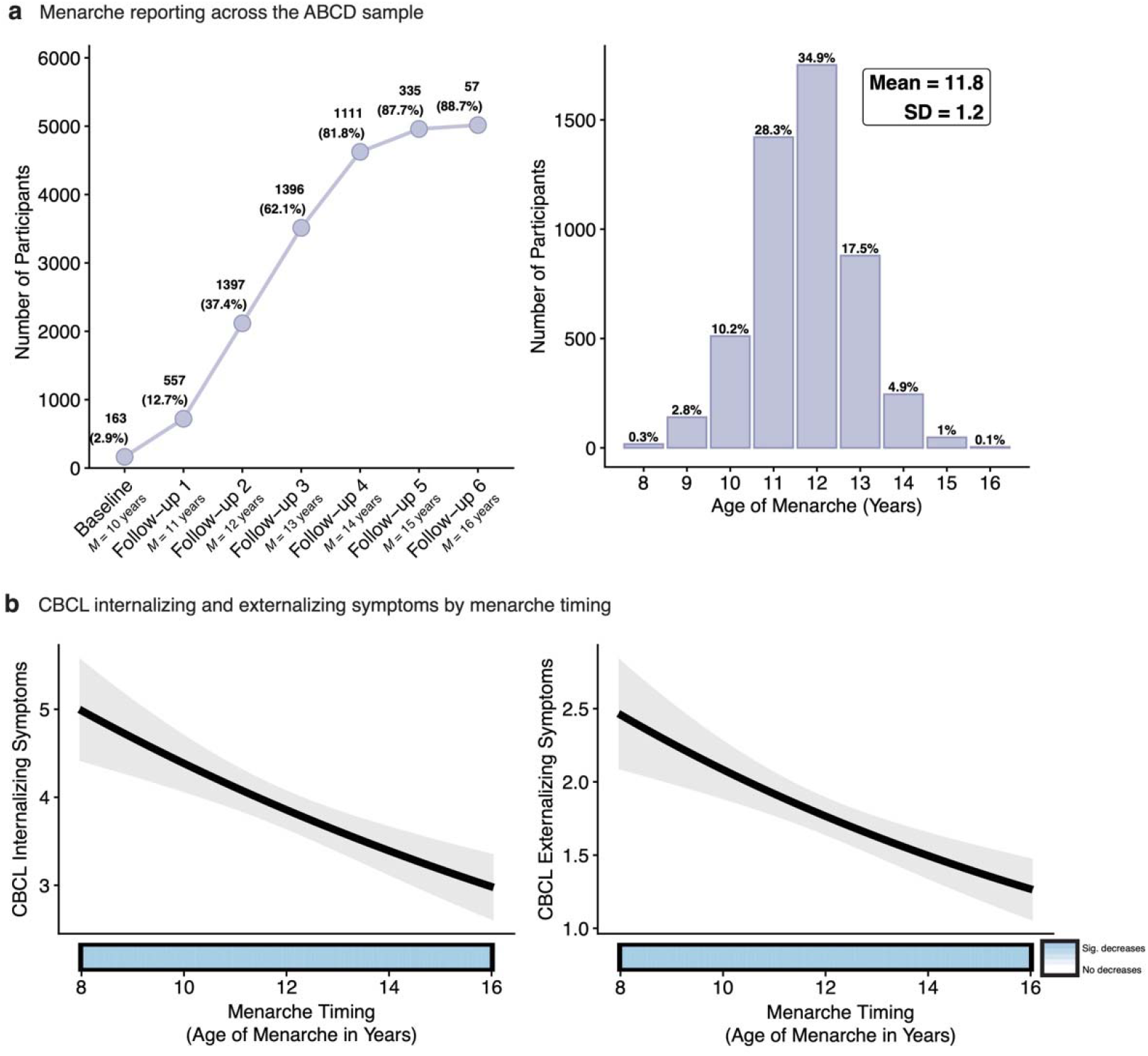
Menarche reporting and associations between menarche timing and internalizing and externalizing symptoms. **a**, Distribution of self-reported menarche across study timepoints (top) and menarche timing (bottom) in the ABCD Study. Line graph displays the cumulative number and percentage of participants reporting menarche at each assessment from baseline (mean age 10 years) through 6-year follow-up (mean age 16 years). The histogram displays self-reported menarche timing across the sample, with an average age of onset at 11.8 years (SD = 1.2 years). **b**, Child Behavior Checklist (CBCL) internalizing and externalizing symptom trajectories in association with menarche timing. Black lines depict predicted values from generalized additive mixed models (GAMMs) relating menarche timing to internalizing and externalizing symptoms. Light gray shaded regions represent 95% credible intervals (CI). Colored horizontal bars along the x-axis denote periods of significant change based on first derivative estimates: red indicates significant increases, blue indicates significant decreases. *n* = 5,016 female participants.

**Extended Data Fig. 2.**
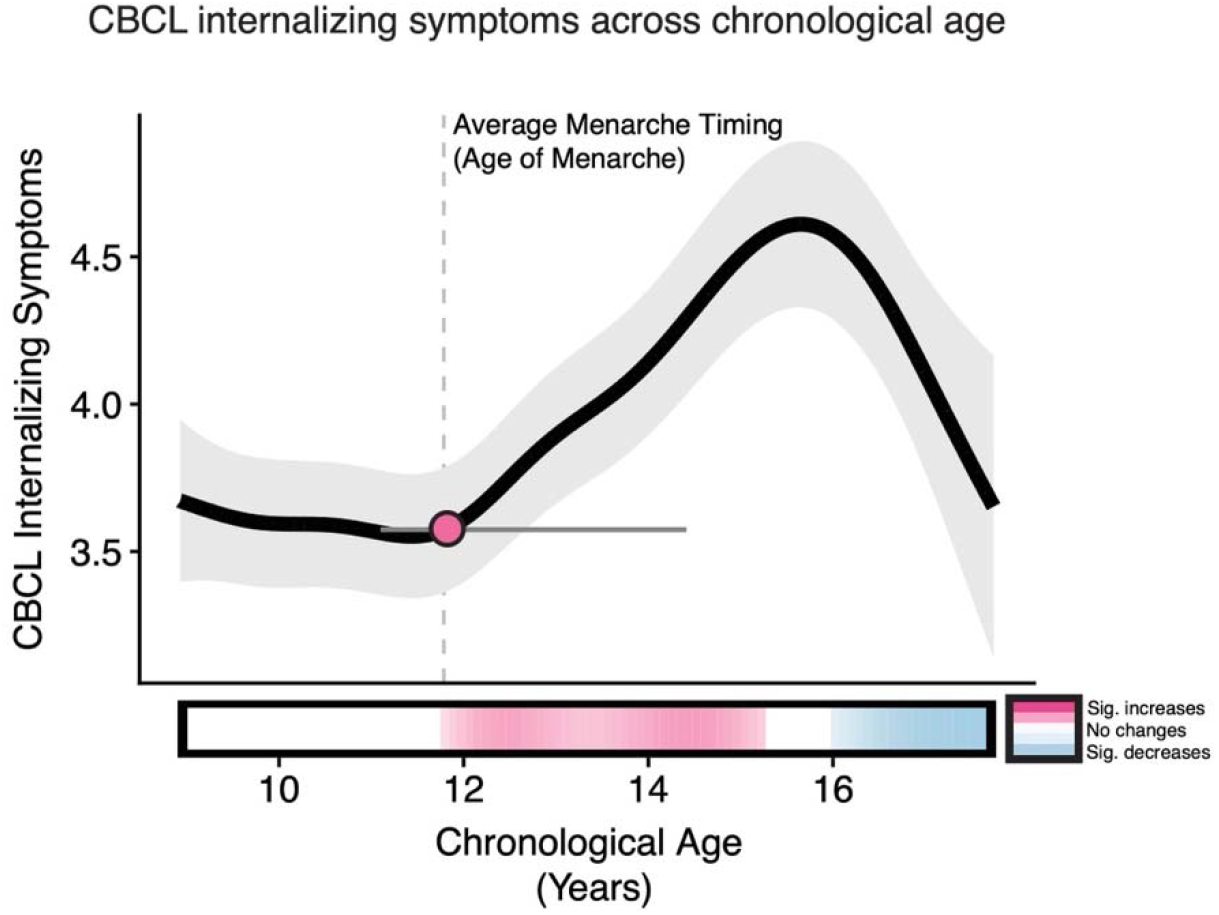
Internalizing symptoms across chronological age. Child Behavior Checklist (CBCL) internalizing symptom trajectory across chronological age. The black line depicts predicted values from generalized additive mixed models (GAMMs) relating chronological to internalizing symptoms. Light gray shaded regions represent 95% credible intervals (CI). The inflection point (derived from the second derivative maximum, indicating the point of fastest acceleration) is shown as a pink point with black outline, with 95% CI derived from posterior sampling depicted as a gray horizontal line. The colored horizontal bar along the x-axis denotes periods of significant change based on first derivative estimates: pink indicates significant increases, blue indicates significant decreases, white indicates no significant change. The dashed vertical line marks average menarche age of the sample. *n* = 5,016 female participants.

**Extended Data Fig. 3.**
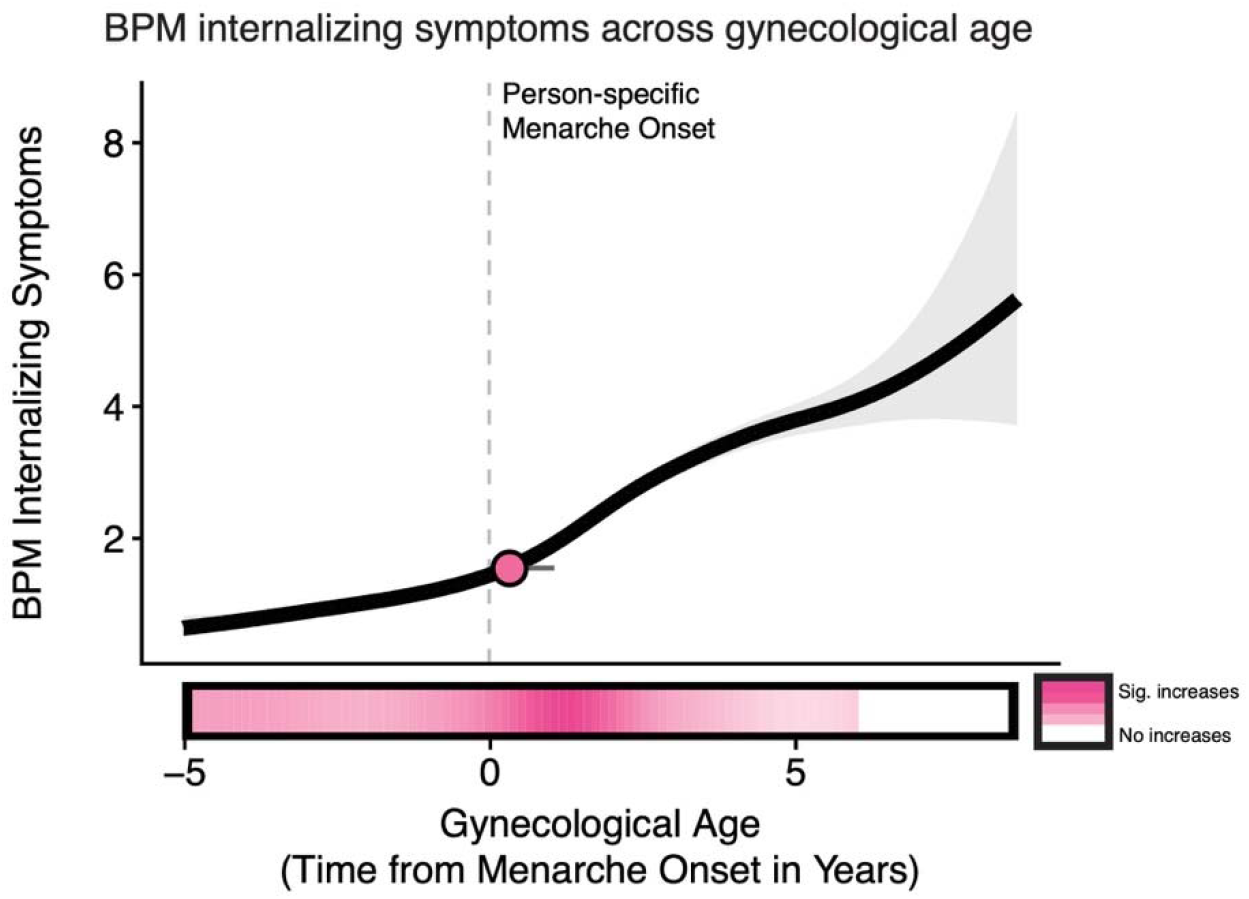
Internalizing symptoms across gynecological age. Brief Problem Monitor (BPM) youth-reported internalizing symptom trajectory across gynecological age. The black line depicts predicted values from generalized additive mixed models (GAMMs) relating gynecological age to internalizing symptoms. Light gray shaded regions represent 95% credible intervals (CI). The inflection point (derived from the second derivative maximum, indicating the point of fastest acceleration) is shown as a pink point with black outline, with 95% CI derived from posterior sampling depicted as a gray horizontal line. The colored horizontal bar along the x-axis denotes periods of significant change based on first derivative estimates: pink indicates significant increases, white indicates no significant change. The dashed vertical line marks the person-specific menarche onset (gynecological age = 0). *n* = 5,016 female participants.

**Extended Data Fig. 4.**
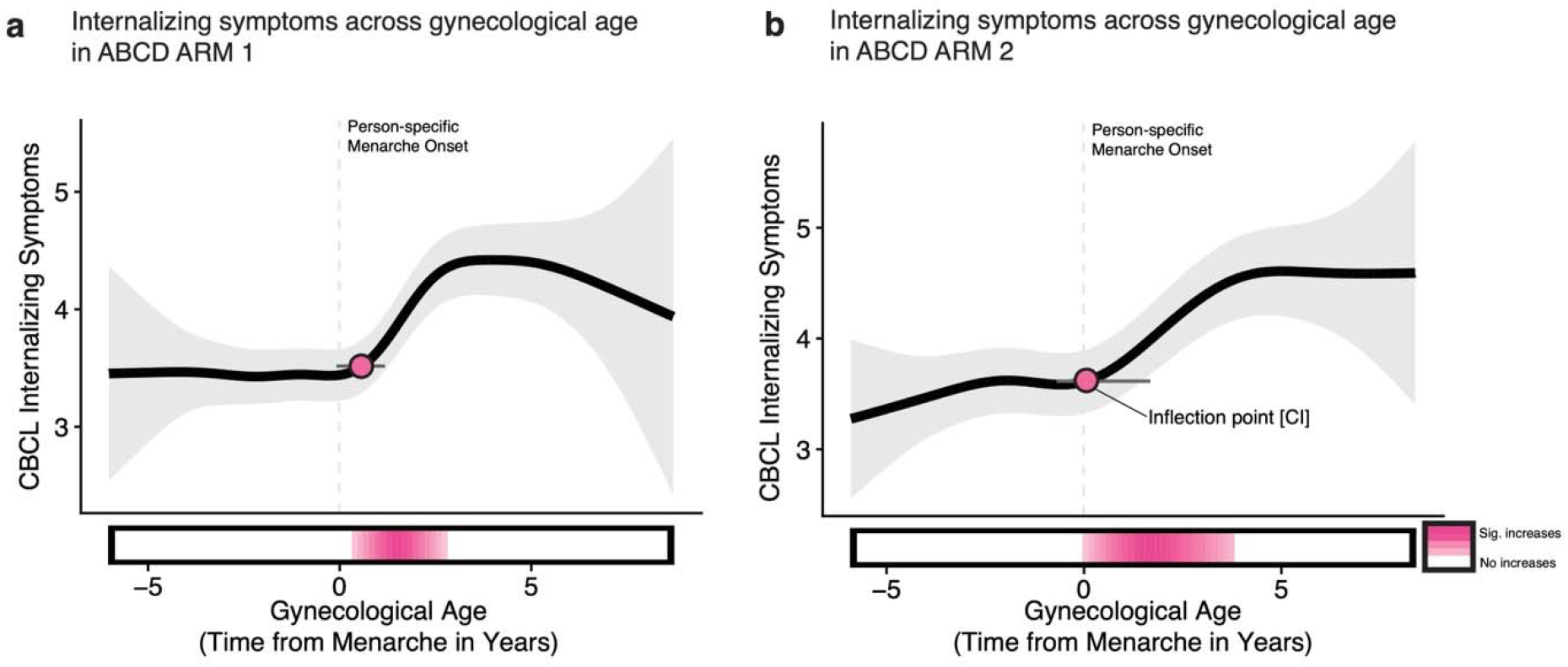
Menarche onset marks an inflection point in internalizing symptoms: Replication in ABCD ARMS matched samples. **a**, ARMS matched sample 1 (*n* = 2,450). **b**, ARMS matched sample 2 (*n* = 2,395). For **a** and **b**, Black line depicts predicted values from generalized additive mixed models (GAMMs) relating gynecological age (time relative to menarche onset) to Child Behavior Checklist (CBCL) internalizing symptoms in both ARMS matched sample 1 and 2. Light gray shaded regions represent 95% credible intervals (CI). The inflection points, derived from second derivative maximum indicating the point of fastest acceleration, are shown as pink points with a black outline. The gray horizontal lines represent the 95% CI of the inflection points derived from posterior sampling. The dashed vertical lines mark person-specific menarche onset (gynecological age = 0). The colored horizontal bars along the x-axis denote periods of significant change derived from first derivative estimates: pink indicates significant increases, white indicates no significant change. The consistent location of inflection points near menarche onset across both matched samples confirms that the acceleration in internalizing symptoms is temporally anchored to menarche.

**Extended Data Fig. 5.**
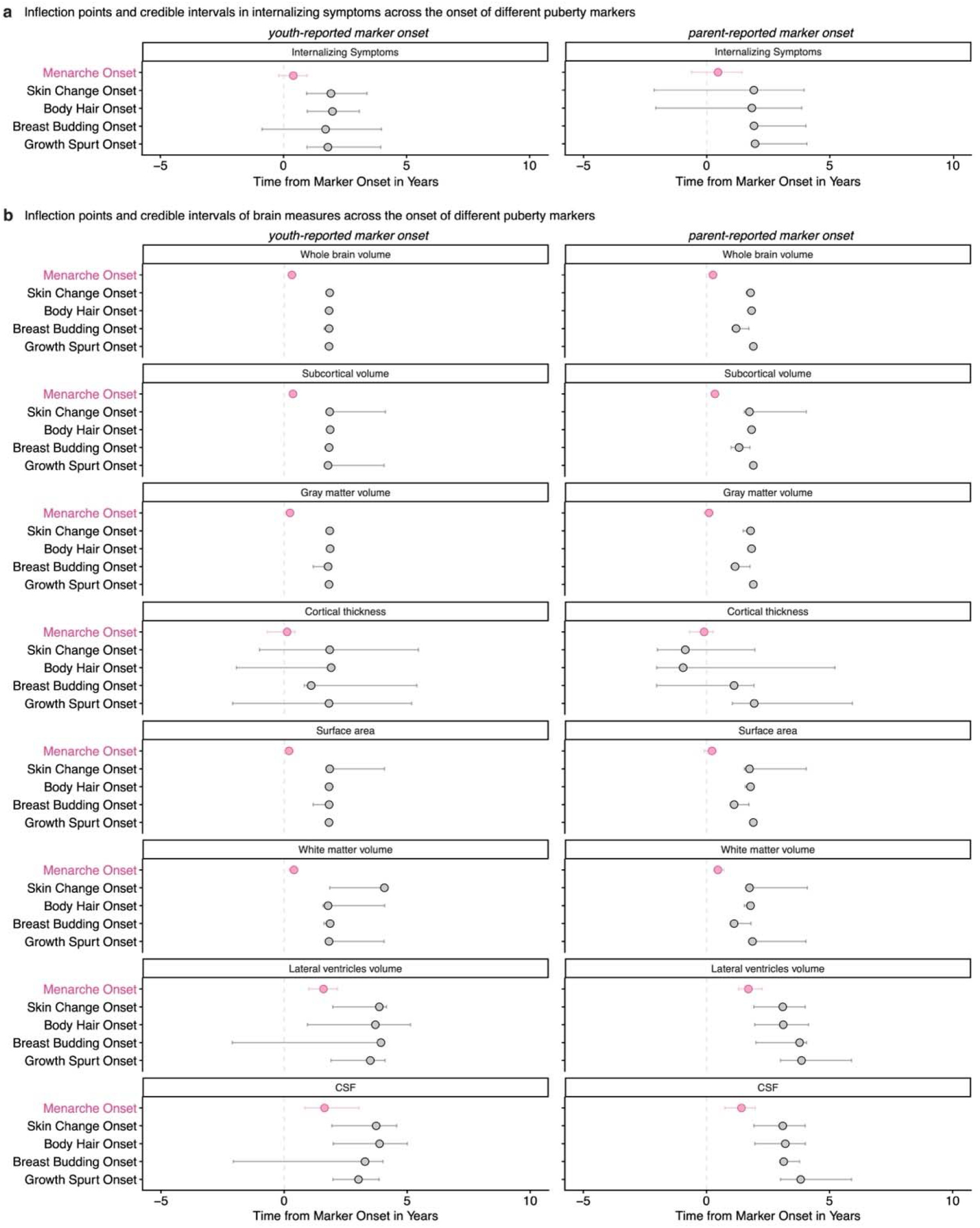
Inflection points for alterative puberty markers do not align with marker onset. **a**, Inflection points, derived from second derivate maxima, for Child Behavior Checklist (CBCL) internalizing symptoms are shown (*n* = 5,016). **b**, Inflection points, derived from second derivative minima, for structural brain measures are shown (*n* = 4,978). For **a** and **b**, Inflection points derived from second derivative extrema from generalized additive mixed models (GAMMs) with time relative to puberty marker onset based both on youth- and parent-report as the predictor. Gray points indicate inflection points for each alternative puberty marker (growth spurt, breast development, body hair, skin changes), pink points indicate inflection points for menarche, with horizontal lines representing 95% credible intervals. Dashed vertical line marks puberty marker and menarche onset. Results demonstrate that while menarche onset consistently aligns with inflection points in both internalizing symptoms and brain development, inflection points for alternative puberty markers occur well after their respective onset times, indicating that menarche uniquely serves as a milestone for adolescent neurodevelopment and mental health.

**Extended Data Fig. 6.**
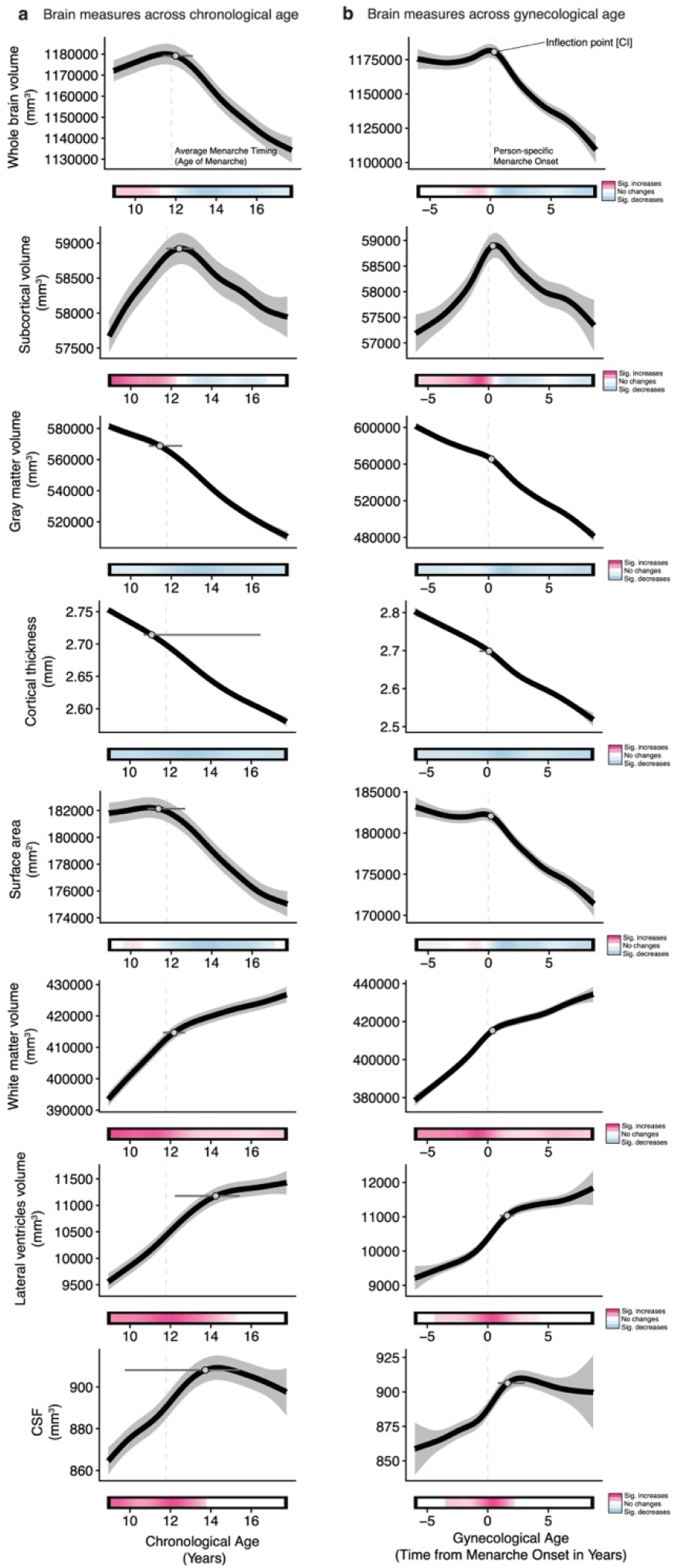
Morphometric brain measures across chronological and gynecological age. **a**, Black lines depict predicted values from generalized additive mixed models (GAMMs) relating chronological age to structural brain measures. **b**, Black lines depict predicted values from GAMMs relating gynecological age to structural brain measures. For **a** and **b**, Inflection points, derived from second derivative minima, are shown as gray points with black outlines, with 95% credible intervals (CI) derived from posterior sampling depicted as gray horizontal lines. For trajectories with decreasing slopes (e.g., declining gray matter volume), the minimum of the second derivative indicates the point of greatest acceleration in decline (steepest decrease in slope). For trajectories with increasing slopes (e.g., e.g., increasing white matter volume), the minimum indicates the point of greatest deceleration (where the rate of increase slows most). Note the location (close to menarche onset for brain tissue metrics) and increased temporal precision of the inflection points (narrower 95% CI) observed for gynecological age. Colored horizontal bars along the x-axis denote periods of significant change based on first derivative estimates: pink indicates significant increases, blue indicates significant decreases, white indicates no significant change. Dashed vertical lines mark average menarche timing (left panel) and person-specific menarche onset (right panel, time = 0). Gross volumetric, gray matter, and white matter metrics show inflection points aligned with menarche, whereas fluid-content metrics do not. CSF = cerebrospinal fluid. *n* = 4,978 female participants.

**Extended Data Fig. 7.**
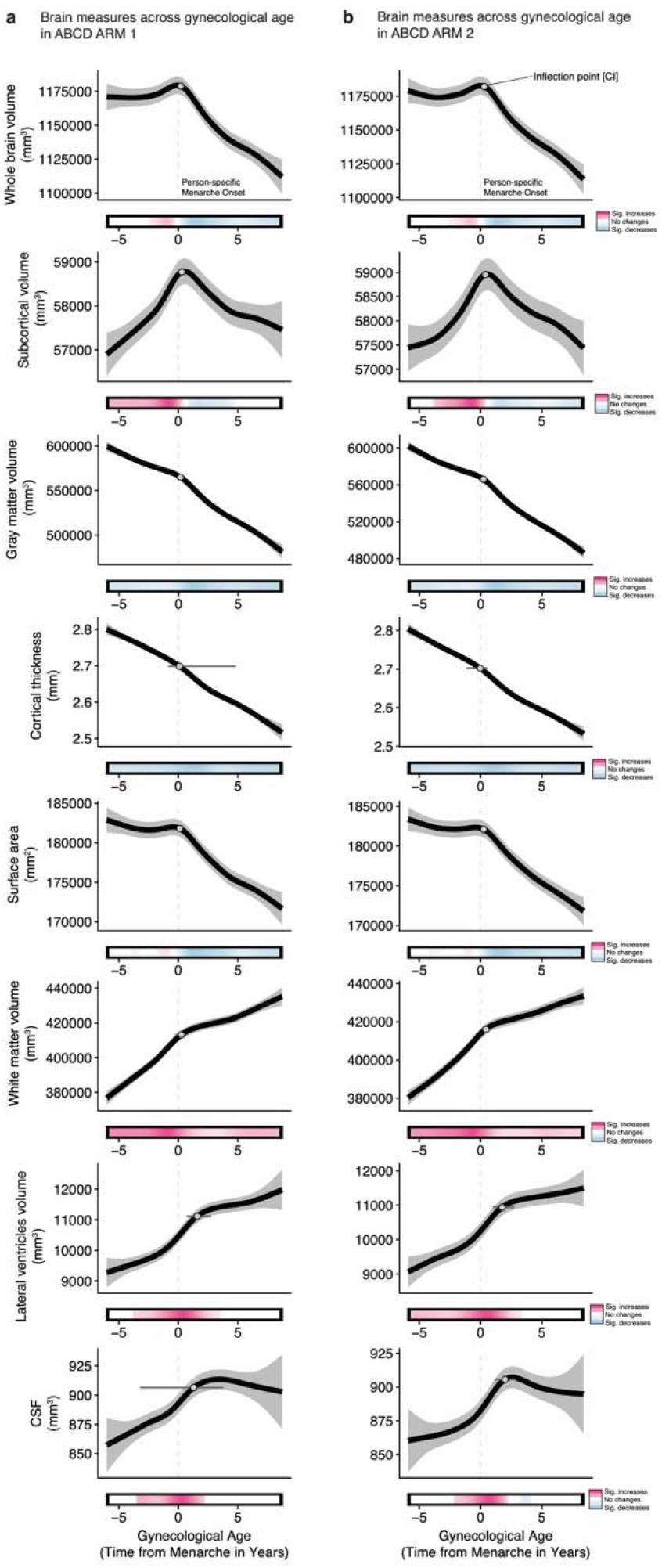
Menarche onset marks an inflection point in structural brain development: Replication in ABCD ARMS matched samples. **a**, ARMS matched sample 1 (*n* = 2,450). **b**, ARMS matched sample 2 (*n* = 2,395). For **a** and **b**, Black line depicts predicted values from generalized additive mixed models (GAMMs) relating gynecological age (time relative to menarche onset) to structural brain measures in both ARMS matched sample 1 and 2. Light gray shaded regions represent 95% credible intervals (CI). The inflection points, derived from second derivative minima, are shown as a gray point with a black outline. For trajectories with decreasing slopes (e.g., declining gray matter volume), the minimum of the second derivative indicates the point of greatest acceleration in decline (steepest decrease in slope). For trajectories with increasing slopes (e.g., increasing white matter volume), the minimum indicates the point of greatest deceleration (where the rate of increase slows most). The gray horizontal lines represent the 95% CI of the inflection points derived from posterior sampling. The dashed vertical lines mark person-specific menarche onset (gynecological age = 0). The colored horizontal bars along the x-axis denote the period of significant change derived from first derivative estimates: Pink indicates significant increases, blue indicates significant decreases, white indicates no significant change. The consistent location of inflection points near menarche onset across both matched samples confirms that inflection points in structural brain development are temporally anchored to menarche.

**Extended Data Fig. 8.**
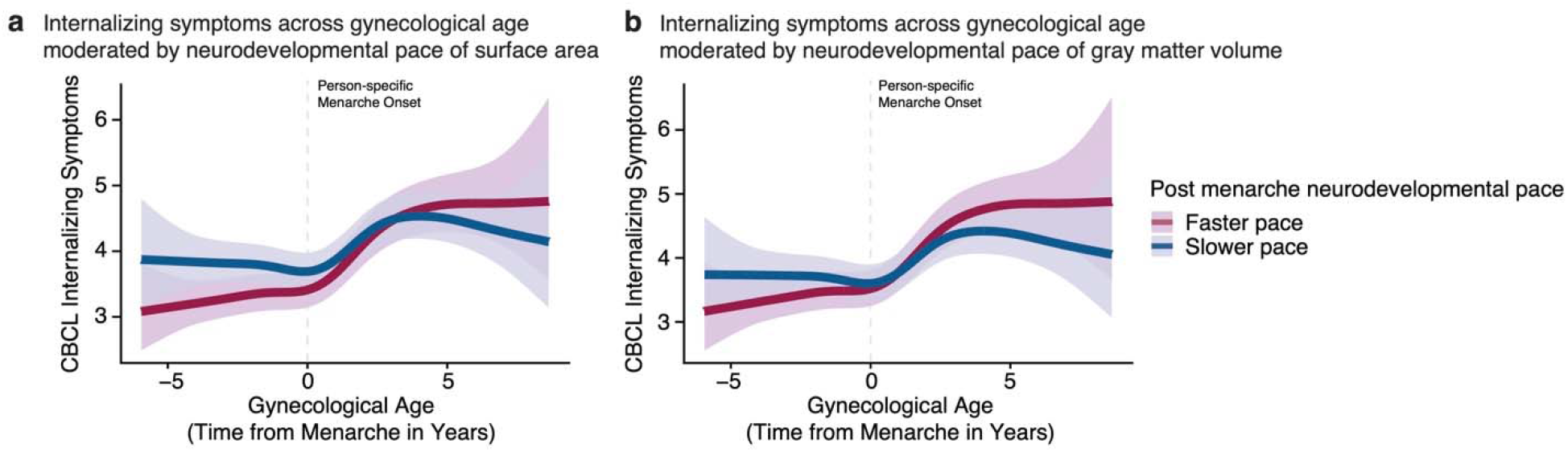
Individual post-menarche brain changes moderate the relationship between gynecological age and internalizing symptoms. **a**, Faster post-menarche decline in surface area predicts greater internalizing symptom severity. **b**, Faster post-menarche decline in gray matter volume predicts greater internalizing symptom severity. For **a** and **b**, Colored lines depict predicted values from generalized additive mixed models (GAMMs) relating gynecological to internalizing symptoms, stratified by post-menarche developmental pace (faster vs. slower). Colored shaded regions represent 95% credible intervals (CI). Dashed vertical lines mark person-specific menarche onset (gynecological age = 0). Adolescents experiencing faster brain structural decline following menarche show steeper increases in internalizing symptoms. *n* = 4, 978 female participants.

**Extended Data Fig. 9.**
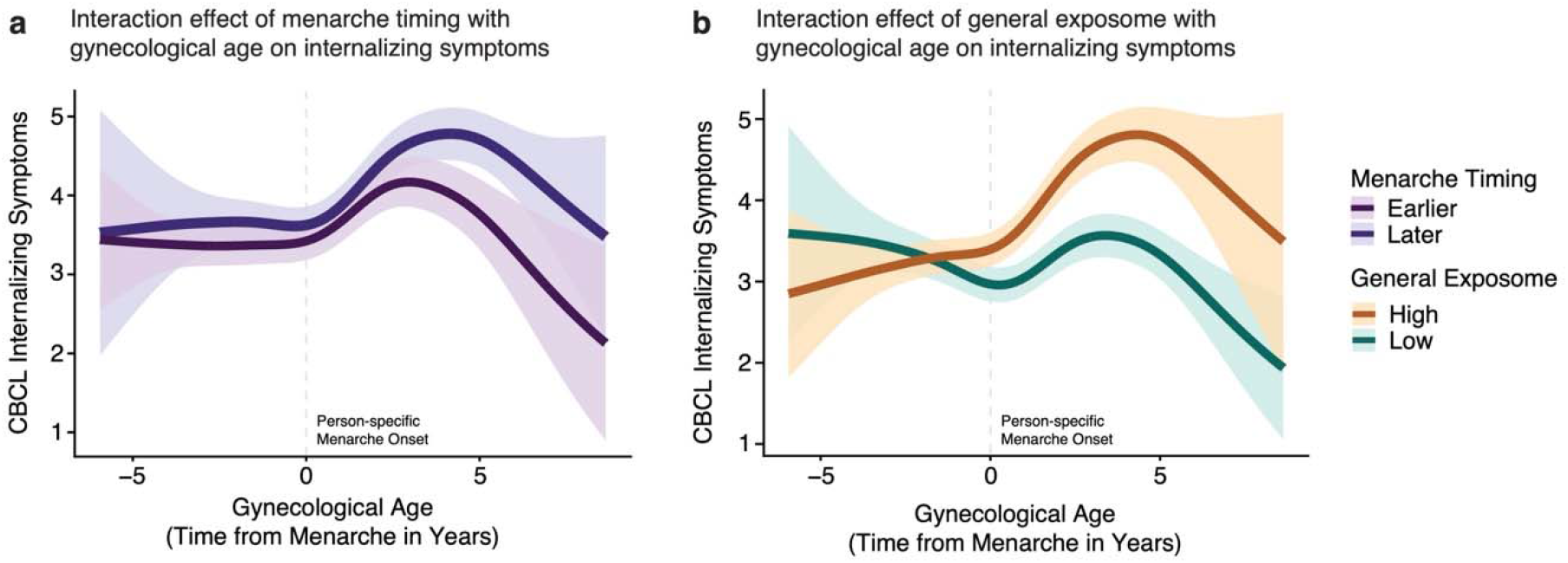
Menarche timing and general exposome moderate internalizing symptom severity. **a**, While internalizing symptoms increase following menarche onset across both early and late menarche timing, early menarche timing is associated with higher symptom levels throughout the post-menarche period. **b**, While internalizing symptoms increase following menarche onset across both high and low general exposome groups, higher general exposome is associated with higher symptom levels throughout the post-menarche period. For **a** and **b**, Colored lines depict predicted values from generalized additive mixed models (GAMMs) relating gynecological age to internalizing symptoms, stratified by menarche timing (early vs. late) and general exposome level (low vs. high), respectively. Shaded regions represent 95% credible intervals. Dashed vertical line marks person-specific menarche onset (gynecological age = 0). These results indicate that while the timing and rate of symptom increase relative to menarche onset are consistent across individuals, overall symptom severity differs based on pubertal timing and environmental context. *n* = 5,016 female participants.

**Extended Data Fig. 10.**
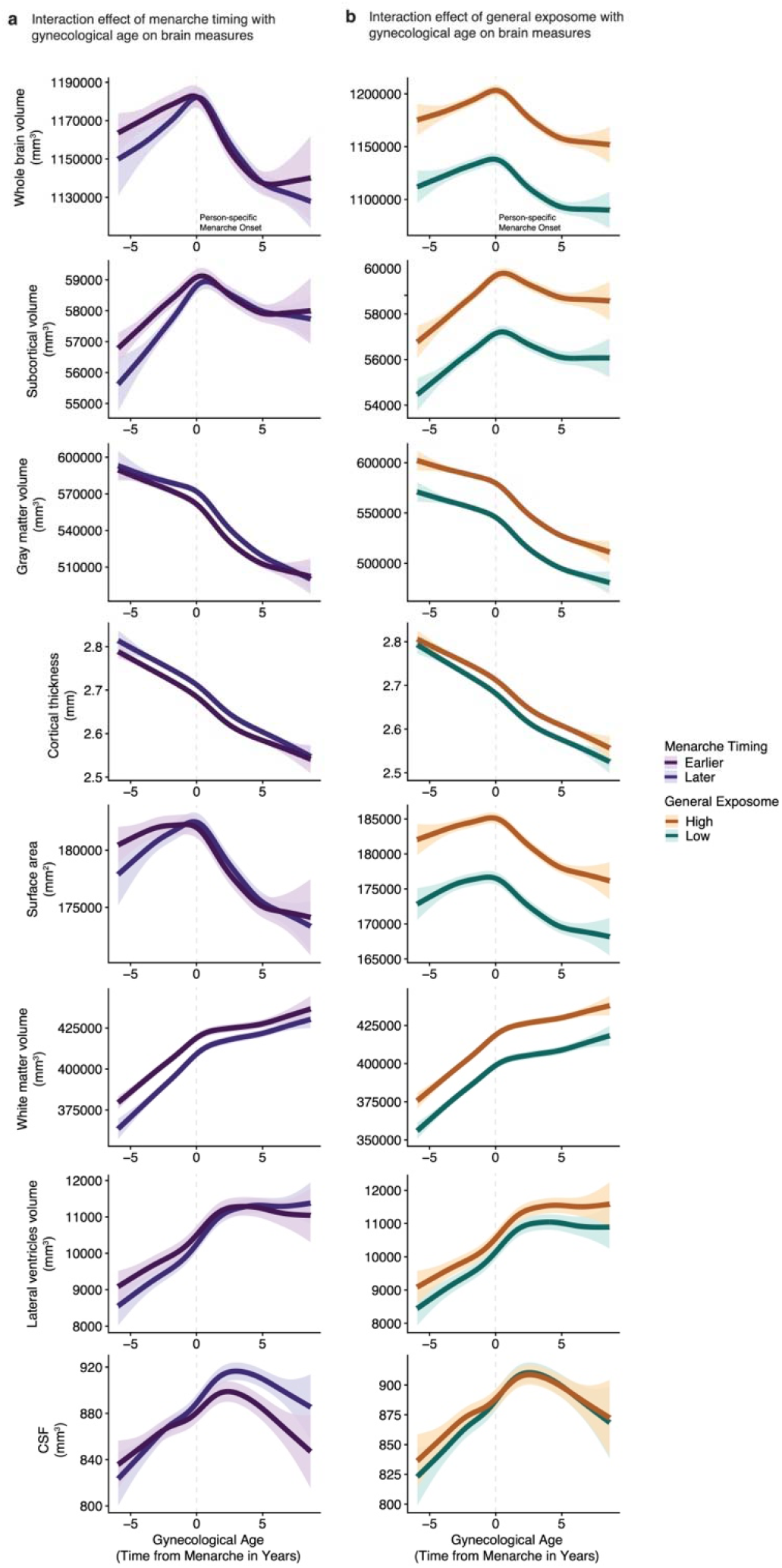
Menarche timing and general exposome moderate structural brain measures. **a**, Menarche timing interacted significantly with menarche timing for all brain measures except cortical thickness. Inflection points around menarche onset were observed across all measures. For whole brain volume, subcortical volume, and surface area, early maturers showed lager volumes pre-menarche, with similar post-menarche decline across timing groups. Gray matter volume was consistently higher in late maturers throughout development, while white matter volume was consistently lower in late maturers. For cerebrospinal fluid (CSF) and lateral ventricles volume, late maturers showed lower volumes pre-menarche compared to early maturers, with a reversal of this pattern post-menarche. **b**, General exposome interacted significantly with gynecological age for all brain measures except CSF and lateral ventricle volumes. Inflection points around menarche onset were observed across all measures. Across all significant measures, higher general exposome (lower environmental adversity) was associated with greater brain volumes and cortical thickness. For **a** and **b**, Colored lines depict predicted values from generalized additive mixed models (GAMMs) relating gynecological age to brain measures, stratified by menarche timing (early vs. late) and general exposome level (low vs. high), respectively. Shaded regions represent 95% credible intervals. Dashed vertical line marks person-specific menarche onset (gynecological age = 0). *n* = 4,978 female participants.

