## Extended Data Figures for "Menarche onset is an inflection point for mental health and brain development"

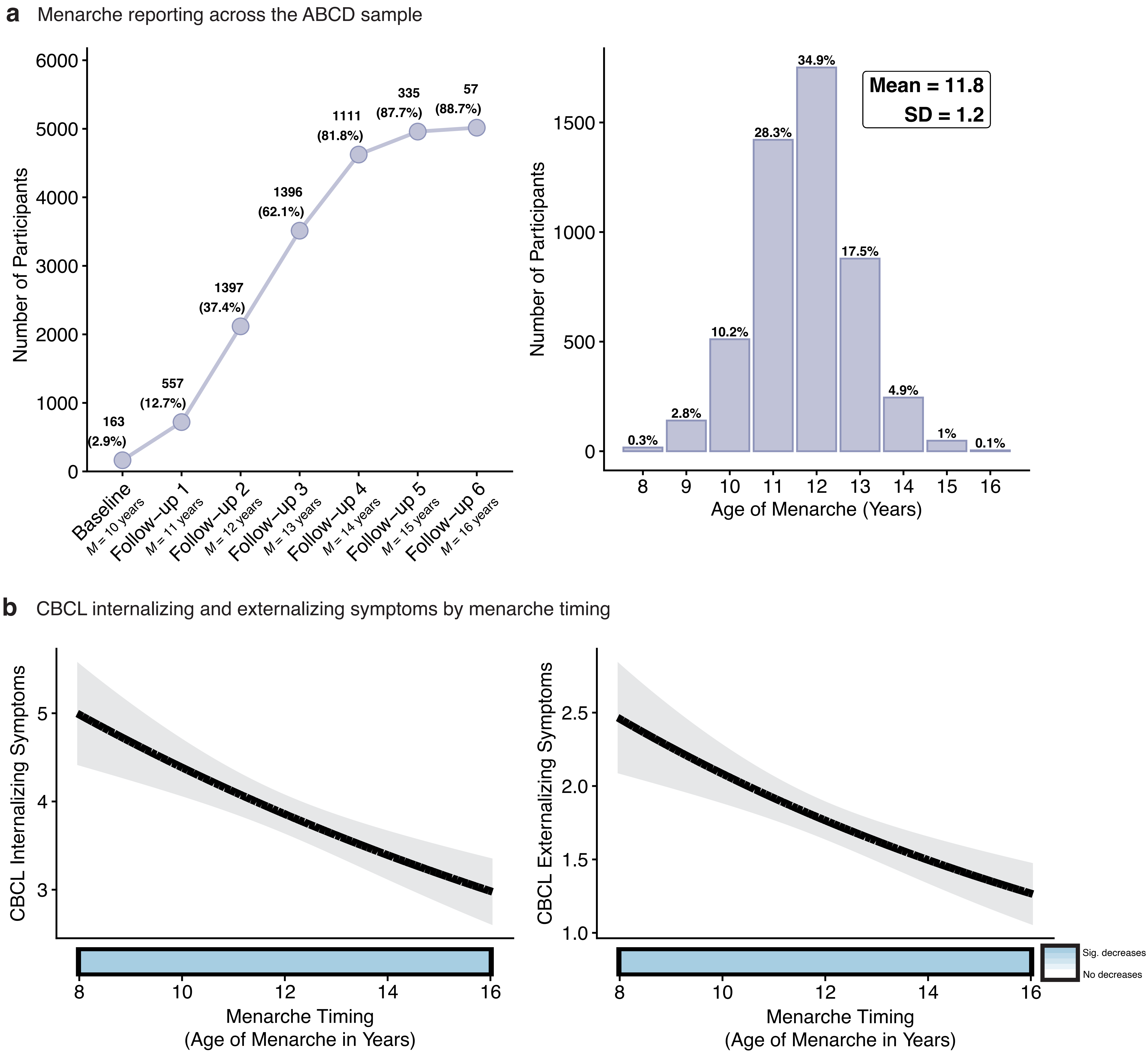

a Menarche reporting across the ABCD sample
Number of Participants
0
1000
2000
3000
4000
5000
6000
163 (2.9%)
557 (12.7%)
1397 (37.4%)
1396 (62.1%)
1111 (81.8%)
335 (87.7%)
57 (88.7%)
Baseline M = 10 years
Follow–up 1 M = 11 years
Follow–up 2 M = 12 years
Follow–up 3 M = 13 years
Follow–up 4 M = 14 years
Follow–up 5 M = 15 years
Follow–up 6 M = 16 years
Number of Participants
0
500
1000
1500
0.3%
2.8%
10.2%
28.3%
34.9%
17.5%
4.9%
1%
0.1%
Mean = 11.8
SD = 1.2
8 9 10 11 12 13 14 15 16
Age of Menarche (Years)
b CBCL internalizing and externalizing symptoms by menarche timing
CBCL Internalizing Symptoms
3
4
5
8 10 12 14 16
Menarche Timing
(Age of Menarche in Years)
CBCL Externalizing Symptoms
1.0
1.5
2.0
2.5
8 10 12 14 16
Menarche Timing
(Age of Menarche in Years)
Sig. decreases
No decreases

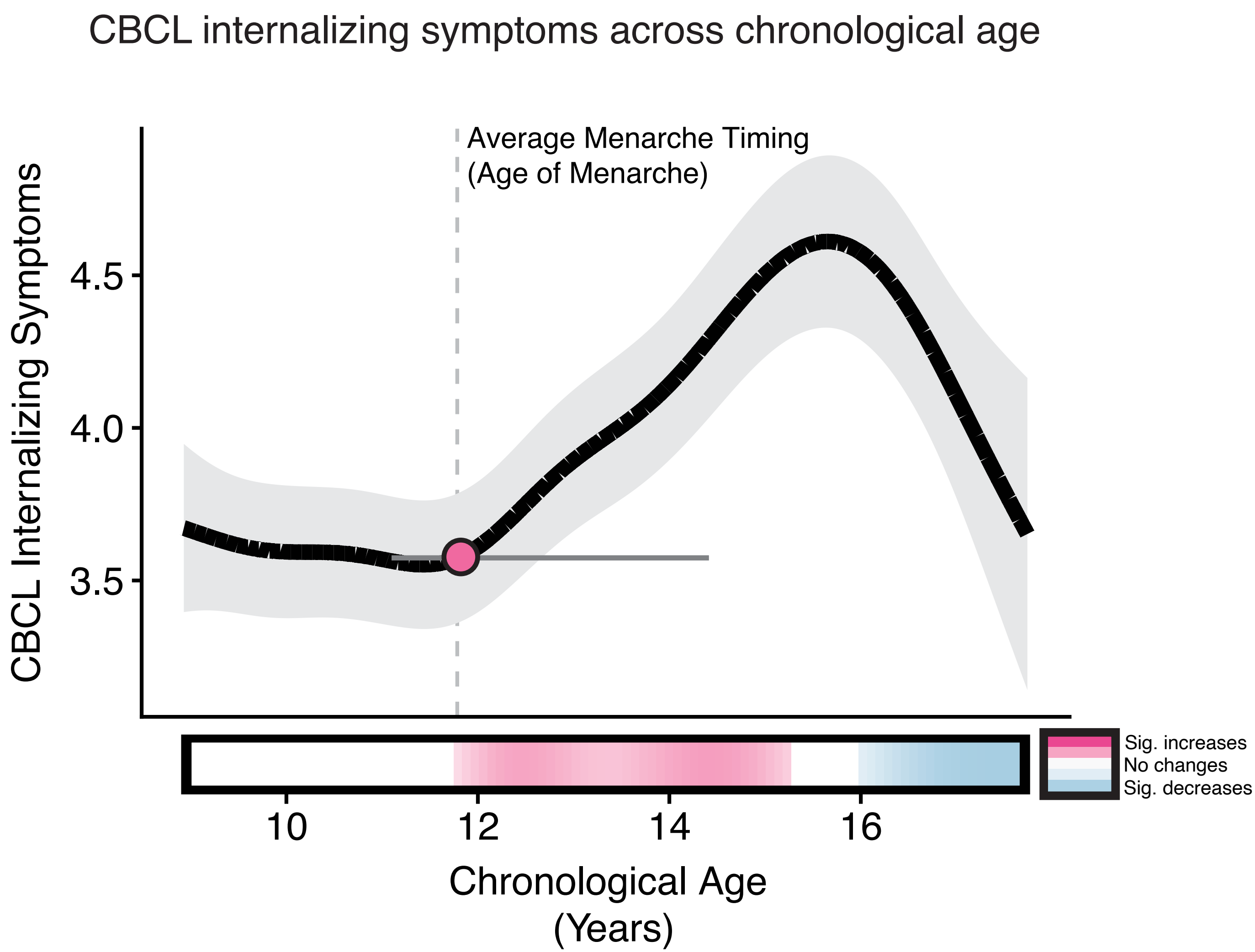

CBCL internalizing symptoms across chronological age
Average Menarche Timing
(Age of Menarche)
CBCL Internalizing Symptoms
4.5
4.0
3.5
10
12
14
16
Chronological Age
(Years)
Sig. increases
No changes
Sig. decreases

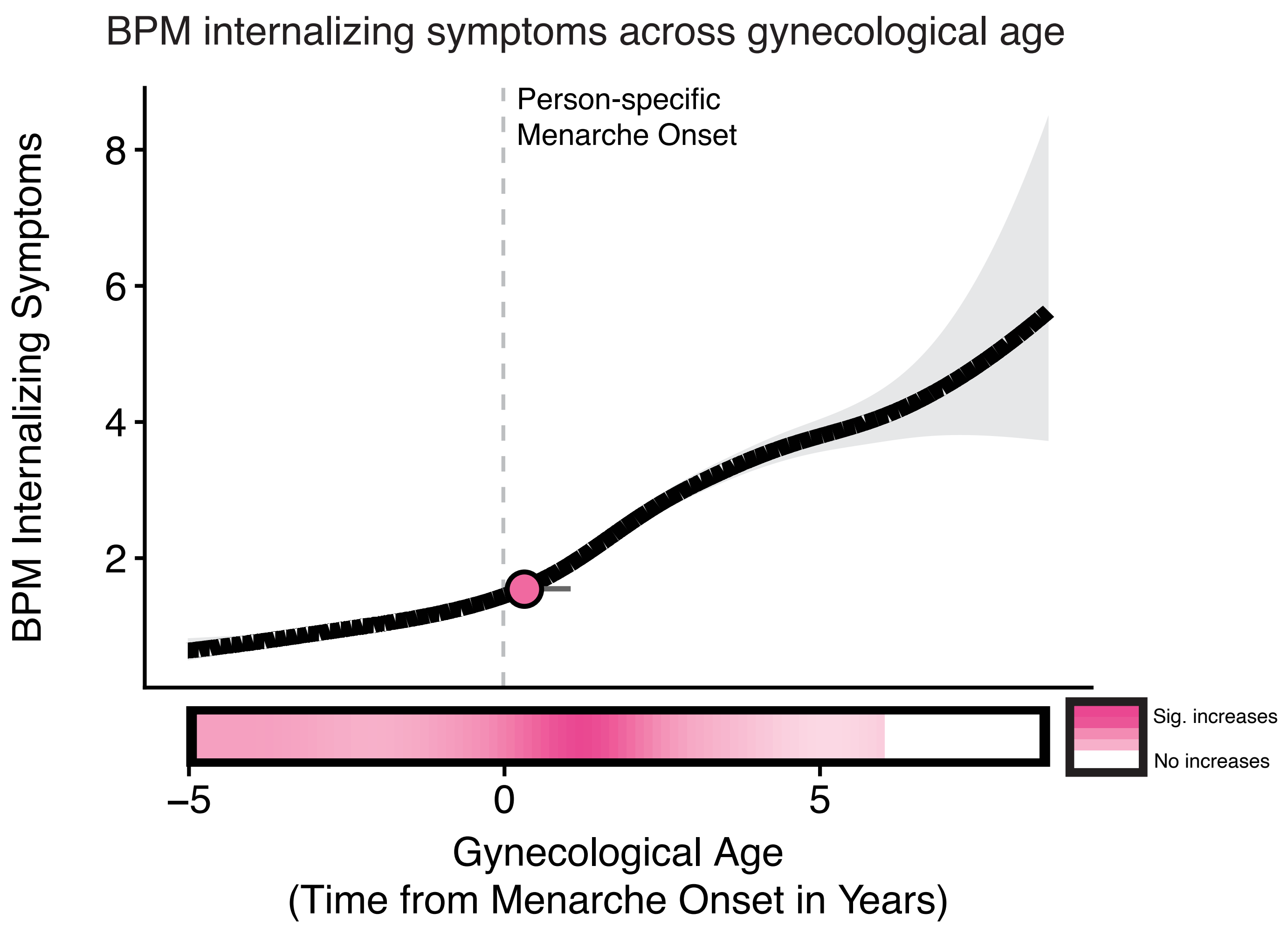

BPM internalizing symptoms across gynecological age
Person-specific
Menarche Onset
BPM Internalizing Symptoms
8
6
4
2
–5
0
5
Sig. increases
No increases
Gynecological Age
(Time from Menarche Onset in Years)

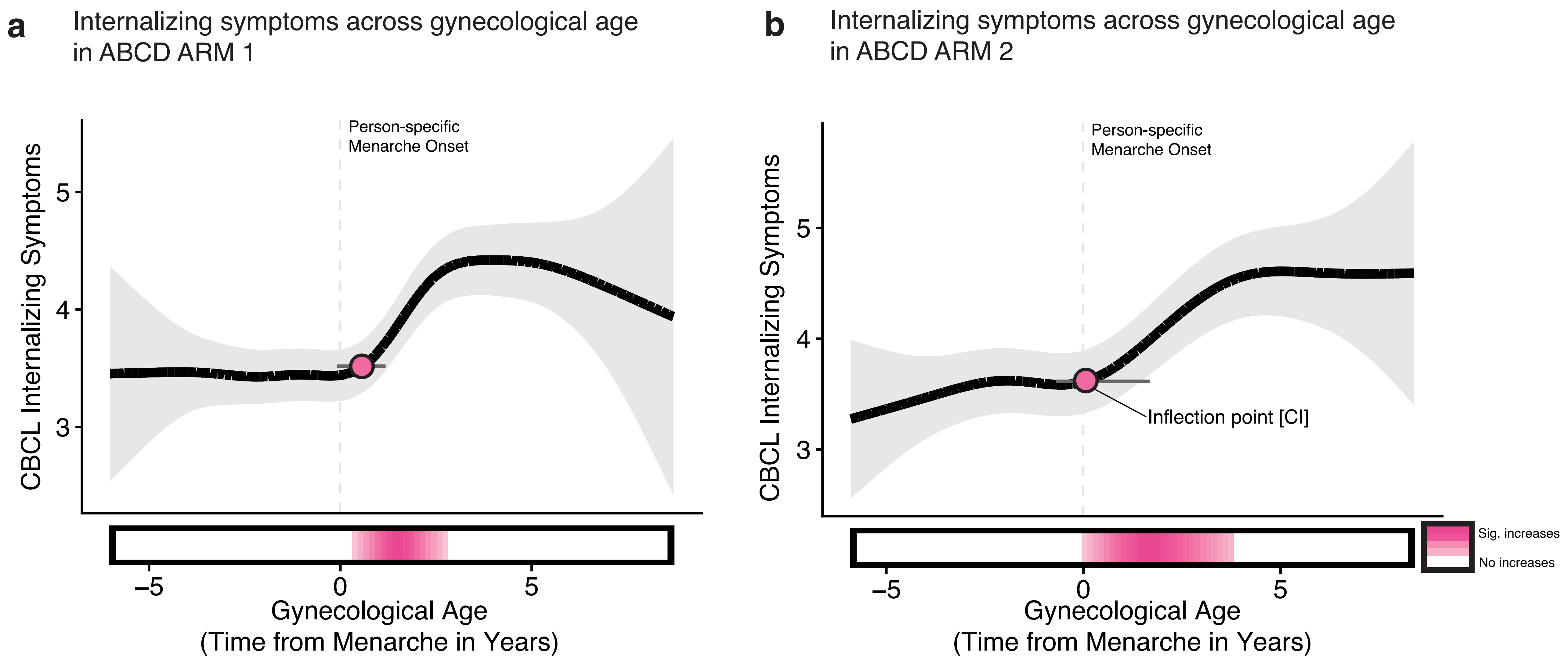
a
Internalizing symptoms across gynecological age
in ABCD ARM 1
Person-specific
Menarche Onset
CBCL Internalizing Symptoms
5
4
3
−5
0
5
Gynecological Age
(Time from Menarche in Years)
b
Internalizing symptoms across gynecological age
in ABCD ARM 2
Person-specific
Menarche Onset
Inflection point [CI]
CBCL Internalizing Symptoms
5
4
3
−5
0
5
Gynecological Age
(Time from Menarche in Years)
Sig. increases
No increases

**a** Inflection points and credible intervals in internalizing symptoms across the onset of different puberty markers

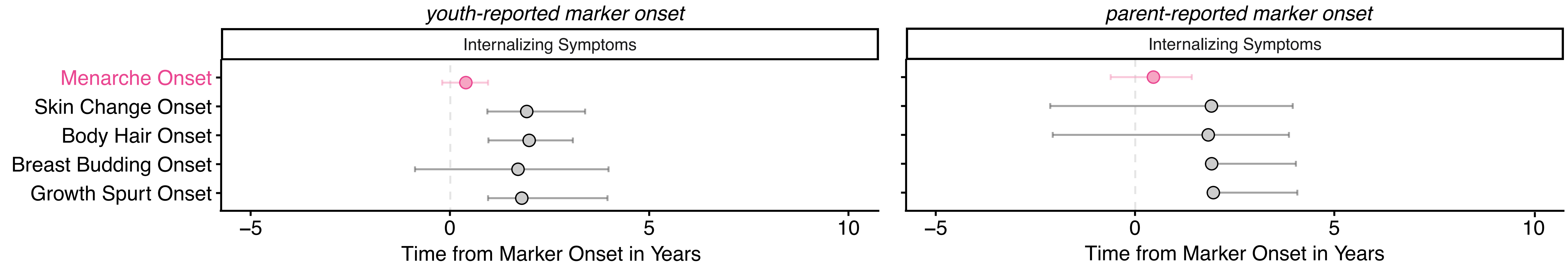


**b** Inflection points and credible intervals of brain measures across the onset of different puberty markers

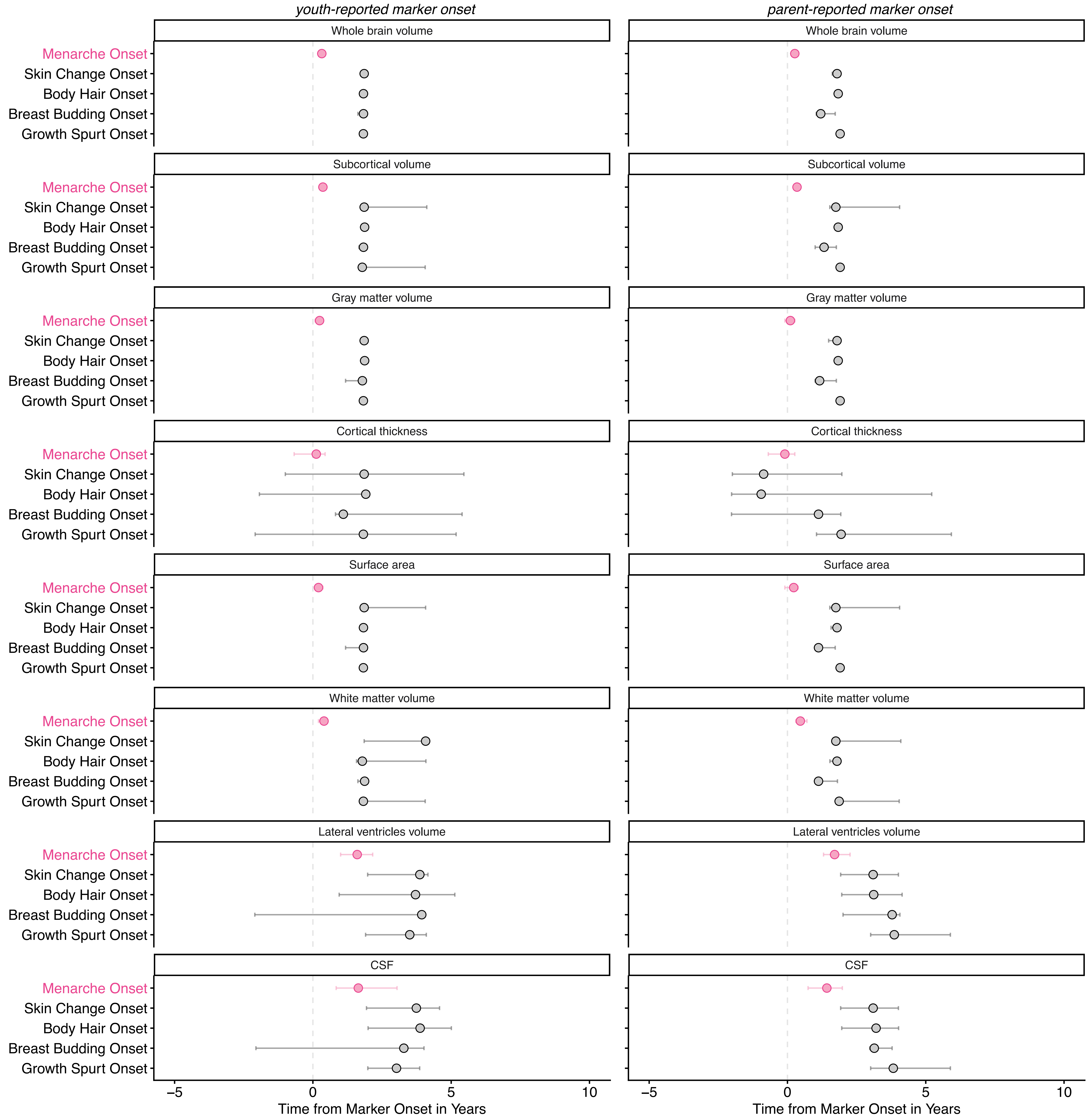

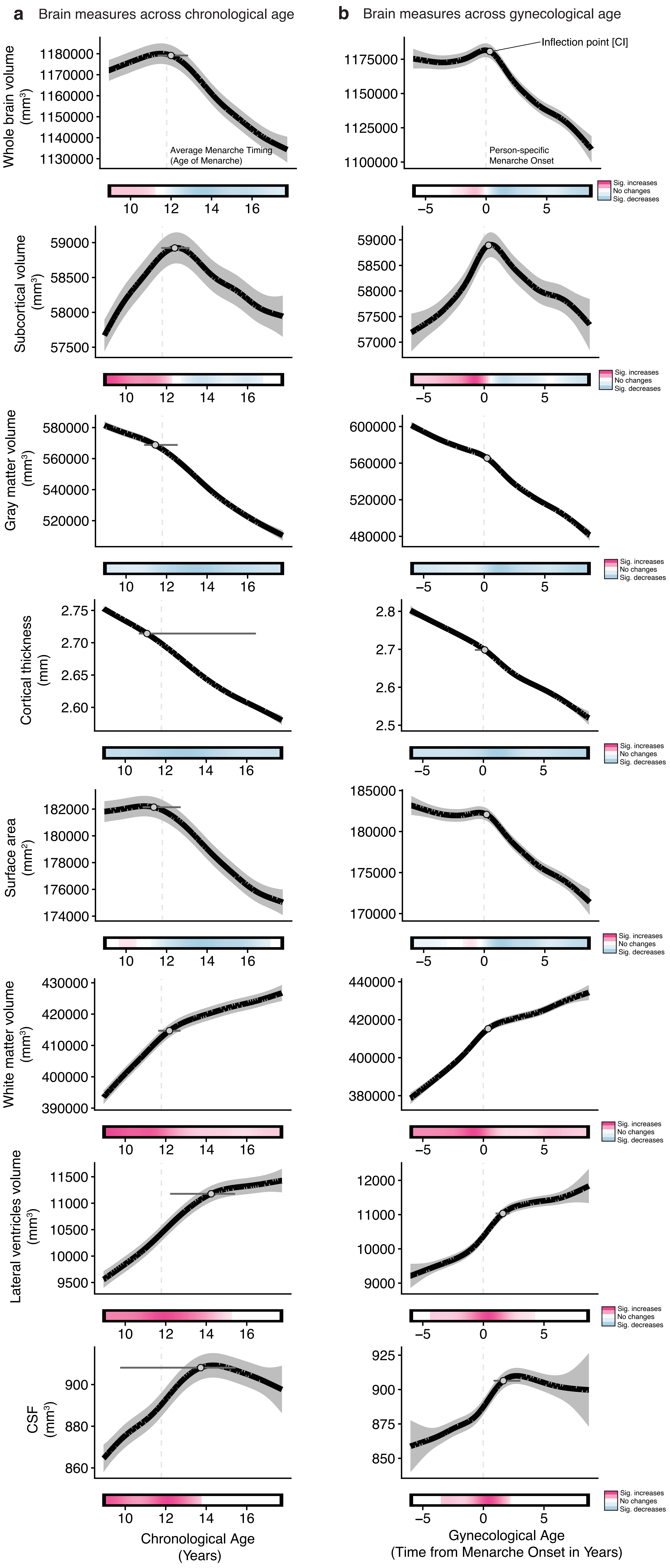

a
Brain measures across chronological age
b
Brain measures across gynecological age
Inflection point [CI]
Average Menarche Timing (Age of Menarche)
Person-specific Menarche Onset
Sig. increases
No changes
Sig. decreases
Whole brain volume (mm³)
Subcortical volume (mm³)
Gray matter volume (mm³)
Cortical thickness (mm)
Surface area (mm²)
White matter volume (mm³)
Lateral ventricles volume (mm³)
CSF (mm³)
Chronological Age (Years)
Gynecological Age (Time from Menarche Onset in Years)

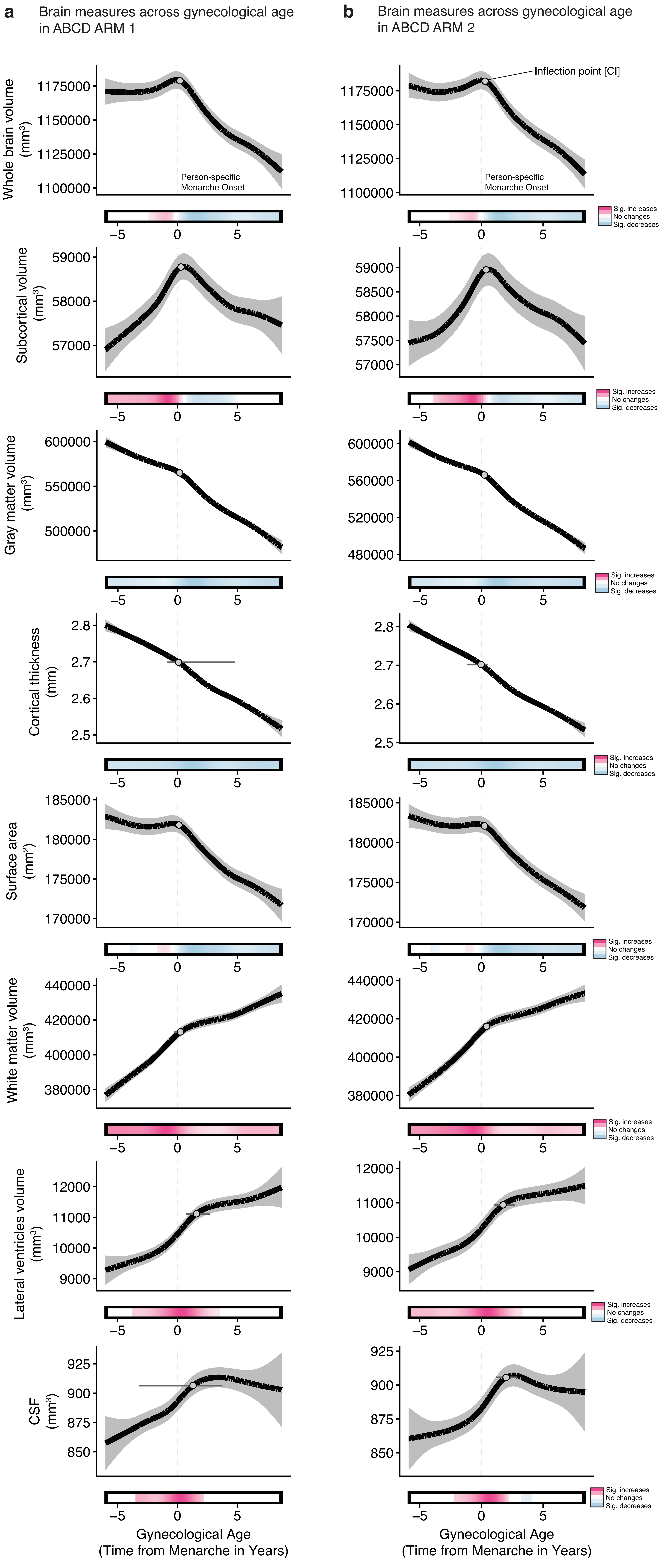
a
Brain measures across gynecological age in ABCD ARM 1
b
Brain measures across gynecological age in ABCD ARM 2
Inflection point [CI]
Person-specific Menarche Onset
Whole brain volume (mm³)
Subcortical volume (mm³)
Gray matter volume (mm³)
Cortical thickness (mm)
Surface area (mm²)
White matter volume (mm³)
Lateral ventricles volume (mm³)
CSF (mm³)
Sig. increases
No changes
Sig. decreases
Gynecological Age (Time from Menarche in Years)

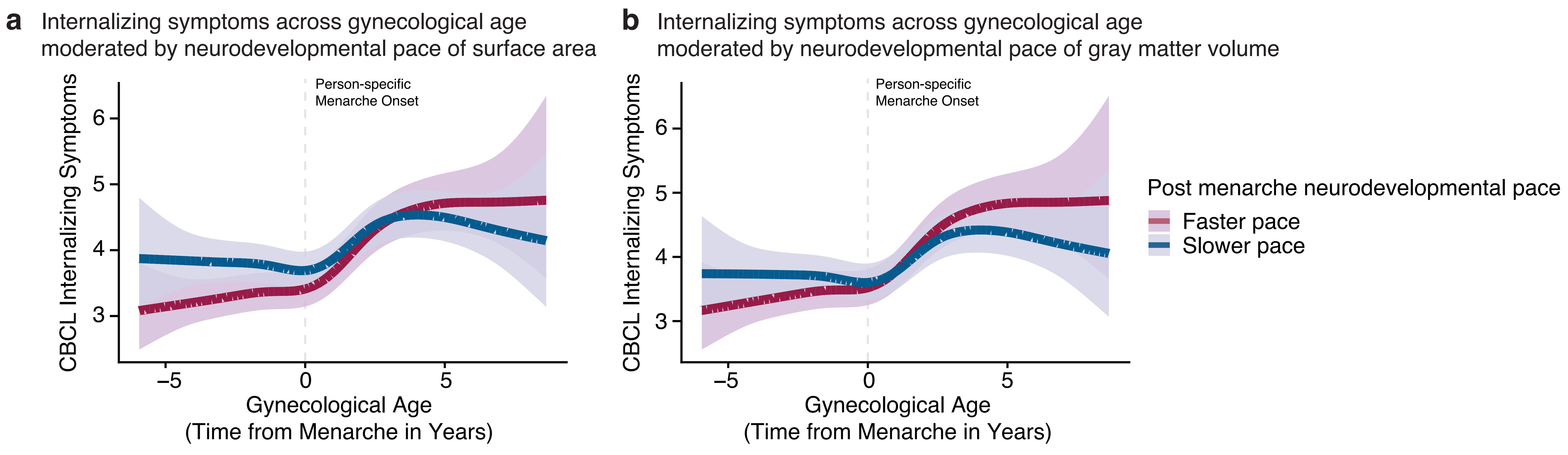
a
Internalizing symptoms across gynecological age
moderated by neurodevelopmental pace of surface area
b
Internalizing symptoms across gynecological age
moderated by neurodevelopmental pace of gray matter volume
Person-specific
Menarche Onset
CBCL Internalizing Symptoms
Gynecological Age
(Time from Menarche in Years)
Post menarche neurodevelopmental pace
Faster pace
Slower pace

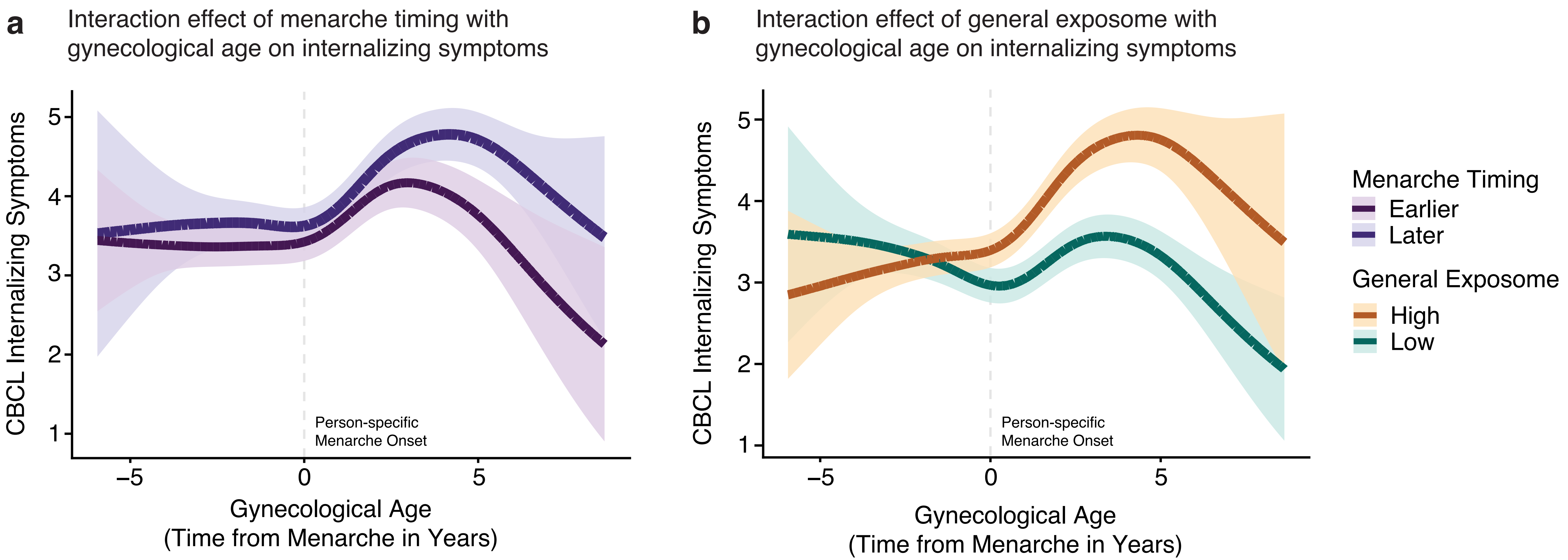

a
Interaction effect of menarche timing with gynecological age on internalizing symptoms
CBCL Internalizing Symptoms
5
4
3
2
1
Person-specific
Menarche Onset
−5
0
5
Gynecological Age
(Time from Menarche in Years)
b
Interaction effect of general exposome with gynecological age on internalizing symptoms
CBCL Internalizing Symptoms
Gynecological Age
(Time from Menarche in Years)
Menarche Timing
Earlier
Later
General Exposome
High
Low

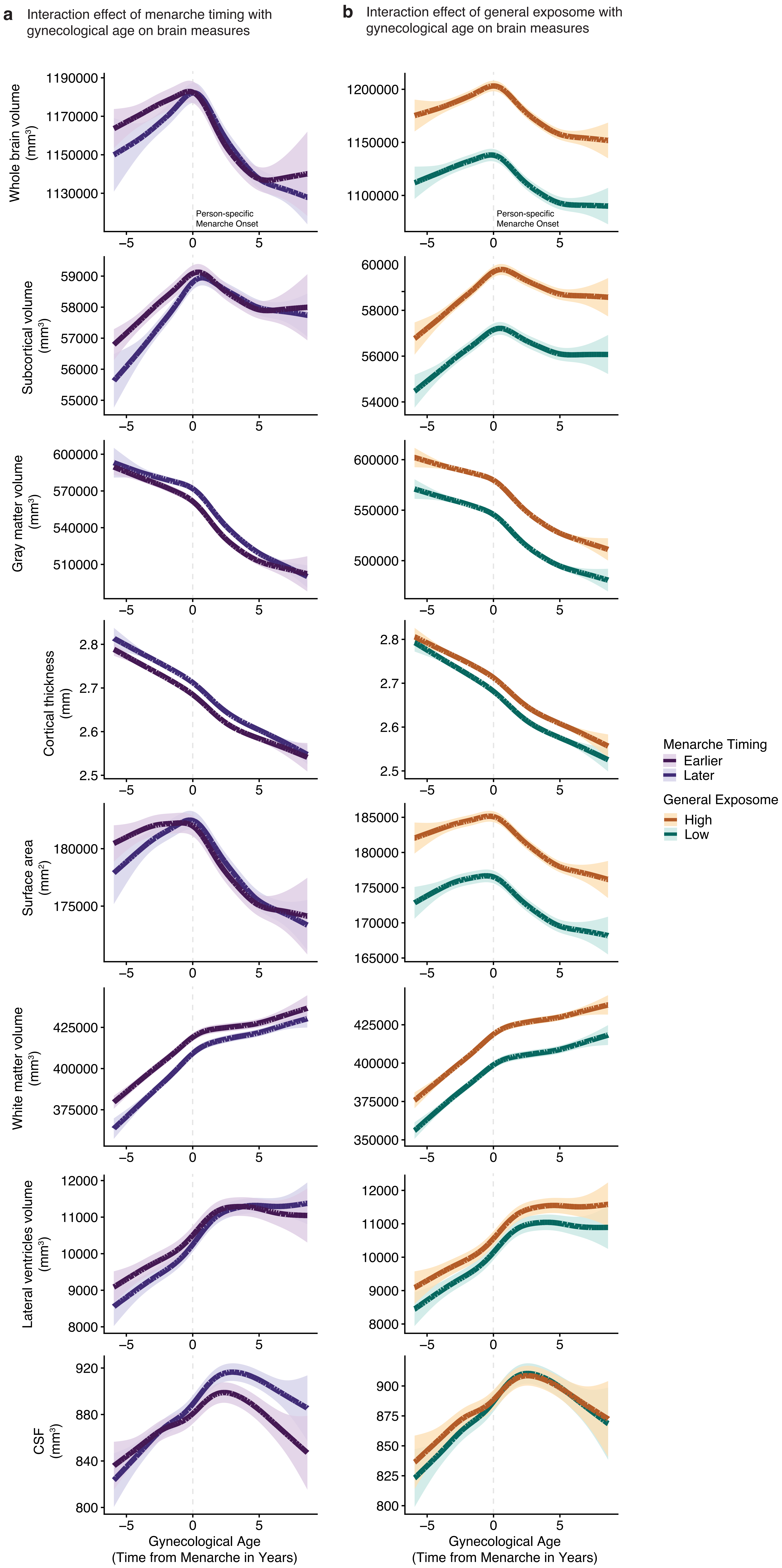
a
Interaction effect of menarche timing with gynecological age on brain measures
b
Interaction effect of general exposome with gynecological age on brain measures
Whole brain volume (mm³)
Subcortical volume (mm³)
Gray matter volume (mm³)
Cortical thickness (mm)
Surface area (mm²)
White matter volume (mm³)
Lateral ventricles volume (mm³)
CSF (mm³)
Person-specific Menarche Onset
Gynecological Age (Time from Menarche in Years)
Menarche Timing
Earlier
Later
General Exposome
High
Low
