## Supplementary Material for "Menarche onset is an inflection point for mental health and brain development"

| **Supplementary Table 1. Results from Generalized Additive Mixed Models (GAMMs) for internalizing and externalizing symptoms using the Child Behavior Checklist.** | | | | | |
| --- | --- | --- | --- | --- | --- |
| **Outcome** | **Term** | **edf** | ***F*** | ***p*** | ***p*_Bonferroni_** |
| Internalizing symptoms | s(Chronological Age) | 6.48 | 63.39 | **<.001** | **<.001** |
|  | s(General Exposome) | 4.501 | 7.482 | **<.001** | **<.001** |
|  |  |  |  |  | *R*^2^ = 0.01 |
|  | s(Menarche Timing) | 1 | 23.22 | **<.001** | **<.001** |
|  | s(General Exposome) | 4.682 | 10.1 | **<.001** | **<.001** |
|  |  |  |  |  | *R*^2^ = 0.006 |
|  | s(Gynecological Age) | 6.042 | 69.45 | **<.001** | **<.001** |
|  | s(Menarche Timing) | 1 | 6.791 | **0.009** | 0.128 |
|  | s(General Exposome) | 4.694 | 9.532 | **<.001** | **<.001** |
|  |  |  |  |  | *R*^2^ = 0.014 |
| Externalizing symptoms | s(Chronological Age) | 3.976 | 34.5 | **<.001** | **<.001** |
|  | s(General Exposome) | 4.208 | 30.08 | **<.001** | **<.001** |
|  |  |  |  |  | *R*^2^ = 0.019 |
|  | s(Menarche Timing) | 1 | 21.95 | **<.001** | **<.001** |
|  | s(General Exposome) | 4.337 | 23.75 | **<.001** | **<.001** |
|  |  |  |  |  | *R*^2^ = 0.019 |
|  | s(Gynecological Age) | 3.735 | 35.01 | **<.001** | **<.001** |
|  | s(Menarche Timing) | 1 | 39.27 | **<.001** | **<.001** |
|  | s(General Exposome) | 4.339 | 23.08 | **<.001** | **<.001** |
|  |  |  |  |  | *R*^2^ = 0.02 |
| Note. The term s() represents a smooth function of the predictor, modeling potential nonlinear fluctuations over time. The effective degrees of freedom (edf) indicate the complexity of the smooth term, with higher values suggesting greater flexibility. *p*_Bonferroni_ = *p*-values after Bonferroni correction for multiple comparisons. Significant results are indicated in bold. Total number of observations, *n* = 29,837. | | | | | |

| **Supplementary Table 2. Second derivative minima and maxima for internalizing and externalizing symptoms using the Child Behavior Checklist** | | | |
| --- | --- | --- | --- |
| **Outcome** | **Predictor** | **2^nd^ derivative minimum [CI]** | **2^nd^ derivative maximum [CI]** |
| Internalizing symptoms | Chronological Age | 16.044 [15.026;16.044] | 11.810 [11.121;14.403] |
| Internalizing symptoms | Gynecological Age | 2.468 [-1.650;4.619] | 0.390 [-0.203;0.947] |
| Externalizing symptoms | Chronological Age | 15.748 [12.565;16.044] | 11.580 [9.775;13.516] |
| Externalizing symptoms | Gynecological Age | 2.393 [-2.169;4.619] | -0.426 [-2.763;0.761] |
| Note. 2^nd^ derivatives describe the rate of change in the slope of the trajectory and identify inflection points where the curvature changes. The second derivative maximum indicates the point of fastest acceleration (steepest increase in slope), occurring where symptoms are increasing most rapidly. The second derivative minimum indicates the point of maximum deceleration (greatest slowing of the slope), occurring where the rate of increase or decrease is slowing most. For chronological age, values represent age in years; for gynecological age, values represent time in years relative to individual menarche onset (negative values = before menarche onset, positive values = after menarche onset). Credible intervals [CI] were derived from posterior sampling of the fitted generalized additive mixed models (GAMMs). | | | |

| **Supplementary Table 3. Results from Generalized Additive Mixed Models (GAMMs) for internalizing using the Brief Problem Monitor.** | | | | |
| --- | --- | --- | --- | --- |
| **Outcome** | **Term** | **edf** | ***F*** | ***p*** |
| Internalizing symptoms | s(Gynecological Age) | 6.473 | 725.194 | **<.001** |
|  | s(Menarche Timing) | 1 | 25.837 | **<.001** |
|  | s(General Exposome) | 3.5 | 4.694 | **0.005** |
|  |  |  |  | *R*^2^ = 0.079 |
| Note. The term s() represents a smooth function of the predictor, modeling potential nonlinear fluctuations over time. The effective degrees of freedom (edf) indicate the complexity of the smooth term, with higher values suggesting greater flexibility. Significant results are indicated in bold. Total number of observations, *n* = 25,985. | | | | |

| **Supplementary Table 4. Second derivative minima and maxima for internalizing symptoms using the Brief Problem Monitor.** | | | |
| --- | --- | --- | --- |
| **Outcome** | **Predictor** | **2^nd^ derivative minimum [CI]** | **2^nd^ derivative maximum [CI]** |
| Internalizing symptoms | Gynecological Age | 2.144 [2.174;4.047] | 0.301 [0.120;1.056] |
| Note. 2^nd^ derivatives describe the rate of change in the slope of the trajectory and identify inflection points where the curvature changes. The second derivative maximum indicates the point of fastest acceleration (steepest increase in slope), occurring where symptoms are increasing most rapidly. The second derivative minimum indicates the point of maximum deceleration (greatest slowing of the slope), occurring where the rate of increase or decrease is slowing most. For chronological age, values represent age in years; for gynecological age, values represent time in years relative to individual menarche onset (negative values = before menarche onset, positive values = after menarche onset). Credible intervals [CI] were derived from posterior sampling of the fitted generalized additive mixed models (GAMMs). | | | |

| **Supplementary Table 5. Results from GAMMs for internalizing symptoms using the Child Behavior Checklist as outcome variable in the ABCD ARMS** | | | | | |
| --- | --- | --- | --- | --- | --- |
| **ABCD ARM** | **Term** | **edf** | ***F*** | ***p*** | ***p*_Bonferroni_** |
| 1 | s(Gynecological Age) | 5.827 | 39.876 | **<.001** | **<.001** |
|  | s(Menarche Timing) | 1 | 2.202 | 0.138 | 0.827 |
|  | s(General Exposome) | 2.141 | 3.515 | **0.041** | 0.244 |
|  |  |  |  |  | *R*^2^ = 0.012 |
| 2 | s(Gynecological Age) | 4.819 | 35.593 | **<.001** | **<.001** |
|  | s(Menarche Timing) | 1 | 4.201 | **0.040** | 0.243 |
|  | s(General Exposome) | 4.019 | 9.077 | **<.001** | **<.001** |
|  |  |  |  |  | *R*^2^ = 0.017 |
| Note. The term s() represents a smooth function of the predictor, modeling potential nonlinear fluctuations over time. The effective degrees of freedom (edf) indicate the complexity of the smooth term, with higher values suggesting greater flexibility. *p*_Bonferroni_ = *p*-values after Bonferroni correction for multiple comparisons. Significant results are indicated in bold. Total number of observations in ARM 1, *n* = 14,549. Total number of observations in ARM 1, *n* = 14,289. | | | | | |

| **Supplementary Table 6. Second derivative minima and maxima for internalizing symptoms using the Child Behavior Checklist in ABCD ARMS** | | | |
| --- | --- | --- | --- |
| **ABCD ARM** | **Predictor** | **2^nd^ derivative minimum [CI]** | **2^nd^ derivative maximum [CI]** |
| 1 | Gynecological Age | 2.439 [2.144;4.581] | 0.593 [-0.072;1.184] |
| 2 | Gynecological Age | 3.727 [-2.463;4.659] | 0.072 [-0.673;1.676] |
| Note. 2^nd^ derivatives describe the rate of change in the slope of the trajectory and identify inflection points where the curvature changes. The second derivative maximum indicates the point of fastest acceleration (steepest increase in slope), occurring where symptoms are increasing most rapidly. The second derivative minimum indicates the point of maximum deceleration (greatest slowing of the slope), occurring where the rate of increase or decrease is slowing most. For gynecological age, values represent time in years relative to individual menarche onset (negative values = before menarche onset, positive values = after menarche onset). Credible intervals [CI] were derived from posterior sampling of the fitted generalized additive mixed models (GAMMs). | | | |

| **Supplementary Table 7. Results from Generalized Additive Mixed Models (GAMMs) for internalizing symptoms using the Child Behavior Checklist without familial dependencies** | | | | |
| --- | --- | --- | --- | --- |
| **Outcome** | **Term** | **edf** | ***F*** | ***p*** |
| Internalizing symptoms | s(Gynecological Age) | 5.955 | 63.642 | **<.001** |
|  | s(Menarche Timing) | 1 | 3.283 | 0.07 |
|  | s(General Exposome) | 4.056 | 9.122 | **<.001** |
|  |  |  |  | *R*^2^ = 0.014 |
| Note. The term s() represents a smooth function of the predictor, modeling potential nonlinear fluctuations over time. The effective degrees of freedom (edf) indicate the complexity of the smooth term, with higher values suggesting greater flexibility. Significant results are indicated in bold. Total number of observations, *n* = 25,985. | | | | |

| **Supplementary Table 8. Second derivative minima and maxima for internalizing symptoms using the Child Behavior Checklist without familial dependencies** | | | |
| --- | --- | --- | --- |
| **Outcome** | **Predictor** | **2^nd^ derivative minimum [CI]** | **2^nd^ derivative maximum [CI]** |
| Internalizing symptoms | Gynecological Age | 2.536 [-1.896;4.604] | 0.320 [-0.492;0.948] |
| Note. 2^nd^ derivatives describe the rate of change in the slope of the trajectory and identify inflection points where the curvature changes. The second derivative maximum indicates the point of fastest acceleration (steepest increase in slope), occurring where symptoms are increasing most rapidly. The second derivative minimum indicates the point of maximum deceleration (greatest slowing of the slope), occurring where the rate of increase or decrease is slowing most. For chronological age, values represent age in years; for gynecological age, values represent time in years relative to individual menarche onset (negative values = before menarche onset, positive values = after menarche onset). Credible intervals [CI] were derived from posterior sampling of the fitted generalized additive mixed models (GAMMs). | | | |

| **Supplementary Table 9. Results from GAMMs for internalizing symptoms using the Child Behavior Checklist as outcome variable in puberty markers based on parent-report** | | | | | |
| --- | --- | --- | --- | --- | --- |
| **Puberty Marker** | **Term** | **edf** | ***F*** | ***p*** | ***p*_Bonferroni_** |
| Menarche Onset | s(Gynecological Age) | 5.581 | 73.982 | **<.001** | **<.001** |
|  | s(Menarche Timing) | 1 | 1.05 | 0.306 | 1 |
|  | s(General Exposome) | 4.862 | 8.551 | **<.001** | **<.001** |
|  |  |  |  |  | *R*^2^ = 0.013 |
| Growth Spurt Onset | s(Puberty Marker Onset) | 7.497 | 60.603 | **<.001** | **<.001** |
|  | s(Menarche Timing) | 1 | 10.561 | **0.001** | **0.017** |
|  | s(General Exposome) | 5.031 | 7.67 | **<.001** | **<.001** |
|  |  |  |  |  | *R*^2^ = 0.014 |
| Breast Budding Onset | s(Puberty Marker Onset) | 7.679 | 56.309 | **<.001** | **<.001** |
|  | s(Menarche Timing) | 1 | 7.116 | **0.008** | 0.115 |
|  | s(General Exposome) | 4.882 | 8.822 | **<.001** | **<.001** |
|  |  |  |  |  | *R*^2^ = 0.015 |
| Body Hair Onset | s(Puberty Marker Onset) | 4.841 | 75.838 | **<.001** | **<.001** |
|  | s(Menarche Timing) | 1 | 4.695 | **0.03** | 0.454 |
|  | s(General Exposome) | 5.087 | 7.843 | **<.001** | **<.001** |
|  |  |  |  |  | *R*^2^ = 0.014 |
| Skin Change Onset | s(Puberty Marker Onset) | 6.762 | 66.062 | **<.001** | **<.001** |
|  | s(Menarche Timing) | 1 | 7.465 | **0.006** | 0.094 |
|  | s(General Exposome) | 5.25 | 7.576 | **<.001** | **<.001** |
|  |  |  |  |  | *R*^2^ = 0.017 |
| Note. The term s() represents a smooth function of the predictor, modeling potential nonlinear fluctuations over time. The effective degrees of freedom (edf) indicate the complexity of the smooth term, with higher values suggesting greater flexibility. *p*_Bonferroni_ = *p*-values after Bonferroni correction for multiple comparisons. Significant results are indicated in bold. Total number of observations range from *n* = 28,588–29,156. | | | | | |

| **Supplementary Table 10. Second derivative minima and maxima for internalizing symptoms using the Child Behavior Checklist in puberty markers based on parent-report** | | | |
| --- | --- | --- | --- |
| **Puberty Marker** | **Predictor** | **2^nd^ derivative minimum [CI]** | **2^nd^ derivative maximum [CI]** |
| Menarche Onset | Gynecological Age | 3.083 [2.417;4.563] | 0.456 [-0.617;1.418] |
| Growth Spurt Onset | Puberty Marker Onset | 5.105 [4.897;5.372] | 1.959 [1.900;4.066] |
| Breast Budding Onset | Puberty Marker Onset | 5.238 [5.138;5.272] | 1.917 [1.816;4.031] |
| Body Hair Onset | Puberty Marker Onset | 5.071 [3.966;5.218] | 1.831 [-2.071;3.856] |
| Skin Change Onset | Puberty Marker Onset | 5.139 [4.953;5.213] | 1.911 [-2.134;3.951] |
| Note. 2^nd^ derivatives describe the rate of change in the slope of the trajectory and identify inflection points where the curvature changes. The second derivative maximum indicates the point of fastest acceleration (steepest increase in slope), occurring where symptoms are increasing most rapidly. The second derivative minimum indicates the point of maximum deceleration (greatest slowing of the slope), occurring where the rate of increase or decrease is slowing most. For gynecological age, values represent time in years relative to individual menarche onset (negative values = before menarche onset, positive values = after menarche onset). For puberty markers, values represent time in years relative to individual onset (negative values = before onset, positive values = after onset). Credible intervals [CI] were derived from posterior sampling of the fitted generalized additive mixed models (GAMMs). | | | |

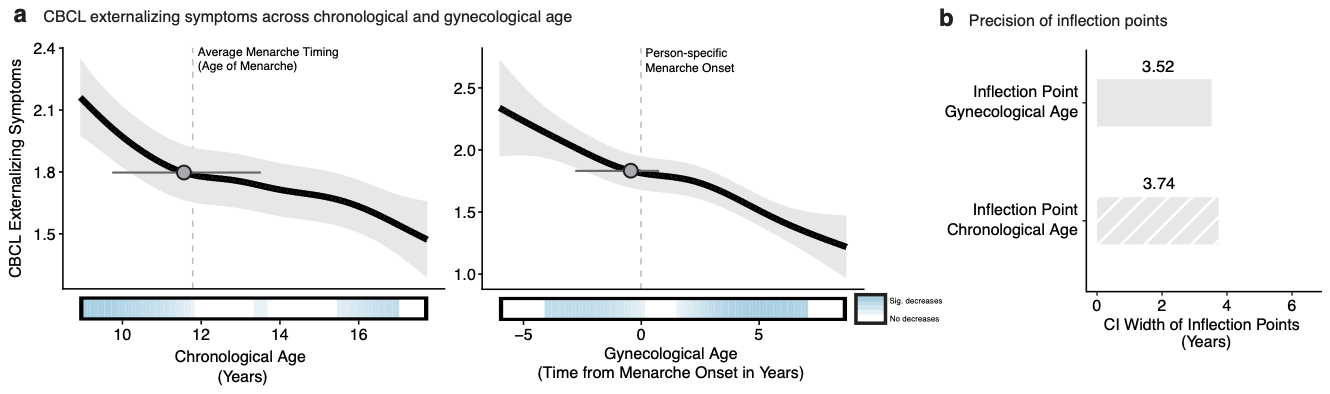

**Supplementary Figure 1.** **Externalizing symptoms across chronological and gynecological age. a**, Child Behavior Checklist (CBCL) externalizing symptom trajectories across chronological age (left) and gynecological age (right). Black lines depict predicted values from generalized additive mixed models (GAMMs) relating chronological and gynecological to externalizing symptoms. Light gray shaded regions represent 95% credible intervals (CI). Inflection points (derived from second derivative maxima, indicating the point of fastest acceleration) are shown as gray points with black outlines, with 95% CI derived from posterior sampling depicted as gray horizontal lines. Colored horizontal bars along the x-axis denote periods of significant change based on first derivative estimates: blue indicates significant decreases, white indicates no significant change. Dashed vertical lines mark average menarche age (left panels) and person-specific menarche onset (right panels, time = 0). **b**, 95% credible interval (CI) widths for the inflection points in externalizing symptoms derived from gynecological and chronological age models, expressed in years. Gary solid and striped bars represent CI widths derived from gynecological and chronological age models, respectively. Black numerical labels on top of the bars indicate the CI width in years. n = 5,016 female participants.

| **Supplementary Table 11. Results from Generalized Additive Mixed Models (GAMMs) for brain measures.** | | | | | |
| --- | --- | --- | --- | --- | --- |
| **Outcome** | **Term** | **edf** | ***F*** | ***p*** | ***p*_Bonferroni_** |
| Whole brain volume | s(Chronological Age) | 7.167 | 1521 | **<.001** | **<.001** |
|  | s(General Exposome) | 4.797 | 147.1 | **<.001** | **<.001** |
|  |  |  |  |  | *R*^2^ = 0.131 |
| Subcortical volume | s(Chronological Age) | 7.206 | 426.1 | **<.001** | **<.001** |
|  | s(General Exposome) | 4.742 | 91.29 | **<.001** | **<.001** |
|  |  |  |  |  | *R*^2^ = 0.089 |
| Gray matter volume | s(Chronological Age) | 6.929 | 8679 | **<.001** | **<.001** |
|  | s(General Exposome) | 4.636 | 178.3 | **<.001** | **<.001** |
|  |  |  |  |  | *R*^2^ = 0.253 |
| Cortical thickness | s(Chronological Age) | 5.588 | 4961 | **<.001** | **<.001** |
|  | s(General Exposome) | 3.905 | 65.56 | **<.001** | **<.001** |
|  |  |  |  |  | *R*^2^ = 0.297 |
| Surface area | s(Chronological Age) | 6.763 | 1684 | **<.001** | **<.001** |
|  | s(General Exposome) | 4.495 | 116.2 | **<.001** | **<.001** |
|  |  |  |  |  | *R*^2^ = 0.103 |
| White matter volume | s(Chronological Age) | 6.831 | 5051 | **<.001** | **<.001** |
|  | s(General Exposome) | 4.407 | 76.23 | **<.001** | **<.001** |
|  |  |  |  |  | *R*^2^ = 0.114 |
| Lateral ventricles volume | s(Chronological Age) | 5.35 | 1079 | **<.001** | **<.001** |
|  | s(General Exposome) | 1 | 5.473 | **0.019** | 1 |
|  |  |  |  |  | *R*^2^ = 0.017 |
| CSF | s(Chronological Age) | 5.044 | 162.9 | **<.001** | **<.001** |
|  | s(General Exposome) | 1 | 0.617 | 0.432 | 1 |
|  |  |  |  |  | *R*^2^ = 0.006 |
| Whole brain volume | s(Menarche Timing) | 1 | 3.946 | **0.047** | 1 |
|  | s(General Exposome) | 4.74 | 128.9 | <.001 | <.001 |
|  |  |  |  |  | *R*^2^ = 0.116 |
| Subcortical volume | s(Menarche Timing) | 2.739 | 1.577 | 0.119 | 1 |
|  | s(General Exposome) | 4.704 | 85.41 | **<.001** | **<.001** |
|  |  |  |  |  | *R*^2^ = 0.084 |
| Gray matter volume | s(Menarche Timing) | 1 | 8.749 | **0.003** | 0.174 |
|  | s(General Exposome) | 4.526 | 139.2 | **<.001** | **<.001** |
|  |  |  |  |  | *R*^2^ = 0.107 |
| Cortical thickness | s(Menarche Timing) | 1.708 | 7.713 | **0.001** | **0.039** |
|  | s(General Exposome) | 3.843 | 37.46 | **<.001** | **<.001** |
|  |  |  |  |  | *R*^2^ = 0.028 |
| Surface area | s(Menarche Timing) | 1 | 3.436 | 0.064 | 1 |
|  | s(General Exposome) | 4.427 | 100.9 | **<.001** | **<.001** |
|  |  |  |  |  | *R*^2^ = 0.086 |
| White matter volume | s(Menarche Timing) | 1 | 2.029 | 0.154 | 1 |
|  | s(General Exposome) | 4.356 | 72.45 | **<.001** | **<.001** |
|  |  |  |  |  | *R*^2^ = 0.063 |
| Lateral ventricles volume | s(Menarche Timing) | 1.432 | 1.34 | 0.178 | 1 |
|  | s(General Exposome) | 1.002 | 8.585 | **0.003** | 0.189 |
|  |  |  |  |  | *R*^2^ = 0.002 |
| CSF | s(Menarche Timing) | 1 | 16.44 | **<.001** | **0.003** |
|  | s(General Exposome) | 1 | 0.164 | 0.686 | 1 |
|  |  |  |  |  | *R*^2^ = 0.003 |
| Whole brain volume | s(Gynecological Age) | 8.685 | 1695 | **<.001** | **<.001** |
|  | s(Menarche Timing) | 1.65 | 0.394 | 0.634 | 1 |
|  | s(General Exposome) | 4.744 | 132.7 | **<.001** | **<.001** |
|  |  |  |  |  | *R*^2^ = 0.133 |
| Subcortical volume | s(Gynecological Age) | 8.531 | 492.1 | **<.001** | **<.001** |
|  | s(Menarche Timing) | 2.808 | 3.972 | **0.006** | 0.345 |
|  | s(General Exposome) | 4.742 | 86.29 | **<.001** | **<.001** |
|  |  |  |  |  | *R*^2^ = 0.091 |
| Gray matter volume | s(Gynecological Age) | 8.575 | 8124 | **<.001** | **<.001** |
|  | s(Menarche Timing) | 1.257 | 56.98 | **<.001** | **<.001** |
|  | s(General Exposome) | 4.518 | 155.7 | **<.001** | **<.001** |
|  |  |  |  |  | *R*^2^ = 0.259 |
| Cortical thickness | s(Gynecological Age) | 7.428 | 3826 | **<.001** | **<.001** |
|  | s(Menarche Timing) | 1 | 203.5 | **<.001** | **<.001** |
|  | s(General Exposome) | 3.778 | 49.72 | **<.001** | **<.001** |
|  |  |  |  |  | *R*^2^ = 0.305 |
| Surface area | s(Gynecological Age) | 8.414 | 1592 | **<.001** | **<.001** |
|  | s(Menarche Timing) | 1.254 | 0.667 | 0.356 | 1 |
|  | s(General Exposome) | 4.425 | 104.2 | **<.001** | **<.001** |
|  |  |  |  |  | *R*^2^ = 0.105 |
| White matter volume | s(Gynecological Age) | 8.372 | 4621 | **<.001** | **<.001** |
|  | s(Menarche Timing) | 1.048 | 79.74 | **<.001** | **<.001** |
|  | s(General Exposome) | 4.403 | 71.42 | **<.001** | **<.001** |
|  |  |  |  |  | *R*^2^ = 0.115 |
| Lateral ventricles volume | s(Gynecological Age) | 7.356 | 818.8 | **<.001** | **<.001** |
|  | s(Menarche Timing) | 1 | 2.264 | 0.132 | 1 |
|  | s(General Exposome) | 1 | 8.05 | **0.005** | 0.255 |
|  |  |  |  |  | *R*^2^ = 0.018 |
| CSF | s(Gynecological Age) | 6.307 | 130.3 | **<.001** | **<.001** |
|  | s(Menarche Timing) | 1 | 4.902 | **0.027** | 1 |
|  | s(General Exposome) | 1 | 0.146 | **0.703** | 1 |
|  |  |  |  |  | *R*^2^ = 0.009 |
| Note. The term s() represents a smooth function of the predictor, modeling potential nonlinear fluctuations over time. The effective degrees of freedom (edf) indicate the complexity of the smooth term, with higher values suggesting greater flexibility. *p*_Bonferroni_ = *p*-values after Bonferroni correction for multiple comparisons. Significant results are indicated in bold. CSF = cerebrospinal fluid.Total number of observations, *n* = 12,971. | | | | | |

| **Supplementary Table 12. Second derivative minima and maxima for brain measures.** | | | |
| --- | --- | --- | --- |
| **Outcome** | **Predictor** | **2^nd^ derivative minimum [CI]** | **2^nd^ derivative maximum [CI]** |
| Whole brain volume | Chronological Age | 12.009 [11.352;12.848] | 14.307 [13.869;16.423] |
| Subcortical volume | Chronological Age | 12.410 [11.753;13.139] | 14.307 [10.878;16.423] |
| Gray matter volume | Chronological Age | 11.461 [10.914;12.556] | 14.416 [9.783;16.423] |
| Cortical thickness | Chronological Age | 11.096 [10.695;16.423] | 14.124 [13.176;16.423] |
| Surface area | Chronological Age | 11.388 [10.841;12.702] | 16.350 [9.820;16.423] |
| White matter volume | Chronological Age | 12.191 [11.644;12.738] | 16.423 [10.476;16.423] |
| Lateral ventricles volume | Chronological Age | 14.234 [12.227;15.401] | 11.024 [9.893;16.423] |
| CSF | Chronological Age | 13.723 [9.783;15.438] | 11.352 [10.805;16.423] |
| Whole brain volume | Gynecological Age | 0.322 [0.202;0.362] | 2.242 [-1.878;4.483] |
| Subcortical volume | Gynecological Age | 0.362 [0.282;0.442] | -1.438 [-1.878;4.603] |
| Gray matter volume | Gynecological Age | 0.242 [0.082;0.322] | 2.202 [2.002;2.442] |
| Cortical thickness | Gynecological Age | 0.122 [-0.678;0.442] | 2.362 [1.922;3.002] |
| Surface area | Gynecological Age | 0.202 [0.082;0.362] | 4.563 [-2.119;4.803] |
| White matter volume | Gynecological Age | 0.402 [0.202;0.522] | -1.558 [-2.038;4.643] |
| Lateral ventricles volume | Gynecological Age | 1.602 [1.002;2.162] | -0.798 [-2.078;-0.238] |
| CSF | Gynecological Age | 1.642 [0.842;3.042] | -0.598 [-2.038;0.242] |
| Note. 2^nd^ derivatives describe the rate of change in the slope of the trajectory and identify inflection points where the curvature changes. The second derivative maximum indicates the point of fastest acceleration (steepest increase in slope), occurring where measures are increasing most rapidly. The second derivative minimum indicates the point of maximum deceleration (greatest slowing of the slope), occurring where the rate of increase or decrease is slowing most. For chronological age, values represent age in years; for gynecological age, values represent time in years relative to individual menarche onset (negative values = before menarche onset, positive values = after menarche onset). Credible intervals [CI] were derived from posterior sampling of the fitted generalized additive mixed models (GAMMs). CSF = cerebrospinal fluid. | | | |

| **Supplementary Table 13. Results from Generalized Additive Mixed Models (GAMMs) for brain measures in ABCD ARMS** | | | | | | | | |
| --- | --- | --- | --- | --- | --- | --- | --- | --- |
| **ABCD ARMS** | **Outcome** | **Term** | **edf** | ***F*** | ***p*** | | | ***p*_Bonferroni_** |
| 1 | Whole brain volume | s(Gynecological Age) | 8.379 | 830.305 | | **<.001** | **<.001** | |
|  |  | s(Menarche Timing) | 1 | 0.056 | | 0.814 | 1 | |
|  |  | s(General Exposome) | 3.965 | 77.555 | | **<.001** | **<.001** | |
|  |  |  |  |  | |  | *R*^2^ = 0.135 | |
|  | Subcortical volume | s(Gynecological Age) | 8.179 | 264.792 | | **<.001** | **<.001** | |
|  |  | s(Menarche Timing) | 1.916 | 2.459 | | 0.137 | 1 | |
|  |  | s(General Exposome) | 3.583 | 59.722 | | **<.001** | **<.001** | |
|  |  |  |  |  | |  | *R*^2^ = 0.094 | |
|  | Gray matter volume | s(Gynecological Age) | 8.189 | 4033.718 | | **<.001** | **<.001** | |
|  |  | s(Menarche Timing) | 1 | 36.266 | | **<.001** | **<.001** | |
|  |  | s(General Exposome) | 3.953 | 84.853 | | **<.001** | **<.001** | |
|  |  |  |  |  | |  | *R*^2^ = 0.256 | |
|  | Cortical thickness | s(Gynecological Age) | 6.729 | 2047.92 | | **<.001** | **<.001** | |
|  |  | s(Menarche Timing) | 2.229 | 44.215 | | **<.001** | **<.001** | |
|  |  | s(General Exposome) | 3.161 | 29.495 | | **<.001** | **<.001** | |
|  |  |  |  |  | |  | *R*^2^ = 0.308 | |
|  | Surface area | s(Gynecological Age) | 8.017 | 800.922 | | **<.001** | **<.001** | |
|  |  | s(Menarche Timing) | 1.484 | 0.391 | | 0.479 | 1 | |
|  |  | s(General Exposome) | 3.697 | 58.81 | | **<.001** | **<.001** | |
|  |  |  |  |  | |  | *R*^2^ = 0.104 | |
|  | White matter volume | s(Gynecological Age) | 7.926 | 2338.715 | | **<.001** | **<.001** | |
|  |  | s(Menarche Timing) | 1.047 | 43.821 | | **<.001** | **<.001** | |
|  |  | s(General Exposome) | 3.322 | 46.066 | | **<.001** | **<.001** | |
|  |  |  |  |  | |  | *R*^2^ = 0.12 | |
|  | Lateral ventricles volume | s(Gynecological Age) | 6.575 | 440.686 | | **<.001** | **<.001** | |
|  |  | s(Menarche Timing) | 1 | 0.124 | | 0.725 | 1 | |
|  |  | s(General Exposome) | 1 | 4.851 | | 0.028 | 1 | |
|  |  |  |  |  | |  | *R*^2^ = 0.018 | |
|  | CSF | s(Gynecological Age) | 5.073 | 71.754 | | **<.001** | **<.001** | |
|  |  | s(Menarche Timing) | 1 | 6.506 | | **0.011** | 0.517 | |
|  |  | s(General Exposome) | 1 | 0.16 | | 0.689 | 1 | |
|  |  |  |  |  | |  | *R*^2^ = 0.01 | |
| 2 | Whole brain volume | s(Gynecological Age) | 8.466 | 865.528 | | **<.001** | **<.001** | |
|  |  | s(Menarche Timing) | 1.57 | 0.246 | | 0.652 | 1 | |
|  |  | s(General Exposome) | 3.925 | 77.466 | | **<.001** | **<.001** | |
|  |  |  |  |  | |  | *R*^2^ = 0.129 | |
|  | Subcortical volume | s(Gynecological Age) | 8.185 | 234.281 | | **<.001** | **<.001** | |
|  |  | s(Menarche Timing) | 2.578 | 2.623 | | **0.038** | 1 | |
|  |  | s(General Exposome) | 4.084 | 46.687 | | **<.001** | **<.001** | |
|  |  |  |  |  | |  | *R*^2^ = 0.087 | |
|  | Gray matter volume | s(Gynecological Age) | 8.284 | 4209.878 | | **<.001** | **<.001** | |
|  |  | s(Menarche Timing) | 1.455 | 22.543 | | **<.001** | **<.001** | |
|  |  | s(General Exposome) | 3.831 | 89.427 | | **<.001** | **<.001** | |
|  |  |  |  |  | |  | *R*^2^ = 0.259 | |
|  | Cortical thickness | s(Gynecological Age) | 6.677 | 2079.729 | | **<.001** | **<.001** | |
|  |  | s(Menarche Timing) | 1.63 | 57.837 | | **<.001** | **<.001** | |
|  |  | s(General Exposome) | 2.628 | 29.953 | | **<.001** | **<.001** | |
|  |  |  |  |  | |  | *R*^2^ = 0.306 | |
|  | Surface area | s(Gynecological Age) | 7.914 | 830.235 | | **<.001** | **<.001** | |
|  |  | s(Menarche Timing) | 1 | 0.504 | | 0.478 | 1 | |
|  |  | s(General Exposome) | 3.93 | 59.204 | | **<.001** | **<.001** | |
|  |  |  |  |  | |  | *R*^2^ = 0.104 | |
|  | White matter volume | s(Gynecological Age) | 8.021 | 2405.786 | | **<.001** | **<.001** | |
|  |  | s(Menarche Timing) | 1 | 34.798 | | **<.001** | **<.001** | |
|  |  | s(General Exposome) | 3.68 | 42.559 | | **<.001** | **<.001** | |
|  |  |  |  |  | |  | *R*^2^ = 0.11 | |
|  | Lateral ventricles volume | s(Gynecological Age) | 6.669 | 434.9 | | **<.001** | **<.001** | |
|  |  | s(Menarche Timing) | 1.648 | 2.892 | | 0.119 | 1 | |
|  |  | s(General Exposome) | 1 | 3.569 | | 0.059 | 1 | |
|  |  |  |  |  | |  | *R*^2^ = 0.017 | |
|  | CSF | s(Gynecological Age) | 5.931 | 72.456 | | **<.001** | **<.001** | |
|  |  | s(Menarche Timing) | 1 | 0.481 | | 0.488 | 1 | |
|  |  | s(General Exposome) | 1 | 0.029 | | 0.864 | 1 | |
|  |  |  |  |  | |  | *R*^2^ = 0.009 | |
| Note. The term s() represents a smooth function of the predictor, modeling potential nonlinear fluctuations over time. The effective degrees of freedom (edf) indicate the complexity of the smooth term, with higher values suggesting greater flexibility. *p*_Bonferroni_ = *p*-values after Bonferroni correction for multiple comparisons. Significant results are indicated in bold. CSF = cerebrospinal fluid. Total number of observations in ARM 1, *n* = 6,345. Total number of observations in ARM 1, *n* = 6,240. | | | | | | | | |

| **Supplementary Table 14. Second derivative minima and maxima for brain measures in ABCD ARMS** | | | | |
| --- | --- | --- | --- | --- |
| **ABCD ARMS** | **Outcome** | **Predictor** | **2^nd^ derivative minimum [CI]** | **2^nd^ derivative maximum [CI]** |
| 1 | Whole brain volume | Gynecological Age | 0.219 [0.059;0.339] | -1.659 [-1.979;4.456] |
|  | Subcortical volume | Gynecological Age | 0.299 [0.099;0.459] | -1.539 [-2.019;4.456] |
|  | Gray matter volume | Gynecological Age | 0.179 [-0.060;0.339] | 2.178 [1.898;4.096] |
|  | Cortical thickness | Gynecological Age | 0.099 [-0.820;4.776] | 2.498 [1.898;3.177] |
|  | Surface area | Gynecological Age | 0.139 [-0.100;0.339] | 4.336 [-3.178;4.536] |
|  | White matter volume | Gynecological Age | 0.259 [-0.260;0.579] | -1.819 [-2.139;4.776] |
|  | Lateral ventricles volume | Gynecological Age | 1.578 [0.699;2.737] | -0.780 [-2.139;0.059] |
|  | CSF | Gynecological Age | 1.298 [-3.178;3.737] | -0.740 [-2.019;0.299] |
| 2 | Whole brain volume | Gynecological Age | 0.305 [0.185;0.426] | 2.154 [-1.945;4.324] |
|  | Subcortical volume | Gynecological Age | 0.386 [0.185;0.506] | -1.383 [-3.151;2.797] |
|  | Gray matter volume | Gynecological Age | 0.225 [0.024;0.345] | 2.114 [1.873;2.556] |
|  | Cortical thickness | Gynecological Age | -0.016 [-1.142;0.546] | 2.355 [1.591;3.802] |
|  | Surface area | Gynecological Age | 0.265 [-0.016;0.426] | 2.154 [-3.151;4.445] |
|  | White matter volume | Gynecological Age | 0.426 [0.265;0.747] | -1.343 [-3.151;3.681] |
|  | Lateral ventricles volume | Gynecological Age | 1.752 [0.988;2.717] | -0.619 [-2.106;-0.016] |
|  | CSF | Gynecological Age | 1.993 [1.189;2.877] | -0.257 [-2.387;4.847] |
| Note. 2^nd^ derivatives describe the rate of change in the slope of the trajectory and identify inflection points where the curvature changes. The second derivative maximum indicates the point of fastest acceleration (steepest increase in slope), occurring where measures are increasing most rapidly. The second derivative minimum indicates the point of maximum deceleration (greatest slowing of the slope), occurring where the rate of increase or decrease is slowing most. For gynecological age, values represent time in years relative to individual menarche onset (negative values = before menarche onset, positive values = after menarche onset). Credible intervals [CI] were derived from posterior sampling of the fitted generalized additive mixed models (GAMMs). CSF = cerebrospinal fluid. | | | | |

| **Supplementary Table 15. Results from Generalized Additive Mixed Models (GAMMs) for brain measures without familial dependencies** | | | | | | | |
| --- | --- | --- | --- | --- | --- | --- | --- |
| **Outcome** | **Term** | **edf** | ***F*** | ***p*** | | | ***p*_Bonferroni_** |
| Whole brain volume | s(Gynecological Age) | 8.648 | 1474.924 | | **<.001** | **<.001** | |
|  | s(Menarche Timing) | 1.654 | 0.405 | | 0.619 | 1 | |
|  | s(General Exposome) | 4.495 | 125.546 | | **<.001** | **<.001** | |
|  |  |  |  | |  | *R*^2^ = 0.136 | |
| Subcortical volume | s(Gynecological Age) | 8.469 | 443.207 | | **<.001** | **<.001** | |
|  | s(Menarche Timing) | 2.431 | 4.034 | | 0.01 | 0.236 | |
|  | s(General Exposome) | 4.582 | 76.947 | | **<.001** | **<.001** | |
|  |  |  |  | |  | *R*^2^ = 0.09 | |
| Gray matter volume | s(Gynecological Age) | 8.519 | 7070.592 | | **<.001** | **<.001** | |
|  | s(Menarche Timing) | 1.524 | 38.166 | | **<.001** | **<.001** | |
|  | s(General Exposome) | 4.162 | 155.498 | | **<.001** | **<.001** | |
|  |  |  |  | |  | *R*^2^ = 0.263 | |
| Cortical thickness | s(Gynecological Age) | 7.22 | 3305.554 | | **<.001** | **<.001** | |
|  | s(Menarche Timing) | 1 | 173.792 | | **<.001** | **<.001** | |
|  | s(General Exposome) | 3.604 | 43.974 | | **<.001** | **<.001** | |
|  |  |  |  | |  | *R*^2^ = 0.301 | |
| Surface area | s(Gynecological Age) | 8.368 | 1359.031 | | **<.001** | **<.001** | |
|  | s(Menarche Timing) | 1.633 | 0.508 | | 0.424 | 1 | |
|  | s(General Exposome) | 4.038 | 105.444 | | **<.001** | **<.001** | |
|  |  |  |  | |  | *R*^2^ = 0.109 | |
| White matter volume | s(Gynecological Age) | 8.316 | 3948.707 | | **<.001** | **<.001** | |
|  | s(Menarche Timing) | 1.053 | 67.055 | | **<.001** | **<.001** | |
|  | s(General Exposome) | 4.153 | 66.822 | | **<.001** | **<.001** | |
|  |  |  |  | |  | *R*^2^ = 0.114 | |
| Lateral ventricles volume | s(Gynecological Age) | 7.262 | 731.66 | | **<.001** | **<.001** | |
|  | s(Menarche Timing) | 1.408 | 1.798 | | 0.256 | 1 | |
|  | s(General Exposome) | 1.006 | 6.648 | | **0.01** | 0.234 | |
|  |  |  |  | |  | 0.018 | |
| CSF | s(Gynecological Age) | 6.09 | 119.397 | | **<.001** | **<.001** | |
|  | s(Menarche Timing) | 1 | 4.173 | | 0.041 | 0.986 | |
|  | s(General Exposome) | 1 | 0.585 | | 0.444 | 1 | |
|  |  |  |  | |  | *R*^2^ = 0.01 | |
| Note. The term s() represents a smooth function of the predictor, modeling potential nonlinear fluctuations over time. The effective degrees of freedom (edf) indicate the complexity of the smooth term, with higher values suggesting greater flexibility. *p*_Bonferroni_ = *p*-values after Bonferroni correction for multiple comparisons. Significant results are indicated in bold. CSF = cerebrospinal fluid. Total number of observations, *n* = 11,305. | | | | | | | |

| **Supplementary Table 16. Second derivative minima and maxima for brain measures without familial dependencies** | | | |
| --- | --- | --- | --- |
| **Outcome** | **Predictor** | **2^nd^ derivative minimum [CI]** | **2^nd^ derivative maximum [CI]** |
| Whole brain volume | Gynecological Age | 0.297 [0.178;0.377] | 2.166 [-1.811;4.632] |
| Subcortical volume | Gynecological Age | 0.377 [0.218;0.456] | -1.453 [-1.970;4.791] |
| Gray matter volume | Gynecological Age | 0.218 [0.019;0.337] | 2.127 [1.928;2.405] |
| Cortical thickness | Gynecological Age | 0.058 [-0.737;0.615] | 2.365 [1.888;3.002] |
| Surface area | Gynecological Age | 0.218 [-0.021;0.337] | 4.632 [-2.924;4.791] |
| White matter volume | Gynecological Age | 0.416 [0.218;0.615] | -1.493 [-2.010;4.791] |
| Lateral ventricles volume | Gynecological Age | 1.570 [0.854;2.166] | -0.737 [-2.010;-0.220] |
| CSF | Gynecological Age | 1.649 [0.814;3.439] | -0.578 [-2.169;0.337] |
| Note. 2^nd^ derivatives describe the rate of change in the slope of the trajectory and identify inflection points where the curvature changes. The second derivative maximum indicates the point of fastest acceleration (steepest increase in slope), occurring where measures are increasing most rapidly. The second derivative minimum indicates the point of maximum deceleration (greatest slowing of the slope), occurring where the rate of increase or decrease is slowing most. For gynecological age, values represent time in years relative to individual menarche onset (negative values = before menarche onset, positive values = after menarche onset). Credible intervals [CI] were derived from posterior sampling of the fitted generalized additive mixed models (GAMMs). CSF = cerebrospinal fluid. | | | |

| **Supplementary Table 17. Results from Generalized Additive Mixed Models (GAMMs) for brain measures in puberty markers based on parent-report** | | | | | | | |
| --- | --- | --- | --- | --- | --- | --- | --- |
| **Puberty Markers** | **Outcome** | **Term** | **edf** | ***F*** | | ***p*** | ***p*_Bonferroni_** |
| Menarche Onset | Whole brain volume | s(Gynecological Age) | 8.639 | 1674.972 | **<.001** | | **<.001** |
|  |  | s(Menarche Timing) | 1 | 5.429 | **0.02** | | 1 |
|  |  | s(General Exposome) | 4.795 | 113.503 | **<.001** | | **<.001** |
|  |  |  |  |  |  | | *R*^2^ = 0.134 |
|  | Subcortical volume | s(Gynecological Age) | 8.543 | 489.323 | **<.001** | | **<.001** |
|  |  | s(Menarche Timing) | 3.314 | 8.064 | **<.001** | | **0.003** |
|  |  | s(General Exposome) | 4.772 | 73.992 | **<.001** | | **<.001** |
|  |  |  |  |  |  | | *R*^2^ = 0.091 |
|  | Gray matter volume | s(Gynecological Age) | 8.512 | 8059.951 | **<.001** | | **<.001** |
|  |  | s(Menarche Timing) | 1 | 38.102 | **<.001** | | **<.001** |
|  |  | s(General Exposome) | 4.495 | 136.095 | **<.001** | | **<.001** |
|  |  |  |  |  |  | | *R*^2^ = 0.262 |
|  | Cortical thickness | s(Gynecological Age) | 7.357 | 3786.448 | **<.001** | | **<.001** |
|  |  | s(Menarche Timing) | 1 | 188.418 | **<.001** | | **<.001** |
|  |  | s(General Exposome) | 3.781 | 45.169 | **<.001** | | **<.001** |
|  |  |  |  |  |  | | *R*^2^ = 0.304 |
|  | Surface area | s(Gynecological Age) | 8.317 | 1576.501 | **<.001** | | **<.001** |
|  |  | s(Menarche Timing) | 1 | 1.263 | 0.261 | | 1 |
|  |  | s(General Exposome) | 4.444 | 89.038 | **<.001** | | **<.001** |
|  |  |  |  |  |  | | *R*^2^ = 0.106 |
|  | White matter volume | s(Gynecological Age) | 8.336 | 4538.878 | **<.001** | | **<.001** |
|  |  | s(Menarche Timing) | 1 | 122.447 | **<.001** | | **<.001** |
|  |  | s(General Exposome) | 4.557 | 58.724 | **<.001** | | **<.001** |
|  |  |  |  |  |  | | *R*^2^ = 0.113 |
|  | Lateral ventricles volume | s(Gynecological Age) | 7.406 | 801.155 | **<.001** | | **<.001** |
|  |  | s(Menarche Timing) | 1 | 2.714 | 0.1 | | 1 |
|  |  | s(General Exposome) | 1 | 6.097 | **0.014** | | 1 |
|  |  |  |  |  |  | | *R*^2^ = 0.017 |
|  | CSF | s(Gynecological Age) | 6.647 | 122.682 | **<.001** | | **<.001** |
|  |  | s(Menarche Timing) | 1 | 4.194 | **0.041** | | 1 |
|  |  | s(General Exposome) | 1 | 0.196 | 0.658 | | 1 |
|  |  |  |  |  |  | | *R*^2^ = 0.009 |
| Growth Spurt | Whole brain volume | s(Puberty Marker Onset) | 8.307 | 1072.642 | **<.001** | | **<.001** |
|  |  | s(Menarche Timing) | 1 | 20.402 | **<.001** | | **0.001** |
|  |  | s(General Exposome) | 4.957 | 105.639 | **<.001** | | **<.001** |
|  |  |  |  |  |  | | *R*^2^ = 0.127 |
|  | Subcortical volume | s(Puberty Marker Onset) | 7.953 | 239.422 | **<.001** | | **<.001** |
|  |  | s(Menarche Timing) | 3.251 | 4.432 | **0.006** | | 0.779 |
|  |  | s(General Exposome) | 4.782 | 72.607 | **<.001** | | **<.001** |
|  |  |  |  |  |  | | *R*^2^ = 0.087 |
|  | Gray matter volume | s(Puberty Marker Onset) | 7.851 | 7091.909 | **<.001** | | **<.001** |
|  |  | s(Menarche Timing) | 1.994 | 13.728 | **<.001** | | **<.001** |
|  |  | s(General Exposome) | 4.671 | 121.581 | **<.001** | | **<.001** |
|  |  |  |  |  |  | | *R*^2^ = 0.231 |
|  | Cortical thickness | s(Puberty Marker Onset) | 5.854 | 4561.5 | **<.001** | | **<.001** |
|  |  | s(Menarche Timing) | 2.566 | 6.419 | **0.001** | | 0.073 |
|  |  | s(General Exposome) | 3.953 | 40.378 | **<.001** | | **<.001** |
|  |  |  |  |  |  | | *R*^2^ = 0.275 |
|  | Surface area | s(Puberty Marker Onset) | 8.243 | 1223.632 | **<.001** | | **<.001** |
|  |  | s(Menarche Timing) | 1 | 17.207 | **<.001** | | **0.004** |
|  |  | s(General Exposome) | 4.627 | 82.013 | **<.001** | | **<.001** |
|  |  |  |  |  |  | | *R*^2^ = 0.098 |
|  | White matter volume | s(Puberty Marker Onset) | 8.26 | 3629.091 | **<.001** | | **<.001** |
|  |  | s(Menarche Timing) | 1.017 | 13.973 | **<.001** | | **0.021** |
|  |  | s(General Exposome) | 4.721 | 53.806 | **<.001** | | **<.001** |
|  |  |  |  |  |  | | *R*^2^ = 0.109 |
|  | Lateral ventricles volume | s(Puberty Marker Onset) | 5.331 | 1110.107 | **<.001** | | **<.001** |
|  |  | s(Menarche Timing) | 1.108 | 0.934 | 0.304 | | 1 |
|  |  | s(General Exposome) | 1 | 5.5 | **0.019** | | 1 |
|  |  |  |  |  |  | | *R*^2^ = 0.014 |
|  | CSF | s(Puberty Marker Onset) | 4.983 | 150.7 | **<.001** | | **<.001** |
|  |  | s(Menarche Timing) | 1 | 11.932 | **0.001** | | 0.066 |
|  |  | s(General Exposome) | 1 | 0.726 | 0.394 | | 1 |
|  |  |  |  |  |  | | *R*^2^ = 0.007 |
| Breast Budding | Whole brain volume | s(Puberty Marker Onset) | 8.351 | 1333.064 | **<.001** | | **<.001** |
|  |  | s(Menarche Timing) | 1 | 16.507 | **<.001** | | **0.006** |
|  |  | s(General Exposome) | 4.835 | 111.103 | **<.001** | | **<.001** |
|  |  |  |  |  |  | | *R*^2^ = 0.131 |
|  | Subcortical volume | s(Puberty Marker Onset) | 8.278 | 351.067 | **<.001** | | **<.001** |
|  |  | s(Menarche Timing) | 3.11 | 5.549 | **0.001** | | 0.133 |
|  |  | s(General Exposome) | 4.812 | 74.529 | **<.001** | | **<.001** |
|  |  |  |  |  |  | | *R*^2^ = 0.09 |
|  | Gray matter volume | s(Puberty Marker Onset) | 8.085 | 7513.739 | **<.001** | | **<.001** |
|  |  | s(Menarche Timing) | 3.872 | 3.653 | **0.01** | | 1 |
|  |  | s(General Exposome) | 4.556 | 123.113 | **<.001** | | **<.001** |
|  |  |  |  |  |  | | *R*^2^ = 0.242 |
|  | Cortical thickness | s(Puberty Marker Onset) | 6.512 | 4173.934 | **<.001** | | **<.001** |
|  |  | s(Menarche Timing) | 4.722 | 3.833 | **0.006** | | 0.73 |
|  |  | s(General Exposome) | 3.754 | 37.981 | **<.001** | | **<.001** |
|  |  |  |  |  |  | | *R*^2^ = 0.289 |
|  | Surface area | s(Puberty Marker Onset) | 7.953 | 1416.888 | **<.001** | | **<.001** |
|  |  | s(Menarche Timing) | 1 | 11.256 | **0.001** | | 0.095 |
|  |  | s(General Exposome) | 4.446 | 86.595 | **<.001** | | **<.001** |
|  |  |  |  |  |  | | *R*^2^ = 0.1 |
|  | White matter volume | s(Puberty Marker Onset) | 8.007 | 4199.91 | **<.001** | | **<.001** |
|  |  | s(Menarche Timing) | 1.037 | 33.418 | **<.001** | | **<.001** |
|  |  | s(General Exposome) | 4.597 | 59.958 | **<.001** | | **<.001** |
|  |  |  |  |  |  | | *R*^2^ = 0.107 |
|  | Lateral ventricles volume | s(Gynecological Age) | 6.427 | 882.513 | **<.001** | | **<.001** |
|  |  | s(Menarche Timing) | 1.156 | 0.097 | 0.764 | | 1 |
|  |  | s(General Exposome) | 1 | 6.539 | **0.011** | | 1 |
|  |  |  |  |  |  | | *R*^2^ = 0.016 |
|  | CSF | s(Puberty Marker Onset) | 6.796 | 116.719 | **<.001** | | **<.001** |
|  |  | s(Menarche Timing) | 1 | 9.899 | **0.002** | | 0.199 |
|  |  | s(General Exposome) | 1 | 0.369 | 0.544 | | 1 |
|  |  |  |  |  |  | | *R*^2^ = 0.01 |
| Body Hair | Whole brain volume | s(Puberty Marker Onset) | 8.382 | 1094.04 | **<.001** | | **<.001** |
|  |  | s(Menarche Timing) | 1.017 | 15.011 | **<.001** | | **0.012** |
|  |  | s(General Exposome) | 5.004 | 104.142 | **<.001** | | **<.001** |
|  |  |  |  |  |  | | *R*^2^ = 0.129 |
|  | Subcortical volume | s(Puberty Marker Onset) | 8.411 | 267.313 | **<.001** | | **<.001** |
|  |  | s(Menarche Timing) | 3.154 | 5.574 | **0.001** | | 0.134 |
|  |  | s(General Exposome) | 4.801 | 74.996 | **<.001** | | **<.001** |
|  |  |  |  |  |  | | *R*^2^ = 0.088 |
|  | Gray matter volume | s(Puberty Marker Onset) | 7.988 | 7016.272 | **<.001** | | **<.001** |
|  |  | s(Menarche Timing) | 2.918 | 2.492 | 0.072 | | 1 |
|  |  | s(General Exposome) | 4.711 | 111.912 | **<.001** | | **<.001** |
|  |  |  |  |  |  | | *R*^2^ = 0.236 |
|  | Cortical thickness | s(Puberty Marker Onset) | 6.144 | 4253.421 | **<.001** | | **<.001** |
|  |  | s(Menarche Timing) | 3.821 | 1.759 | 0.125 | | 1 |
|  |  | s(General Exposome) | 3.817 | 31.737 | **<.001** | | **<.001** |
|  |  |  |  |  |  | | *R*^2^ = 0.25 |
|  | Surface area | s(Puberty Marker Onset) | 7.983 | 1260.833 | **<.001** | | **<.001** |
|  |  | s(Menarche Timing) | 1 | 11.243 | **0.001** | | 0.096 |
|  |  | s(General Exposome) | 4.641 | 80.15 | **<.001** | | **<.001** |
|  |  |  |  |  |  | | *R*^2^ = 0.102 |
|  | White matter volume | s(Puberty Marker Onset) | 8.151 | 3690.214 | **<.001** | | **<.001** |
|  |  | s(Menarche Timing) | 1 | 31.883 | **<.001** | | **<.001** |
|  |  | s(General Exposome) | 4.773 | 57.572 | **<.001** | | **<.001** |
|  |  |  |  |  |  | | *R*^2^ = 0.098 |
|  | Lateral ventricles volume | s(Puberty Marker Onset) | 5.988 | 905.617 | **<.001** | | **<.001** |
|  |  | s(Menarche Timing) | 1 | 0.053 | 0.818 | | 1 |
|  |  | s(General Exposome) | 1 | 7.449 | 0.006 | | 0.763 |
|  |  |  |  |  |  | | *R*^2^ = 0.016 |
|  | CSF | s(Puberty Marker Onset) | 5.479 | 134.551 | **<.001** | | **<.001** |
|  |  | s(Menarche Timing) | 1 | 8.781 | **0.003** | | 0.366 |
|  |  | s(General Exposome) | 1 | 0.936 | 0.333 | | 1 |
|  |  |  |  |  |  | | *R*^2^ = 0.008 |
| Skin Change | Whole brain volume | s(Puberty Marker Onset) | 8.568 | 1048.415 | **<.001** | | **<.001** |
|  |  | s(Menarche Timing) | 1 | 14.707 | **<.001** | | **0.015** |
|  |  | s(General Exposome) | 4.848 | 110.108 | **<.001** | | **<.001** |
|  |  |  |  |  |  | | *R*^2^ = 0.126 |
|  | Subcortical volume | s(Puberty Marker Onset) | 8.491 | 237.813 | **<.001** | | **<.001** |
|  |  | s(Menarche Timing) | 3.126 | 4.909 | **0.003** | | 0.352 |
|  |  | s(General Exposome) | 4.691 | 71.534 | **<.001** | | **<.001** |
|  |  |  |  |  |  | | *R*^2^ = 0.086 |
|  | Gray matter volume | s(Puberty Marker Onset) | 8.076 | 6910.248 | **<.001** | | **<.001** |
|  |  | s(Menarche Timing) | 1 | 4.675 | **0.031** | | 1 |
|  |  | s(General Exposome) | 4.714 | 129.068 | **<.001** | | **<.001** |
|  |  |  |  |  |  | | *R*^2^ = 0.222 |
|  | Cortical thickness | s(Puberty Marker Onset) | 6.082 | 4330.018 | **<.001** | | **<.001** |
|  |  | s(Menarche Timing) | 1.664 | 0.647 | 0.617 | | 1 |
|  |  | s(General Exposome) | 4.1 | 46.179 | **<.001** | | **<.001** |
|  |  |  |  |  |  | | *R*^2^ = 0.244 |
|  | Surface area | s(Puberty Marker Onset) | 8.351 | 1211.975 | **<.001** | | **<.001** |
|  |  | s(Menarche Timing) | 1 | 10.349 | **0.001** | | 0.156 |
|  |  | s(General Exposome) | 4.569 | 85.051 | **<.001** | | **<.001** |
|  |  |  |  |  |  | | *R*^2^ = 0.098 |
|  | White matter volume | s(Puberty Marker Onset) | 8.486 | 3507.066 | **<.001** | | **<.001** |
|  |  | s(Menarche Timing) | 1.009 | 30.553 | **<.001** | | **<.001** |
|  |  | s(General Exposome) | 4.413 | 53.535 | **<.001** | | **<.001** |
|  |  |  |  |  |  | | *R*^2^ = 0.103 |
|  | Lateral ventricles volume | s(Puberty Marker Onset) | 5.844 | 930.221 | **<.001** | | **<.001** |
|  |  | s(Menarche Timing) | 1 | 0.089 | 0.766 | | 1 |
|  |  | s(General Exposome) | 1 | 5.443 | **0.02** | | 1 |
|  |  |  |  |  |  | | *R*^2^ = 0.016 |
|  | CSF | s(Puberty Marker Onset) | 5.537 | 137.889 | **<.001** | | **<.001** |
|  |  | s(Menarche Timing) | 1 | 8.787 | **0.003** | | 0.365 |
|  |  | s(General Exposome) | 1 | 0.061 | 0.805 | | 1 |
|  |  |  |  |  |  | | *R*^2^ = 0.009 |
| Note. The term s() represents a smooth function of the predictor, modeling potential nonlinear fluctuations over time. The effective degrees of freedom (edf) indicate the complexity of the smooth term, with higher values suggesting greater flexibility. *p*_Bonferroni_ = *p*-values after Bonferroni correction for multiple comparisons. Significant results are indicated in bold. CSF = cerebrospinal fluid. Total number of observations range from *n* = 12,432–12,685. | | | | | | | |

| **Supplementary Table 18. Second derivative minima and maxima for brain measures in puberty markers**  **Based on parent-report** | | | | |
| --- | --- | --- | --- | --- |
| **Puberty Markers** | **Outcome** | **Predictor** | **2^nd^ derivative minimum [CI]** | **2^nd^ derivative maximum [CI]** |
| Menarche Onset | Whole brain volume | Gynecological Age | 0.265 [0.145;0.345] | 2.263 [-2.891;2.862] |
|  | Subcortical volume | Gynecological Age | 0.345 [0.265;0.425] | -1.493 [-3.211;2.583] |
|  | Gray matter volume | Gynecological Age | 0.105 [-0.094;0.265] | 2.183 [1.983;2.583] |
|  | Cortical thickness | Gynecological Age | -0.094 [-0.694;0.265] | 2.063 [1.624;2.782] |
|  | Surface area | Gynecological Age | 0.225 [-0.094;0.385] | 2.383 [-3.211;4.141] |
|  | White matter volume | Gynecological Age | 0.465 [0.305;0.705] | -1.613 [-3.211;3.142] |
|  | Lateral ventricles volume | Gynecological Age | 1.704 [1.304;2.263] | -1.453 [-2.172;-0.174] |
|  | CSF | Gynecological Age | 1.424 [0.745;1.983] | -0.654 [-1.533;0.025] |
| Growth Spurt | Whole brain volume | Puberty Marker Onset | 1.904 [1.833;1.904] | 5.748 [2.972;5.855] |
|  | Subcortical volume | Puberty Marker Onset | 1.904 [1.833;1.940] | 5.784 [4.965;5.891] |
|  | Gray matter volume | Puberty Marker Onset | 1.904 [1.833;1.975] | 5.819 [3.043;5.891] |
|  | Cortical thickness | Puberty Marker Onset | 1.940 [1.050;5.926] | 3.897 [3.150;5.891] |
|  | Surface area | Puberty Marker Onset | 1.904 [1.833;1.975] | 5.855 [5.357;5.891] |
|  | White matter volume | Puberty Marker Onset | 1.869 [1.833;4.040] | 5.285 [2.865;5.855] |
|  | Lateral ventricles volume | Puberty Marker Onset | 3.862 [3.008;5.891] | -0.979 [-1.014;5.926] |
|  | CSF | Puberty Marker Onset | 3.826 [3.008;5.891] | -1.014 [-1.014;5.891] |
| Breast Budding | Whole brain volume | Puberty Marker Onset | 1.203 [1.042;1.727] | -0.934 [-0.975;5.760] |
|  | Subcortical volume | Puberty Marker Onset | 1.324 [1.001;1.767] | -0.934 [-0.975;5.760] |
|  | Gray matter volume | Puberty Marker Onset | 1.163 [1.001;1.767] | 3.139 [-1.015;5.437] |
|  | Cortical thickness | Puberty Marker Onset | 1.122 [-2.023;1.929] | 3.340 [3.018;4.106] |
|  | Surface area | Puberty Marker Onset | 1.122 [1.001;1.727] | -0.934 [-1.015;5.760] |
|  | White matter volume | Puberty Marker Onset | 1.122 [1.001;1.808] | -0.934 [-1.015;-0.894] |
|  | Lateral ventricles volume | Puberty Marker Onset | 3.784 [2.009;4.066] | 0.840 [-2.023;5.760] |
|  | CSF | Puberty Marker Onset | 3.139 [2.977;3.784] | 4.953 [-2.023;5.195] |
| Body Hair | Whole brain volume | Puberty Marker Onset | 1.834 [1.748;1.877] | 3.118 [3.033;3.161] |
|  | Subcortical volume | Puberty Marker Onset | 1.834 [1.791;1.877] | 3.075 [3.033;5.388] |
|  | Gray matter volume | Puberty Marker Onset | 1.834 [1.748;1.919] | 3.161 [3.075;3.332] |
|  | Cortical thickness | Puberty Marker Onset | -0.949 [-2.020;5.216] | 3.461 [3.033;4.274] |
|  | Surface area | Puberty Marker Onset | 1.791 [1.577;1.877] | 3.075 [3.033;5.302] |
|  | White matter volume | Puberty Marker Onset | 1.791 [1.534;1.877] | 3.075 [-0.992;5.473] |
|  | Lateral ventricles volume | Puberty Marker Onset | 3.118 [1.962;4.146] | -0.992 [-2.063;5.345] |
|  | CSF | Puberty Marker Onset | 3.204 [1.962;4.017] | 5.173 [-2.105;5.345] |
| Skin Change | Whole brain volume | Puberty Marker Onset | 1.793 [1.619;1.836] | 3.054 [3.054;5.404] |
|  | Subcortical volume | Puberty Marker Onset | 1.749 [1.532;4.055] | 5.404 [3.054;5.447] |
|  | Gray matter volume | Puberty Marker Onset | 1.793 [1.488;1.880] | 3.098 [3.054;5.317] |
|  | Cortical thickness | Puberty Marker Onset | -0.861 [-1.992;1.967] | 3.402 [2.228;5.230] |
|  | Surface area | Puberty Marker Onset | 1.749 [1.532;4.055] | 5.404 [3.054;5.404] |
|  | White matter volume | Puberty Marker Onset | 1.749 [1.619;4.098] | 5.447 [3.011;5.447] |
|  | Lateral ventricles volume | Puberty Marker Onset | 3.098 [1.923;4.011] | -0.948 [-1.992;5.360] |
|  | CSF | Puberty Marker Onset | 3.098 [1.923;4.011] | 5.143 [-1.992;5.447] |
| Note. 2^nd^ derivatives describe the rate of change in the slope of the trajectory and identify inflection points where the curvature changes. The second derivative maximum indicates the point of fastest acceleration (steepest increase in slope), occurring where measures are increasing most rapidly. The second derivative minimum indicates the point of maximum deceleration (greatest slowing of the slope), occurring where the rate of increase or decrease is slowing most. For gynecological age, values represent time in years relative to individual menarche onset (negative values = before menarche onset, positive values = after menarche onset). For puberty markers onset, values represent time in years relative to individual onset (negative values = before onset, positive values = after onset). Credible intervals [CI] were derived from posterior sampling of the fitted generalized additive mixed models (GAMMs). | | | | |

| **Supplementary Table 19. Results from moderation analyses with internalizing symptoms using the Child Behavior Checklist as outcome variable.** | | | | |
| --- | --- | --- | --- | --- |
| **Term** | **edf** | ***F*** | ***p*** | ***p*_Bonferroni_** |
| s(Gynecological Age) | 4.866 | 25.583 | **<.001** | **<.001** |
| s(Whole brain volume) | 1 | 0.159 | 0.690 | 1 |
| ti(Gynecological Age,Whole brain volume) | 1 | 3.621 | 0.057 | 1 |
| s(Menarche Timing) | 1 | 5.806 | **0.016** | 0.640 |
| s(General Exposome) | 4.357 | 4.597 | **0.001** | **0.020** |
|  |  |  |  | *R*^2^ = 0.011 |
| s(Gynecological Age) | 4.774 | 26.989 | **<.001** | **<.001** |
| s(Subcortical volume) | 1 | 1.343 | 0.246 | 1 |
| ti(Gynecological Age,Subcortical volume) | 1 | 1.247 | 0.264 | 1 |
| s(Menarche Timing) | 1 | 5.587 | **0.018** | 0.724 |
| s(General Exposome) | 4.428 | 5.901 | **<.001** | **0.006** |
|  |  |  |  | *R*^2^ = 0.011 |
| s(Gynecological Age) | 5.125 | 17.838 | **<.001** | **<.001** |
| s(Gray matter volume) | 1 | 1.129 | 0.288 | 1 |
| ti(Gynecological Age,Gray matter volume) | 1.636 | 6.759 | **0.024** | 0.974 |
| s(Menarche Timing) | 1 | 5.169 | **0.023** | 0.921 |
| s(General Exposome) | 4.322 | 4.362 | **0.001** | **0.033** |
|  |  |  |  | *R*^2^ = 0.011 |
| s(Gynecological Age) | 5.035 | 18.651 | **<.001** | **<.001** |
| s(Cortical thickness) | 1 | 3.902 | **0.048** | 1 |
| ti(Gynecological Age,Cortical thickness) | 1 | 5.943 | **0.015** | 0.592 |
| s(Menarche Timing) | 1 | 4.36 | **0.037** | 1 |
| s(General Exposome) | 4.343 | 5.193 | **0.001** | **0.023** |
|  |  |  |  | *R*^2^ = 0.011 |
| s(Gynecological Age) | 4.953 | 24.426 | **<.001** | **<.001** |
| s(Surface area) | 1 | 0.074 | 0.786 | 1 |
| ti(Gynecological Age,Surface area) | 1.715 | 3.058 | 0.147 | 1 |
| s(Menarche Timing) | 1 | 5.835 | **0.016** | 0.629 |
| s(General Exposome) | 4.373 | 4.703 | **<.001** | **0.016** |
|  |  |  |  | *R*^2^ = 0.011 |
| s(Gynecological Age) | 4.813 | 23.478 | **<.001** | **<.001** |
| s(White matter volume) | 1 | 0.145 | 0.703 | 1 |
| ti(Gynecological Age,White matter volume) | 1 | 0.892 | 0.345 | 1 |
| s(Menarche Timing) | 1 | 5.513 | **0.019** | 0.756 |
| s(General Exposome) | 4.404 | 4.956 | **<.001** | **0.010** |
|  |  |  |  | *R*^2^ = 0.011 |
| s(Gynecological Age) | 4.849 | 23.876 | **<.001** | **<.001** |
| s(Lateral ventricles volume) | 1.788 | 0.886 | 0.347 | 1 |
| ti(Gynecological Age,Lateral ventricles volume) | 1 | 0.136 | 0.712 | 1 |
| s(Menarche Timing) | 1 | 6.143 | **0.013** | 0.528 |
| s(General Exposome) | 4.4 | 5.656 | **<.001** | **0.009** |
|  |  |  |  | *R*^2^ = 0.011 |
| s(Gynecological Age) | 4.837 | 25.187 | **<.001** | **<.001** |
| s(CSF) | 2.551 | 2.285 | 0.144 | 1 |
| ti(Gynecological Age,CSF) | 1 | 0.243 | 0.622 | 1 |
| s(Menarche Timing) | 1 | 5.777 | **0.016** | 0.650 |
| s(General Exposome) | 4.4 | 5.625 | **<.001** | **0.010** |
|  |  |  |  | *R*^2^ = 0.011 |
| Note. The term s() represents a smooth function of the predictor, modeling potential nonlinear fluctuations over time. The effective degrees of freedom (edf) indicate the complexity of the smooth term, with higher values suggesting greater flexibility. The term ti() represents the interaction between two predictors. *p*_Bonferroni_ = *p*-values after Bonferroni correction for multiple comparisons. Significant results are indicated in bold. Total number of observations, *n* = 12,910. | | | | |

| **Supplementary Table 20. Results from Generalized Additive Mixed Models (GAMMs) with individual post-menarche brain pace and internalizing symptoms using the Child Behavior Checklist as outcome.** | | | | |
| --- | --- | --- | --- | --- |
| **Term** | **edf** | ***F*** | ***p*** | ***p*_Bonferroni_** |
| s(Gynecological Age) | 6.03 | 71.041 | **<.001** | **<.001** |
| s(Post Menarche Whole brain volume pace) | 1 | 1.575 | 0.209 | 1 |
| ti(Gynecological Age,Post Menarche Whole brain volume pace) | 1 | 5.24 | **0.022** | 0.883 |
| s(Menarche Timing) | 1 | 4.488 | **0.034** | 1 |
| s(General Exposome) | 4.812 | 8.063 | **<.001** | **<.001** |
|  |  |  |  | *R*^2^ = 0.014 |
| s(Gynecological Age) | 6.032 | 70.935 | **<.001** | **<.001** |
| s(Post Menarche Subcortical volume pace) | 1 | 1.819 | 0.177 | 1 |
| ti(Gynecological Age,Post Menarche Gray matter volume pace) | 1 | 3.863 | **0.049** | 1 |
| s(Menarche Timing) | 1 | 4.337 | **0.037** | 1 |
| s(General Exposome) | 4.808 | 7.88 | **<.001** | **<.001** |
|  |  |  |  | *R*^2^ = 0.014 |
| s(Gynecological Age) | 6.02 | 71.037 | **<.001** | **<.001** |
| s(Post Menarche Gray matter volume pace) | 1 | 0.002 | 0.964 | 1 |
| ti(Gynecological Age,Post Menarche Cortical thickness pace) | 1 | 10.76 | **0.001** | **0.042** |
| s(Menarche Timing) | 1 | 6.272 | **0.012** | 0.491 |
| s(General Exposome) | 4.817 | 9.252 | **<.001** | **<.001** |
|  |  |  |  | *R*^2^ = 0.014 |
| s(Gynecological Age) | 6.05 | 70.308 | **<.001** | **<.001** |
| s(Post Menarche Cortical thickness pace) | 1 | 0.225 | 0.635 | 1 |
| ti(Gynecological Age,Post Menarche Cortical thickness pace) | 1 | 0.195 | 0.659 | 1 |
| s(Menarche Timing) | 1 | 6.356 | **0.012** | 0.468 |
| s(General Exposome) | 4.837 | 9.313 | **<.001** | **<.001** |
|  |  |  |  | *R*^2^ = 0.014 |
| s(Gynecological Age) | 6.024 | 71.086 | **<.001** | **<.001** |
| s(Post Menarche Surface area pace) | 1 | 1.332 | 0.248 | 1 |
| ti(Gynecological Age,Post Menarche Surface area pace) | 1 | 13.151 | **<.001** | **0.012** |
| s(Menarche Timing) | 1 | 5.261 | **0.022** | 0.873 |
| s(General Exposome) | 4.827 | 9.23 | **<.001** | **<.001** |
|  |  |  |  | *R*^2^ = 0.014 |
| s(Gynecological Age) | 6.044 | 70.536 | **<.001** | **<.001** |
| s(Post Menarche White matter volume pace) | 1 | 1.499 | 0.221 | 1 |
| ti(Gynecological Age,Post Menarche White matter volume pace) | 1 | 0.816 | 0.366 | 1 |
| s(Menarche Timing) | 1 | 4.717 | **0.030** | 1 |
| s(General Exposome) | 4.801 | 7.774 | **<.001** | **<.001** |
|  |  |  |  | *R*^2^ = 0.014 |
| s(Gynecological Age) | 6.042 | 70.542 | **<.001** | **<.001** |
| s(Post Menarche Lateral ventricle pace) | 2.617 | 5.104 | **0.008** | 0.315 |
| ti(Gynecological Age,Post Menarche Lateral ventricle pace) | 1 | 0.092 | 0.761 | 1 |
| s(Menarche Timing) | 1 | 5.734 | **0.017** | 0.666 |
| s(General Exposome) | 4.858 | 9.313 | **<.001** | **<.001** |
|  |  |  |  | *R*^2^ = 0.016 |
| s(Gynecological Age) | 6.043 | 70.444 | **<.001** | **<.001** |
| s(Post Menarche CSF pace) | 1 | 8.272 | **0.004** | 0.161 |
| ti(Gynecological Age,Post Menarche CSF pace) | 1 | 0.078 | 0.78 | 1 |
| s(Menarche Timing) | 1 | 5.973 | **0.015** | 0.581 |
| s(General Exposome) | 4.81 | 9.43 | **<.001** | **<.001** |
|  |  |  |  | *R*^2^ = 0.016 |
| Note. The term s() represents a smooth function of the predictor, modeling potential nonlinear fluctuations over time. The effective degrees of freedom (edf) indicate the complexity of the smooth term, with higher values suggesting greater flexibility. The term ti() represents the interaction between two predictors. *p*_Bonferroni_ = *p*-values after Bonferroni correction for multiple comparisons. Significant results are indicated in bold. CSF = cerebrospinal fluid. Total number of observations, *n* = 29,650. | | | | |

| **Supplementary Table 21. Results from Generalized Additive Mixed Models (GAMMs) for internalizing symptoms with interaction terms using the Child Behavior Checklist.** | | | | |
| --- | --- | --- | --- | --- |
| **Outcome** | **Term** | **edf** | ***F*** | ***p*** |
| Internalizing symptoms | s(Gynecological Age) | 6.274 | 48.8 | **<.001** |
|  | s(Menarche Timing) | 1 | 10.44 | **0.001** |
|  | s(General Exposome) | 4.641 | 8.688 | **<.001** |
|  | ti(Gynecological Age,General Exposome) | 5.033 | 33.33 | **<.001** |
|  | ti(Menarche Timing,General Exposome) | 1 | 1.896 | 0.168 |
|  | ti(Gynecological Age, Menarche Timing) | 5.013 | 2.785 | **0.016** |
|  |  |  |  | *R*^2^ = 0.016 |
| Note. The term s() represents a smooth function of the predictor, modeling potential nonlinear fluctuations over time. The effective degrees of freedom (edf) indicate the complexity of the smooth term, with higher values suggesting greater flexibility. The term ti() represents the interaction between two predictors. Significant results are indicated in bold. Total number of observations, *n* = 29,837. | | | | |

| **Supplementary Table 22. Results from Generalized Additive Mixed Models (GAMMs) for brain measures with interaction terms** | | | | | | | |
| --- | --- | --- | --- | --- | --- | --- | --- |
| **Outcome** | **Term** | **edf** | ***F*** | ***p*** | | ***p*_Bonferroni_** | |
| Whole brain volume | s(Gynecological Age) | 8.556 | 1243.862 | | **<.001** | | **<.001** |
|  | s(Menarche Timing) | 1 | 0.761 | | 0.383 | | 1 |
|  | s(General Exposome) | 4.753 | 130.586 | | **<.001** | | **<.001** |
|  | ti(Gynecological Age,General Exposome) | 4.829 | 24.208 | | **<.001** | | **<.001** |
|  | ti(Menarche Timing,General Exposome) | 1.43 | 0.962 | | 0.488 | | 1 |
|  | ti(Gynecological Age, Menarche Timing) | 8.11 | 46.938 | | **<.001** | | **<.001** |
|  |  |  |  | |  | | *R*^2^ = 0.133 |
| Subcortical volume | s(Gynecological Age) | 8.366 | 388.133 | | **<.001** | | **<.001** |
|  | s(Menarche Timing) | 1.792 | 6.175 | | **0.01** | | 0.48 |
|  | s(General Exposome) | 4.748 | 84.855 | | **<.001** | | **<.001** |
|  | ti(Gynecological Age,General Exposome) | 6.288 | 14.566 | | **<.001** | | **<.001** |
|  | ti(Menarche Timing,General Exposome) | 1.691 | 0.988 | | 0.491 | | 1 |
|  | ti(Gynecological Age, Menarche Timing) | 5.995 | 79.308 | | **<.001** | | **<.001** |
|  |  |  |  | |  | | *R*^2^ = 0.092 |
| Gray matter volume | s(Gynecological Age) | 8.412 | 5141.025 | | **<.001** | | **<.001** |
|  | s(Menarche Timing) | 1 | 50.563 | | **<.001** | | **<.001** |
|  | s(General Exposome) | 4.514 | 150.583 | | **<.001** | | **<.001** |
|  | ti(Gynecological Age,General Exposome) | 3.377 | 12.535 | | **<.001** | | **<.001** |
|  | ti(Menarche Timing,General Exposome) | 2.195 | 0.62 | | 0.5 | | 1 |
|  | ti(Gynecological Age, Menarche Timing) | 7.441 | 17.21 | | **<.001** | | **<.001** |
|  |  |  |  | |  | | *R*^2^ = 0.259 |
| Cortical thickness | s(Gynecological Age) | 7.183 | 3146.128 | | **<.001** | | **<.001** |
|  | s(Menarche Timing) | 1 | 173.129 | | **<.001** | | **<.001** |
|  | s(General Exposome) | 3.756 | 48.625 | | **<.001** | | **<.001** |
|  | ti(Gynecological Age,General Exposome) | 2.607 | 12.253 | | **<.001** | | **<.001** |
|  | ti(Menarche Timing,General Exposome) | 1.832 | 1.302 | | 0.352 | | 1 |
|  | ti(Gynecological Age, Menarche Timing) | 2.635 | 3.672 | | **0.009** | | 0.441 |
|  |  |  |  | |  | | *R*^2^ = 0.305 |
| Surface area | s(Gynecological Age) | 8.245 | 1152.496 | | **<.001** | | **<.001** |
|  | s(Menarche Timing) | 1 | 0.145 | | 0.704 | | 1 |
|  | s(General Exposome) | 4.458 | 102.164 | | **<.001** | | **<.001** |
|  | ti(Gynecological Age,General Exposome) | 3.579 | 8.831 | | **<.001** | | **0.02** |
|  | ti(Menarche Timing,General Exposome) | 1 | 0.503 | | 0.478 | | 1 |
|  | ti(Gynecological Age, Menarche Timing) | 6.31 | 27.943 | | **<.001** | | **<.001** |
|  |  |  |  | |  | | *R*^2^ = 0.105 |
| White matter volume | s(Gynecological Age) | 8.196 | 3266.474 | | **<.001** | | **<.001** |
|  | s(Menarche Timing) | 1 | 87.414 | | **<.001** | | **<.001** |
|  | s(General Exposome) | 4.413 | 70.126 | | **<.001** | | **<.001** |
|  | ti(Gynecological Age,General Exposome) | 7.06 | 15.281 | | **<.001** | | **<.001** |
|  | ti(Menarche Timing,General Exposome) | 1 | 0.03 | | 0.863 | | 1 |
|  | ti(Gynecological Age, Menarche Timing) | 4.552 | 98.667 | | **<.001** | | **<.001** |
|  |  |  |  | |  | | *R*^2^ = 0.115 |
| Lateral ventricles volume | s(Gynecological Age) | 7.24 | 741.89 | | **<.001** | | **<.001** |
|  | s(Menarche Timing) | 1.001 | 2.823 | | 0.093 | | 1 |
|  | s(General Exposome) | 1.001 | 8.48 | | **0.004** | | 0.172 |
|  | ti(Gynecological Age,General Exposome) | 2.432 | 3.272 | | **0.023** | | 1 |
|  | ti(Menarche Timing,General Exposome) | 1.787 | 0.818 | | 0.51 | | 1 |
|  | ti(Gynecological Age, Menarche Timing) | 1.73 | 16.501 | | **<.001** | | **<.001** |
|  |  |  |  | |  | | *R*^2^ = 0.018 |
| CSF | s(Gynecological Age) | 6.371 | 138.062 | | **<.001** | | **<.001** |
|  | s(Menarche Timing) | 1.511 | 3.399 | | 0.098 | | 1 |
|  | s(General Exposome) | 1 | 0.313 | | 0.576 | | 1 |
|  | ti(Gynecological Age,General Exposome) | 3.869 | 4.255 | | **0.002** | | 0.092 |
|  | ti(Menarche Timing,General Exposome) | 1.001 | 3.104 | | 0.078 | | 1 |
|  | ti(Gynecological Age, Menarche Timing) | 1 | 39.465 | | **<.001** | | **<.001** |
|  |  |  |  | |  | | *R*^2^ = 0.009 |
| Note. The term s() represents a smooth function of the predictor, modeling potential nonlinear fluctuations over time. The effective degrees of freedom (edf) indicate the complexity of the smooth term, with higher values suggesting greater flexibility. The term ti() represents the interaction between two predictors. *p*_Bonferroni_ = *p*-values after Bonferroni correction for multiple comparisons. Significant results are indicated in bold. CSF = cerebrospinal fluid. Total number of observations, *n* = 12,971. | | | | | | | |

| **Supplementary Table 23. Results from GAMMs for internalizing symptoms using the Child Behavior Checklist as outcome variable in other puberty markers based on self-report** | | | | | |
| --- | --- | --- | --- | --- | --- |
| **Puberty Marker** | **Term** | **edf** | ***F*** | ***p*** | ***p*_Bonferroni_** |
| Growth Spurt Onset | s(Puberty Marker Onset) | 5.594 | 65.88 | **<.001** | **<.001** |
|  | s(Menarche Timing) | 1 | 22.28 | **<.001** | **<.001** |
|  | s(General Exposome) | 4.455 | 8.937 | **<.001** | **<.001** |
|  |  |  |  |  | *R*^2^ = 0.013 |
| Breast Budding Onset | s(Puberty Marker Onset) | 6.998 | 58.82 | **<.001** | **<.001** |
|  | s(Menarche Timing) | 1 | 21.01 | **<.001** | **<.001** |
|  | s(General Exposome) | 4.673 | 8.804 | **<.001** | **<.001** |
|  |  |  |  |  | *R*^2^ = 0.014 |
| Body Hair Onset | s(Puberty Marker Onset) | 6.057 | 64.33 | **<.001** | **<.001** |
|  | s(Menarche Timing) | 1 | 18.5 | **<.001** | **<.001** |
|  | s(General Exposome) | 4.207 | 8.558 | **<.001** | **<.001** |
|  |  |  |  |  | *R*^2^ = 0.013 |
| Skin Change Onset | s(Puberty Marker Onset) | 7.12 | 63.88 | **<.001** | **<.001** |
|  | s(Menarche Timing) | 1 | 22.81 | **<.001** | **<.001** |
|  | s(General Exposome) | 4.42 | 7.969 | **<.001** | **<.001** |
|  |  |  |  |  | *R*^2^ = 0.015 |
| Note. The term s() represents a smooth function of the predictor, modeling potential nonlinear fluctuations over time. The effective degrees of freedom (edf) indicate the complexity of the smooth term, with higher values suggesting greater flexibility. *p*_Bonferroni_ = *p*-values after Bonferroni correction for multiple comparisons. Significant results are indicated in bold. Total number of observations range from *n* = 29,049–29,612. | | | | | |

| **Supplementary Table 24. Second derivative minima and maxima for internalizing symptoms using the Child Behavior Checklist in other puberty markers based on self-report** | | | |
| --- | --- | --- | --- |
| **Puberty Marker** | **Predictor** | **2^nd^ derivative minimum [CI]** | **2^nd^ derivative maximum [CI]** |
| Growth Spurt Onset | Puberty Marker Onset | 5.073 [3.028;5.214] | 1.794 [0.947;3.944] |
| Breast Budding Onset | Puberty Marker Onset | 5.123 [4.948;5.158] | 1.699 [-0.886;3.970] |
| Body Hair Onset | Puberty Marker Onset | 5.084 [3.952;5.267] | 1.979 [0.956;3.075] |
| Skin Change Onset | Puberty Marker Onset | 5.176 [4.993;5.249] | 1.918 [0.930;3.382] |
| Note. 2^nd^ derivatives describe the rate of change in the slope of the trajectory and identify inflection points where the curvature changes. The second derivative maximum indicates the point of fastest acceleration (steepest increase in slope), occurring where symptoms are increasing most rapidly. The second derivative minimum indicates the point of maximum deceleration (greatest slowing of the slope), occurring where the rate of increase or decrease is slowing most. For puberty markers, values represent time in years relative to individual onset (negative values = before onset, positive values = after onset). Credible intervals [CI] were derived from posterior sampling of the fitted generalized additive mixed models (GAMMs). | | | |

| **Supplementary Table 25. Results from Generalized Additive Mixed Models (GAMMs) for brain measures in other puberty markers based self-report** | | | | | | | |
| --- | --- | --- | --- | --- | --- | --- | --- |
| **Puberty Markers** | **Outcome** | **Term** | **edf** | ***F*** | | ***p*** | ***p*_Bonferroni_** |
| Growth Spurt | Whole brain volume | s(Puberty Marker Onset) | 8.539 | 921.446 | **<.001** | | **<.001** |
|  |  | s(Menarche Timing) | 1 | 4.447 | **0.035** | | 1 |
|  |  | s(General Exposome) | 4.701 | 130.248 | **<.001** | | **<.001** |
|  |  |  |  |  |  | | *R*^2^ = 0.123 |
|  | Subcortical volume | s(Puberty Marker Onset) | 8.515 | 163.062 | **<.001** | | **<.001** |
|  |  | s(Menarche Timing) | 2.687 | 1.491 | 0.132 | | 1 |
|  |  | s(General Exposome) | 4.592 | 85.581 | **<.001** | | **<.001** |
|  |  |  |  |  |  | | *R*^2^ = 0.09 |
|  | Gray matter volume | s(Puberty Marker Onset) | 8.216 | 6528.684 | **<.001** | | **<.001** |
|  |  | s(Menarche Timing) | 1 | 5.029 | **0.025** | | 1 |
|  |  | s(General Exposome) | 4.516 | 153.305 | **<.001** | | **<.001** |
|  |  |  |  |  |  | | *R*^2^ = 0.204 |
|  | Cortical thickness | s(Puberty Marker Onset) | 6.249 | 4229.431 | **<.001** | | **<.001** |
|  |  | s(Menarche Timing) | 1.825 | 3.381 | **0.023** | | 1 |
|  |  | s(General Exposome) | 3.79 | 55.773 | **<.001** | | **<.001** |
|  |  |  |  |  |  | | *R*^2^ = 0.241 |
|  | Surface area | s(Puberty Marker Onset) | 8.032 | 1166.097 | **<.001** | | **<.001** |
|  |  | s(Menarche Timing) | 1 | 3.149 | 0.076 | | 1 |
|  |  | s(General Exposome) | 4.388 | 102.005 | **<.001** | | **<.001** |
|  |  |  |  |  |  | | *R*^2^ = 0.093 |
|  | White matter volume | s(Puberty Marker Onset) | 8.39 | 3384.21 | **<.001** | | **<.001** |
|  |  | s(Menarche Timing) | 1 | 4.988 | **0.026** | | 1 |
|  |  | s(General Exposome) | 4.216 | 65.481 | **<.001** | | **<.001** |
|  |  |  |  |  |  | | *R*^2^ = 0.11 |
|  | Lateral ventricles volume | s(Puberty Marker Onset) | 5.824 | 940.585 | **<.001** | | **<.001** |
|  |  | s(Menarche Timing) | 1.687 | 1.357 | 0.169 | | 1 |
|  |  | s(General Exposome) | 1.003 | 6.623 | **0.01** | | 0.963 |
|  |  |  |  |  |  | | *R*^2^ = 0.016 |
|  | CSF | s(Puberty Marker Onset) | 5.672 | 129.858 | **<.001** | | **<.001** |
|  |  | s(Menarche Timing) | 1 | 17.283 | **<.001** | | **0.003** |
|  |  | s(General Exposome) | 1 | 0.007 | 0.931 | | 1 |
|  |  |  |  |  |  | | *R*^2^ = 0.01 |
| Breast Budding | Whole brain volume | s(Puberty Marker Onset) | 8.291 | 1139.931 | **<.001** | | **<.001** |
|  |  | s(Menarche Timing) | 1.012 | 3.131 | 0.077 | | 1 |
|  |  | s(General Exposome) | 4.643 | 133.728 | **<.001** | | **<.001** |
|  |  |  |  |  |  | | *R*^2^ = 0.129 |
|  | Subcortical volume | s(Puberty Marker Onset) | 8.187 | 262.78 | **<.001** | | **<.001** |
|  |  | s(Menarche Timing) | 2.712 | 1.666 | 0.107 | | 1 |
|  |  | s(General Exposome) | 4.587 | 86.906 | **<.001** | | **<.001** |
|  |  |  |  |  |  | | *R*^2^ = 0.089 |
|  | Gray matter volume | s(Puberty Marker Onset) | 7.95 | 7213.916 | **<.001** | | **<.001** |
|  |  | s(Menarche Timing) | 1 | 1.336 | 0.248 | | 1 |
|  |  | s(General Exposome) | 4.469 | 158.673 | **<.001** | | **<.001** |
|  |  |  |  |  |  | | *R*^2^ = 0.234 |
|  | Cortical thickness | s(Puberty Marker Onset) | 6.466 | 4194.963 | **<.001** | | **<.001** |
|  |  | s(Menarche Timing) | 2.182 | 1.733 | 0.178 | | 1 |
|  |  | s(General Exposome) | 4.035 | 52.777 | **<.001** | | **<.001** |
|  |  |  |  |  |  | | *R*^2^ = 0.272 |
|  | Surface area | s(Puberty Marker Onset) | 7.687 | 1372.856 | **<.001** | | **<.001** |
|  |  | s(Menarche Timing) | 1.011 | 2.395 | 0.122 | | 1 |
|  |  | s(General Exposome) | 4.277 | 106.053 | **<.001** | | **<.001** |
|  |  |  |  |  |  | | *R*^2^ = 0.099 |
|  | White matter volume | s(Puberty Marker Onset) | 8.15 | 3852.176 | **<.001** | | **<.001** |
|  |  | s(Menarche Timing) | 1.061 | 7.449 | **0.007** | | 0.631 |
|  |  | s(General Exposome) | 4.126 | 68.429 | **<.001** | | **<.001** |
|  |  |  |  |  |  | | *R*^2^ = 0.099 |
|  | Lateral ventricles volume | s(Gynecological Age) | 6.592 | 848.285 | **<.001** | | **<.001** |
|  |  | s(Menarche Timing) | 1.728 | 0.578 | 0.411 | | 1 |
|  |  | s(General Exposome) | 1.007 | 7.101 | **0.008** | | 0.73 |
|  |  |  |  |  |  | | *R*^2^ = 0.013 |
|  | CSF | s(Puberty Marker Onset) | 5.509 | 136.52 | **<.001** | | **<.001** |
|  |  | s(Menarche Timing) | 1 | 13.743 | **<.001** | | **0.02** |
|  |  | s(General Exposome) | 1 | 0.072 | 0.789 | | 1 |
|  |  |  |  |  |  | | *R*^2^ = 0.009 |
| Body Hair | Whole brain volume | s(Puberty Marker Onset) | 8.621 | 1006.538 | **<.001** | | **<.001** |
|  |  | s(Menarche Timing) | 1 | 3.188 | 0.074 | | 1 |
|  |  | s(General Exposome) | 4.691 | 128.609 | **<.001** | | **<.001** |
|  |  |  |  |  |  | | *R*^2^ = 0.126 |
|  | Subcortical volume | s(Puberty Marker Onset) | 8.55 | 214.437 | **<.001** | | **<.001** |
|  |  | s(Menarche Timing) | 2.631 | 1.581 | 0.122 | | 1 |
|  |  | s(General Exposome) | 4.606 | 83.345 | **<.001** | | **<.001** |
|  |  |  |  |  |  | | *R*^2^ = 0.087 |
|  | Gray matter volume | s(Puberty Marker Onset) | 8.41 | 6637.743 | **<.001** | | **<.001** |
|  |  | s(Menarche Timing) | 1 | 1.264 | 0.261 | | 1 |
|  |  | s(General Exposome) | 4.556 | 146.9 | **<.001** | | **<.001** |
|  |  |  |  |  |  | | *R*^2^ = 0.224 |
|  | Cortical thickness | s(Puberty Marker Onset) | 6.123 | 4363.767 | **<.001** | | **<.001** |
|  |  | s(Menarche Timing) | 1.653 | 0.582 | 0.386 | | 1 |
|  |  | s(General Exposome) | 3.822 | 49.328 | **<.001** | | **<.001** |
|  |  |  |  |  |  | | *R*^2^ = 0.249 |
|  | Surface area | s(Puberty Marker Onset) | 8.22 | 1187.654 | **<.001** | | **<.001** |
|  |  | s(Menarche Timing) | 1 | 2.021 | 0.155 | | 1 |
|  |  | s(General Exposome) | 4.379 | 100.332 | **<.001** | | **<.001** |
|  |  |  |  |  |  | | *R*^2^ = 0.098 |
|  | White matter volume | s(Puberty Marker Onset) | 8.364 | 3521.364 | **<.001** | | **<.001** |
|  |  | s(Menarche Timing) | 1.019 | 7.815 | **0.005** | | 0.498 |
|  |  | s(General Exposome) | 4.172 | 68.057 | **<.001** | | **<.001** |
|  |  |  |  |  |  | | *R*^2^ = 0.102 |
|  | Lateral ventricles volume | s(Puberty Marker Onset) | 6.05 | 916.103 | **<.001** | | **<.001** |
|  |  | s(Menarche Timing) | 1.444 | 0.561 | 0.396 | | 1 |
|  |  | s(General Exposome) | 1.002 | 7.048 | **0.008** | | 0.759 |
|  |  |  |  |  |  | | *R*^2^ = 0.016 |
|  | CSF | s(Puberty Marker Onset) | 5.612 | 138.687 | **<.001** | | **<.001** |
|  |  | s(Menarche Timing) | 1 | 13.693 | **<.001** | | **0.021** |
|  |  | s(General Exposome) | 1 | 0.195 | 0.659 | | 1 |
|  |  |  |  |  |  | | *R*^2^ = 0.011 |
| Skin Change | Whole brain volume | s(Puberty Marker Onset) | 8.722 | 949.031 | **<.001** | | **<.001** |
|  |  | s(Menarche Timing) | 1.05 | 2.724 | 0.103 | | 1 |
|  |  | s(General Exposome) | 4.637 | 134.345 | **<.001** | | **<.001** |
|  |  |  |  |  |  | | *R*^2^ = 0.122 |
|  | Subcortical volume | s(Puberty Marker Onset) | 8.707 | 184.194 | **<.001** | | **<.001** |
|  |  | s(Menarche Timing) | 2.686 | 1.575 | 0.121 | | 1 |
|  |  | s(General Exposome) | 4.578 | 82.941 | **<.001** | | **<.001** |
|  |  |  |  |  |  | | *R*^2^ = 0.086 |
|  | Gray matter volume | s(Puberty Marker Onset) | 8.392 | 6587.3 | **<.001** | | **<.001** |
|  |  | s(Menarche Timing) | 1 | 2.922 | 0.087 | | 1 |
|  |  | s(General Exposome) | 4.423 | 170.99 | **<.001** | | **<.001** |
|  |  |  |  |  |  | | *R*^2^ = 0.209 |
|  | Cortical thickness | s(Puberty Marker Onset) | 6.204 | 4305.843 | **<.001** | | **<.001** |
|  |  | s(Menarche Timing) | 1.383 | 2.751 | 0.054 | | 1 |
|  |  | s(General Exposome) | 3.977 | 67.06 | **<.001** | | **<.001** |
|  |  |  |  |  |  | | *R*^2^ = 0.245 |
|  | Surface area | s(Puberty Marker Onset) | 8.533 | 1137.709 | **<.001** | | **<.001** |
|  |  | s(Menarche Timing) | 1 | 1.979 | 0.159 | | 1 |
|  |  | s(General Exposome) | 4.322 | 106.798 | **<.001** | | **<.001** |
|  |  |  |  |  |  | | *R*^2^ = 0.093 |
|  | White matter volume | s(Puberty Marker Onset) | 8.698 | 3342.466 | **<.001** | | **<.001** |
|  |  | s(Menarche Timing) | 1 | 3.927 | **0.048** | | 1 |
|  |  | s(General Exposome) | 4.171 | 59.383 | **<.001** | | **<.001** |
|  |  |  |  |  |  | | *R*^2^ = 0.101 |
|  | Lateral ventricles volume | s(Puberty Marker Onset) | 5.718 | 995.962 | **<.001** | | **<.001** |
|  |  | s(Menarche Timing) | 1 | 1.033 | 0.309 | | 1 |
|  |  | s(General Exposome) | 1 | 4.775 | 0.029 | | 1 |
|  |  |  |  |  |  | | *R*^2^ = 0.014 |
|  | CSF | s(Puberty Marker Onset) | 5.404 | 139.438 | **<.001** | | **<.001** |
|  |  | s(Menarche Timing) | 1 | 14.885 | **<.001** | | **0.011** |
|  |  | s(General Exposome) | 1 | 0.002 | 0.965 | | 1 |
|  |  |  |  |  |  | | *R*^2^ = 0.009 |
| Note. The term s() represents a smooth function of the predictor, modeling potential nonlinear fluctuations over time. The effective degrees of freedom (edf) indicate the complexity of the smooth term, with higher values suggesting greater flexibility. *p*_Bonferroni_ = *p*-values after Bonferroni correction for multiple comparisons. Significant results are indicated in bold. CSF = cerebrospinal fluid. Total number of observations range from *n* = 12,614–12,868. | | | | | | | |

| **Supplementary Table 26. Second derivative minima and maxima for brain measures in other puberty markers**  **based on self-report** | | | | |
| --- | --- | --- | --- | --- |
| **Puberty Markers** | **Outcome** | **Predictor** | **2^nd^ derivative minimum [CI]** | **2^nd^ derivative maximum [CI]** |
| Growth Spurt | Whole brain volume | Puberty Marker Onset | 1.825 [1.745;1.865] | 3.063 [3.063;5.299] |
|  | Subcortical volume | Puberty Marker Onset | 1.785 [1.666;4.061] | 5.299 [3.063;5.339] |
|  | Gray matter volume | Puberty Marker Onset | 1.825 [1.745;1.905] | 3.103 [3.063;3.223] |
|  | Cortical thickness | Puberty Marker Onset | 1.825 [-2.087;5.179] | 3.223 [3.023;3.981] |
|  | Surface area | Puberty Marker Onset | 1.825 [1.745;1.905] | 3.103 [-0.890;5.299] |
|  | White matter volume | Puberty Marker Onset | 1.825 [1.705;4.061] | 5.259 [5.179;5.299] |
|  | Lateral ventricles volume | Puberty Marker Onset | 3.502 [1.905;4.101] | -0.890 [-1.928;5.339] |
|  | CSF | Puberty Marker Onset | 3.023 [1.985;3.861] | 4.979 [-1.968;5.179] |
| Breast Budding | Whole brain volume | Puberty Marker Onset | 1.829 [1.627;1.911] | 3.126 [-0.966;5.354] |
|  | Subcortical volume | Puberty Marker Onset | 1.829 [1.667;1.911] | 5.394 [-1.006;5.476] |
|  | Gray matter volume | Puberty Marker Onset | 1.789 [1.181;1.870] | 3.166 [-1.006;3.409] |
|  | Cortical thickness | Puberty Marker Onset | 1.100 [0.817;5.394] | 3.328 [3.045;4.017] |
|  | Surface area | Puberty Marker Onset | 1.829 [1.181;1.911] | 5.232 [-1.938;5.354] |
|  | White matter volume | Puberty Marker Onset | 1.870 [1.627;1.911] | 5.394 [-0.966;5.516] |
|  | Lateral ventricles volume | Puberty Marker Onset | 3.936 [-2.100;4.017] | 5.192 [-0.966;5.435] |
|  | CSF | Puberty Marker Onset | 3.288 [-2.060;4.017] | 5.151 [-1.006;5.435] |
| Body Hair | Whole brain volume | Puberty Marker Onset | 1.828 [1.787;1.870] | 3.041 [3.041;5.424] |
|  | Subcortical volume | Puberty Marker Onset | 1.870 [1.787;1.912] | 3.041 [3.041;5.508] |
|  | Gray matter volume | Puberty Marker Onset | 1.870 [1.787;1.912] | 3.083 [3.041;3.166] |
|  | Cortical thickness | Puberty Marker Onset | 1.912 [-1.935;2.037] | 3.375 [3.041;5.090] |
|  | Surface area | Puberty Marker Onset | 1.828 [1.745;1.912] | 3.041 [3.041;5.382] |
|  | White matter volume | Puberty Marker Onset | 1.787 [1.578;4.086] | 5.466 [3.041;5.592] |
|  | Lateral ventricles volume | Puberty Marker Onset | 3.710 [0.950;5.132] | -0.973 [-1.935;5.257] |
|  | CSF | Puberty Marker Onset | 3.877 [1.996;5.006] | -1.015 [-1.935;5.801] |
| Skin Change | Whole brain volume | Puberty Marker Onset | 1.854 [1.812;1.896] | 3.028 [2.986;5.460] |
|  | Subcortical volume | Puberty Marker Onset | 1.854 [1.854;4.118] | 5.460 [3.028;5.460] |
|  | Gray matter volume | Puberty Marker Onset | 1.854 [1.812;1.938] | 3.070 [3.028;3.154] |
|  | Cortical thickness | Puberty Marker Onset | 1.854 [-0.997;5.460] | 3.363 [3.028;4.076] |
|  | Surface area | Puberty Marker Onset | 1.854 [1.812;4.076] | 5.418 [3.028;5.460] |
|  | White matter volume | Puberty Marker Onset | 4.076 [1.854;4.118] | 5.418 [5.376;5.460] |
|  | Lateral ventricles volume | Puberty Marker Onset | 3.866 [1.980;4.160] | -0.830 [-2.171;5.795] |
|  | CSF | Puberty Marker Onset | 3.741 [1.938;4.579] | 5.418 [-2.129;5.753] |
| Note. 2^nd^ derivatives describe the rate of change in the slope of the trajectory and identify inflection points where the curvature changes. The second derivative maximum indicates the point of fastest acceleration (steepest increase in slope), occurring where measures are increasing most rapidly. The second derivative minimum indicates the point of maximum deceleration (greatest slowing of the slope), occurring where the rate of increase or decrease is slowing most. For puberty markers onset, values represent time in years relative to individual onset (negative values = before onset, positive values = after onset). Credible intervals [CI] were derived from posterior sampling of the fitted generalized additive mixed models (GAMMs). | | | | |
